# Genome-wide estimates of heritability and genetic correlations in Essential Tremor

**DOI:** 10.1101/555235

**Authors:** Monica Diez-Fairen, Sara Bandres-Ciga, Gabrielle Houle, Mike A. Nalls, Simon L. Girard, Patrick A. Dion, Cornelis Blauwendraat, Andrew B. Singleton, Guy A. Rouleau, Pau Pastor

## Abstract

Despite considerable efforts to identify disease-causing and risk factors contributing to essential tremor (ET), no comprehensive assessment of heritable risk has been performed to date. We use GREML-LDMS to estimate narrow-sense heritability due to additive effects (h^2^) and GREMLd to calculate non-additive heritability due to dominance variance (δ^2^) using data from 1,748 ET cases and 5,302 controls. We evaluate heritability per 10Mb segments across the genome and assess the impact of Parkinson’s disease (PD) misdiagnosis on heritability estimates. We apply genetic risk score (GRS) from PD and restless legs syndrome (RLS) to explore its contribution to ET risk and further assess genetic correlations with 832 traits by Linkage disequilibrium score regression. Our results show for the first time that ET is a highly heritable condition (h^2^=0.755, s.e=0.075) in which additive common variability plays a prominent role. In contrast, dominance variance shows insignificant effect on the overall estimates. Heritability split by 10Mb regions revealed increased estimates at chromosomes 6 and 21 suggesting that these may contain causative risk variants influencing susceptibility to ET. The proportion of genetic variance due to PD misdiagnosed cases was estimated to be 5.33%. PD and RLS GRS were not significantly predictive of ET case-control status demonstrating that despite overlapping symptomatology, ET does not seem to share genetic etiologies with PD or RLS. Our study suggests that most of ET genetic component is yet to be discovered and future GWAS will reveal additional risk factors that will improve our understanding of this disabling disorder.

## INTRODUCTION

Essential tremor (ET) is one of the most prevalent neurological disorder and the most common adult-onset movement disorder after restless legs syndrome (RLS) (1). Its prevalence increases markedly with age, being 0.9% in the general population and 4.6% in individuals older than 65 years (2). ET is defined by the presence of action (postural and kinetic) tremor (3). We suggest that the lack of disease-specific diagnostic markers for ET, clinical overlap with other neurological disorders such as Parkinson’s disease (PD) and dystonia, and coexistence with other neurological diseases such as RLS (4), have made the identification of the underlying causes of ET difficult. In fact, several studies have reported misdiagnosis rates of ET ranging from 37% (5) to 50% (6), mainly with PD.

Both genetic and environmental factors are likely to contribute to ET etiology. Twin studies have shown a greater concordance in monozygotic twins (60-93%) than in dizygotic twins (27-29%), suggesting a role of genetic factors in disease pathogenesis (7,8). Twin studies estimated the heritable component of essential tremor between 45% and 90% (9). However, the lack of complete disease concordance in monozygotic twins, the existence of sporadic cases, and the variability in age at onset, location and severity of tremor within families, support the hypothesis that ET is genetically complex and that environmental factors may also contribute to the etiology of ET (10–12).

The inheritance pattern of ET remains unclear. Several family-based studies have reported an autosomal dominant pattern of inheritance (13–17). However, the estimated 23% frequency of affected first-degree relatives is much lower than the 50% and 25% expected for an autosomal dominant inheritance with complete penetrance and autosomal recessive inheritance, respectively (18,19). Hence, the ET genetic architecture is clearly complex and it is likely that both monogenic and multifactorial inheritance contribute to the genetic etiology of ET.

Despite evidence suggesting a strong genetic component in the etiology of ET, only a few susceptibility loci have been identified. Family-based genome-wide linkage studies were the first to map three susceptibility loci for familial ET (14,15,20), although no causal mutations have been clearly identified within these loci. More recently, genome-wide association studies (GWAS) have identified five loci to be associated with risk for sporadic ET (21–23). Whole-exome sequencing (WES) has also identified putative, rare candidate variants for ET risk (24,25); further replication in larger cohorts remains essential to confirm these associations.

Despite the success of GWAS at identifying common genetic variability associated with human complex diseases, the associated variants only explain a small fraction of the heritability for most traits (26,27). Heritability is defined as the proportion of phenotypic variance in a population explained by the genetic variation between individuals. Genetic sources of variation can be divided into three subcategories: additive variance, dominance variance and epistatic variance. Additive genetic variance (commonly known as narrow-sense heritability or h^2^) relates to an allele’s independent effect on a phenotype; dominance variance (δ^2^) refers to the effect on a phenotype caused by interactions between alternative alleles at a specific locus and epistatic variance refers to the interaction between different alleles in different loci (28). Genomic-relationship-matrix restricted maximum likelihood (GREML-LDMS) has recently emerged as an unbiased tool to estimate the phenotypic variance explained by genetics, considering all the genome-wide single nucleotide polymorphisms (SNPs) simultaneously and regardless of the minor allele frequency (MAF) and linkage disequilibrium (LD) properties of variants (29).

Here, we aim to investigate the genetic architecture of ET by providing heritable estimates of risk. We estimate the narrow-sense heritability of ET using data from the largest ET GWAS to date (23). We further attempt to study the contribution of non-additive variance to the total heritability and evaluate heritability per 10Mb segments across the genome to nominate regions of interest. We assess whether the estimated ET heritability could be explained by the presence of misdiagnosed cases by calculating the ET heritability due to PD misdiagnosis (30). In addition, we examine the genetic risk scores (GRS) from PD and RLS to explore the genetic influence of these diseases to the risk of ET. Finally, we use LD score regression to investigate genetic correlations and overlapping etiologies across 832 traits and diseases.

## RESULTS

### Heritability estimates

We applied GREML-LDMS to 1,751 cases and 5,311 control individuals of European ancestry to estimate the proportion of phenotypic variance associated with ET. We estimated the narrow-sense heritability of ET to be h^2^ = 0.755, s.e = ± 0.075, p <2 x10^−16^. To determine if non-additive variance explains to some extent the total disease heritability, we calculated the disease dominance variance as implemented in the tool GCTA-GREMLd (31). Our results suggest that ET does not show significant dominance variance with an overall estimate δ2=−0.003, s.e. = 0.0008.

We further estimated heritability split by 10 Mb intervals after adjusting by sex, age, principal components (PCs), ET prevalence and the number of variants present in each of the 294 segments across the genome (**Figure 1**, **Supplementary Table 1**).

**Figure 1.**
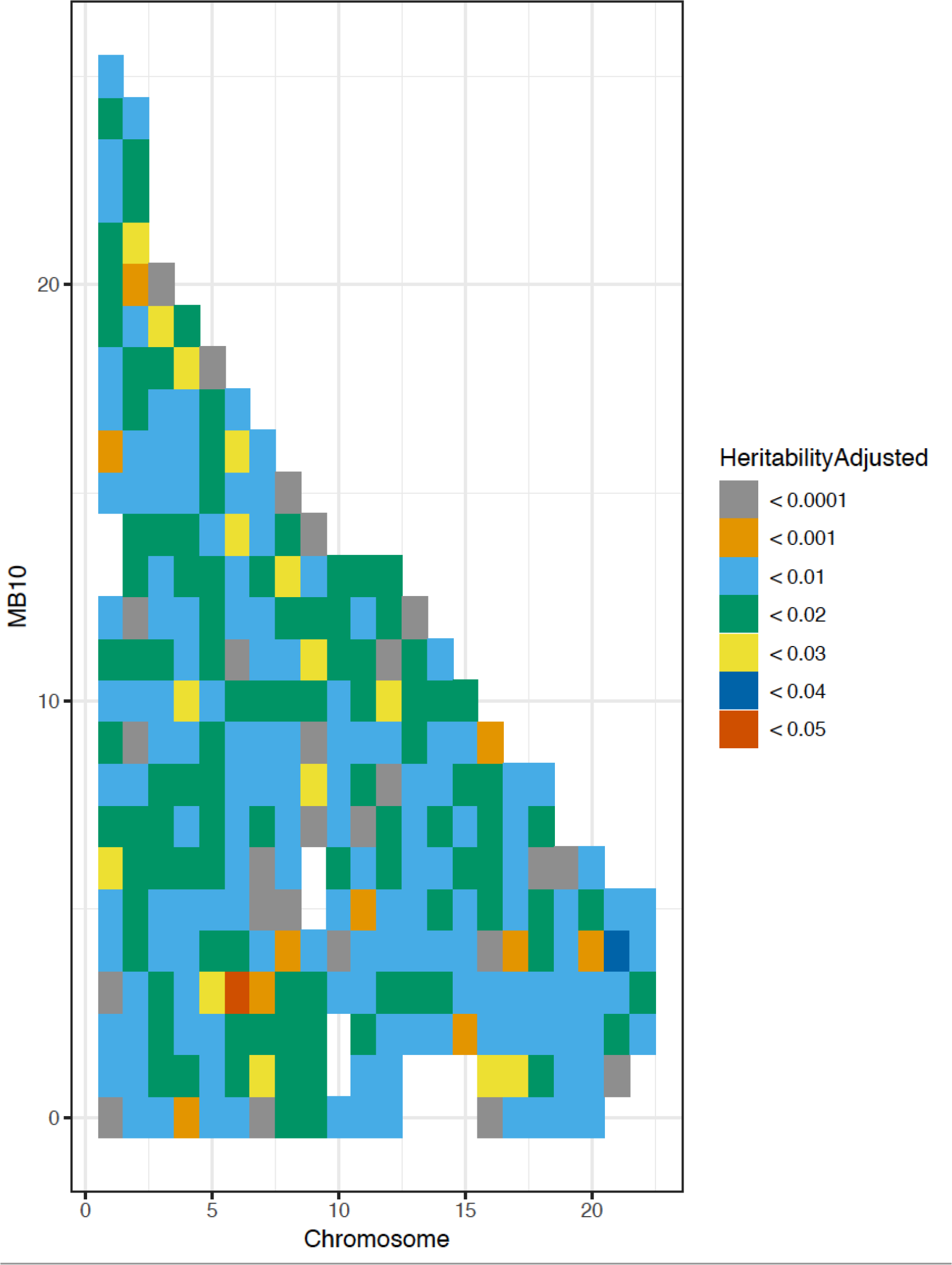
Genome-wide heritability estimates in 10 Mb imputed segments. Heat plots show heritability estimates across the genome. The x-axis represents chromosomes 1 to 22, and the y-axis represents the length of each 10 Mb region in base pairs. The segments with higher estimated heritability are denoted by dark blue (chr 21) and dark red (chr 6). Of note, this plot displays adjusted heritability estimates by number of variants per segment.

Since ET heritability may be driven in part by misdiagnosed PD cases, we estimated how much heritability could be expected due to a proportion of our clinically diagnosed cases receiving a misdiagnosis. Based on Jain and colleagues, we considered that approximately 15% of our cases were false positives (true PD cases) and in 22% the clinical picture was due to other neurological conditions (5). PD comprises the majority of these false positives since it has a higher disease prevalence compared to other movement disorders mimicking ET such as dystonia (32). The estimated heritability of PD is 20.9% (33) and of other diseases is 10% (30). We gave a conservatively low heritability of 10% to false positives of diseases other than PD as previously reported (30). Assuming that all individuals have been clinically diagnosed, we calculated an expected heritability due to PD misdiagnosis of 5.33% (see **Supplementary materials**).

### Genetic risk profiling

We used genetic risk profiling to estimate the extent to which the risk of developing ET is attributable to underlying PD and RLS risk. We demonstrate that individuals with the highest burden of PD genetic risk (top quartile of the analysis) were not more likely to develop ET than individuals in the lowest quartile of the analysis (OR=1.18, 95% CI=0.97-1.45, Z-score p-value=0.085). Similarly, the increase in ET risk associated with RLS was not statistically significant (OR=0.98, 95% CI=0.91-1.05, Z-score p-value=0.59). After removing the expected 15% ET cases (262) carrying the highest PD genetic burden among all ET cases, ET heritability estimates increased (h^2^ =0.829, s.e = ±0.051, p<2 × 10^−16^).

### Genetic correlations across multiple traits

LD score regression was applied to examine the genetic correlation between the recently published GWAS of ET (23) and 832 available phenotypes. We did not observe any genetic correlation significantly associated with ET after Bonferroni correction (832 tests; p-value <0.00006) which could be likely due to lack of statistical power (**Supplementary Table 2**).

## DISCUSSION

Here we present a comprehensive study of the genetic basis of ET. We provide estimates of narrow-sense heritability for ET and heritability due to PD misdiagnosis, quantify the variance explained by PD and RLS genetic risk and investigate genetic correlations between ET and over 832 traits and diseases.

Heritability estimates using GCTA have been recently performed in other neurodegenerative diseases reporting values of 20.9% for PD, 21% for amyotrophic lateral sclerosis (ALS) and 2.09-6.65% for multiple system atrophy (MSA) (31,33,34). Since it is known that GCTA heritability estimates could be biased if causal variants have different MAF spectrum that the variants used in the analyses or are enriched in genomic regions with higher or lower LD than average; we used the GREML-LDMS method to obtain unbiased estimates of heritability as implemented before (29,35). Our study shows that phenotypic variance for ET is roughly 75% under the additive model, approaching to that reported in twin studies (9).

Multi-component models such as GREML-LDMS are thought to provide the most robust estimates, although they are more computationally intensive, have higher standard errors than single-component models, and require larger datasets to achieve reliable estimates (36). However, even when using multi-component approaches, heritability is likely underestimated. We believe that these estimates might still improve as sample sizes increase and larger imputation panels are used. In addition to the additive genetic component of a certain trait, non-additive factors including dominance variance and epistatic variance are thought to contribute considerably to a trait’s total heritable component. In concordance with recent findings from other reported traits (35), our results do not support the notion that dominance variance has a substantial effect on the genetic heritability of ET.

Many studies have failed to consistently unravel and replicate strong GWAS associated signals in ET (37–41), which suggests that ET is genetically very complex in nature and as other diseases, might not be driven by a single locus but rather by many loci with very small effect sizes that cumulatively increase disease risk. In that way, we provided data showing the contribution of 294 intervals of 10Mb size across the genome to the overall heritability. We hypothesize that chromosomes 6 and 21 contribute most to ET heritability and may contain causative risk variants that influence susceptibility to ET and therefore warrant further study. Interestingly, loci within chromosome 6 have been previously associated with ET (20,42). Since it remains difficult to definitively identify which genes within these regions are responsible of the genetic risk in the current data, these regions are good candidates for future functional studies.

Due to the fact that ET has a heterogeneous clinical presentation and shows overlapping phenotypic features with other neurodegenerative disorders, mostly PD and dystonia, misdiagnosis is relatively common (5,6). Thus, we could expect that ET may have a similar heritability estimate as these diseases. However, our ET heritability estimates are markedly higher than those previously reported for PD and MSA (30,33,43). In fact, after excluding those individuals with the highest burden of PD risk, our heritability estimates increase significantly, demonstrating that PD misdiagnosis reduces heritability estimates in ET.

We used cumulative GRS to estimate how much of the phenotypic variance in our ET cohort is due to PD and RLS known genetic risk factors. PD GRS was not significantly predictive of case-control status for ET after adjusting for misdiagnosis, demonstrating that individuals with the highest burden of PD genetic risk are not more likely to develop ET than individuals with the lowest burden. No significant association was observed for RLS. Despite overlapping symptomatology, our findings do not support the existence of shared genetic etiologies between PD and ET although further research is needed to replicate these findings.

However, our study has several limitations. First, we have to assume that heritability estimates provided by GREML-LDMS represent a lower limit of heritability since rare and structural variants are not fully covered by current genotyping technologies or imputation. Second, despite being the largest to date, our ET cohort is relatively small compared to other recently published GWAS performed in other diseases. Therefore, we believe our analyses were underpowered to detect any significant pleiotropic effect across 832 phenotypes after multiple testing correction. Larger GWAS will be needed to shed some light on the complex genetic architecture of ET, including the identification of new causal variants that will consequently improve our understanding of the ET heritability as larger genetic contribution to disease risk is known. Third, the ET dataset used only comprises individuals from European and North American ancestry; thus, studying populations of different descent may identify novel risk loci and novel genetic correlations which could improve our understanding of ET etiology.

In conclusion, ET is a highly heritable disease in which common genetic variability appears to play a prominent role. We showed that not only a large proportion of ET heritability can be ascribed to additive common genetic variation but also that there are yet-to-be discovered loci contributing to the etiology of ET. Similarly to other genetic complex diseases, our data provide strong evidence that many of the disease-associated loci will have a small contribution to the phenotypic variation and thus, very large sample sizes will be required for their individual effects to become statistically significant.

## MATERIALS AND METHODS

### Sample description

We used data from the discovery stage of the most recent published GWAS of ET (23), comprised of 1,751 ET cases and 5,311 healthy controls of primarily North-Western European descent. Study descriptives for the GWAS dataset are summarized in **Table 1**.

ET dataset (Table 1. Demographic data of the Muller et al., 2016)

| Discovery stage |  |  |  |  |  |
| --- | --- | --- | --- | --- | --- |
|  | Side | N | Sex (%Female) | Age (Mean±SD; N=missing) | Age of Onset (Mean±SD; N=missing) |
| **CASES** |  |  |  |  |  |
|  | Europe | 887 | 37.1 | 62.34(±14.64; N=25) | 39.41(±21.95;N=94) |
|  | North America | 891 | 53.2 | 69.88(±14.45; N=590) | 50.20(±22.73; N=299) |
|  | **Total** | **1,778** | **45.2** | **64.29(±14.96; N=615)** | **44.02(±22.91; N=393)** |
| **CONTROLS** |  |  |  |  |  |
|  | Europe | 4,603 | 49.8 | 50.37(±14.13; N=8) | --- |
|  | North America | 773 | 56.0 | NA (N=773) | --- |
|  | **Total** | **5,376** | **50.7** | **50.37(±14.13; N=781)** | **---** |
| Replication stage |  |  |  |  |  |
|  | Side | N | Sex (%Female) | Age (Mean±SD; N=missing) | Age of Onset (Mean±SD; N=missing) |
| **CASES** |  |  |  |  |  |
|  | Europe | 425 | 49.6 | 63.53(±15.19; N=0) | 42.00(±22.31; N=22) |
|  | North America | 604 | 54.4 | 69.06(±14.70; N=62) | 49.63(±21.16; N=90) |
|  | **Total** | **1,029** | **52.3** | **66.94(±15.12; N=62)** | **46.28(±21.99; N=112)** |
| **CONTROLS** |  |  |  |  |  |
|  | Europe | 431 | 44.3 | 42.24(±17.22; N=0) | --- |
|  | North America | 634 | 67.7 | 71.49(±12.53; N=368) | --- |
|  | **Total** | **1,065** | **53.2** | **52.95(±21.07; N=368)** | **---** |

Abbreviations: ET = essential tremor; N = number; SD = standard deviation.

### Quality control analyses and imputation

Genotyping is described in detail elsewhere (23). Briefly, samples were genotyped using Axiom Genome-Wide CEU 1 Array Plate by Affymetrix and genotypes were called using the Affymetrix Power Tools (v1.15.1). Quality control analysis was performed as follows; Samples with call rates of less than 95% and whose genetically determined sex from X chromosome heterogeneity did not match that from clinical data were excluded from the analysis. Samples exhibiting excess heterozygosity estimated by an F statistic > +/− 0.25 were also excluded. Once preliminary sample-level QC was completed, SNPs with MAF < 0.01, Hardy-Weinberg Equilibrium (HWE) p-value < 1E-5 and missingness rates > 5% were excluded. Genotyped SNPs thought to be in LD in a sliding window of 50 adjacent SNPs which scrolled through the genome at a rate of 5 overlapping SNPs were also removed from the following analyses (as were palindromic SNPs). Next, samples were clustered using principal component analysis (PCA) to evaluate European ancestry as compared to the HapMap3 CEU/TSI populations (International HapMap Consortium, 2003) **(Supplementary Figure 1)**. Confirmed European ancestry samples were extracted and PCs 1-20 were used as covariates in all analysis. Samples related at the level of cousins or closer (sharing proportionally more than 12.5% of alleles) were dropped from the following analysis. After quality control analyses, the total number of SNPs numbered 349,321. Then, the ET dataset was imputed using the Haplotype Reference Consortium under default settings with phasing using the EAGLE option (44,45). Imputed variants numbered 8,957,963 after removing SNPs with MAF < 0.1% and poor imputation quality (R^2^ < 0.30).

### Heritability estimates

For a given disease trait, Genome-wide Complex Trait Analysis (GCTA) estimates the heritability explained by all genome-wide SNPs simultaneously (27,46,47). Therefore, variants with a smaller effect that would not pass the significance threshold in GWAS analysis are included in the analysis, providing a more robust measurement of the heritability. In GCTA total genetic variation is calculated by generating a genetic relationship matrix (GRM) that estimates the genetic relationship between pairs of individuals (46). The relative similarity across cases compared to that between cases and controls is used to estimate total heritability of the disease phenotype. GRM is then used as input for the restricted maximum likelihood analysis (REML) to estimate the SNP-based heritability (46).

Recently, an alternative method called GREML-LDMS has been developed to avoid the bias that occurs in GCTA SNP-based estimates due to MAF and LD properties of variants (29). Segment-based LD scores are calculated using a segment length of 200Kb (with 100Kb overlap between two adjacent segments), these are then used to stratify the SNPs into quartiles. Then, a GRM is computed for each sample using the stratified SNPs. Lastly, REML analysis is performed using the multiple GRMs.

We used GREML-LDMS to estimate ET heritability based on imputed data. All analyses were adjusted for 20 PCs, sex, age and prevalence (46,48). Heritability estimates were transformed to the liability scale using the disease prevalence to account for ascertainment bias (34,43,46). The disease prevalence was estimated at 0.009 as previously reported (2).

We estimated the narrow-sense heritability of ET considering all imputed SNPs simultaneously (34,43). Then, we used genome-wide imputed SNPs as implemented in GCTA-GREMLd to estimate the dominance GRM between pairs of individuals (31). This method calculates the additive and dominance GRMs and fits both GRMs in a mixed linear model to estimate additive and dominance variance simultaneously. Additionally, we localized heritability signals within the genome by applying a mixed model heritability analysis to unmerged segments of the genome in 10Mb increments.

Finally, based on a comprehensive literature overview of misdiagnosis rates, we considered the estimated ET misdiagnosis rate as 37% (false positive rate), where 41.5% of false positives (15% of cases) were misdiagnosed as PD and 58.5% (22% of cases) as other conditions (5), and heritability estimates of 20.9% for PD (33) and 10% for all other disorders. We calculated the proportion of ET heritability due to PD misdiagnosis as described elsewhere (30; **Supplementary materials**).

### Genetic risk profiling

In order to explore the genetic influence of PD and RLS on the risk of ET, cumulative genetic risk scores (GRS) were calculated in the ET dataset as described elsewhere (49). Genetic risk profiling was estimated incorporating SNPs either associated with PD (33) or with RLS (50) in both the most recent and largest GWAS meta-analyses to date. Risk allele dosages were summed and a GRS was generated across loci per sample. All the SNPs were weighted by their log odds ratios, giving greater weight to alleles with higher risk estimates. Then, the dataset was divided into quintiles based on the GRS. Logistic regression was performed regressing disease status (PD or RLS) against quintile membership and odds ratios were reported comparing the reference group (lowest risk quintile) to the remaining 4 quintiles. GRS was adjusted by 20 PCs, age and sex.

### Genetic correlations across multiple traits

We performed LD score regression analyses to study shared genetic risk with ET. This method is based on the premises that SNPs in regions of high LD tag a greater proportion of the genome and will show stronger associations than SNPs in regions of low LD. Using the known LD structure of a reference SNP panel, reliable estimates of overlapping inheritance can be estimated. We used *LD Hub* (http://ldsc.broadinstitute.org/ldhub/), a centralized database of summary-level GWAS results across 832 diseases/traits gathered from publicly available resources (51). Default settings were used in the analyses and final results were adjusted for multiple testing by using Bonferroni correction **(Supplementary materials)**.

## ACKNOWLEDGMENTS

This work was supported by the Spanish Ministry of Science and Innovation to PP [SAF2013-47939-R (2013-2018)]. This work was supported in part by the Intramural Research Programs of the National Institute of Neurological Disorders and Stroke (NINDS), the National Institute on Aging (NIA), and the National Institute of Environmental Health Sciences both part of the National Institutes of Health, Department of Health and Human Services [project numbers 1ZIA-NS003154, Z01-AG000949-02 and Z01-ES101986].

## CONFLICT OF INTEREST STATEMENT

Mike A. Nalls’ participation is supported by a consulting contract between Data Tecnica International and the National Institute on Aging, NIH, Bethesda, MD, USA, as a possible conflict of interest Dr. Nalls also consults for Neuron23 Inc., Lysosomal Therapeutics Inc., Illumina Inc., the Michael J. Fox Foundation and Vivid Genomics among others. All other authors declare that they have no conflicts of interest.

