## Supplementary Materials for "Genome-wide estimates of heritability and genetic correlations in Essential Tremor"

### MATERIALS AND METHODS

#### Heritability estimates due to PD misdiagnosis

Because ET is a common disorder and has a variable clinical presentation, other less common disorders can be mistaken for ET. We estimated how much heritability could be expected due to a proportion of clinically diagnosed cases receiving a misdiagnosis.

Here, we used rates of false clinical diagnosis from Jain et al., 2006. We consider ET misdiagnosis rate as 37%, where 15% of false positives were diagnosed as PD (15% of 37% = 40.54%) and 22% as other conditions (22% of 37%= 59.46%), and heritability estimates of 20.9% for PD (Nalls et al., 2018) and 10% for all other disorders.

Based on these rates, we calculated heritability due to misdiagnosis following the formula proposed by Federoff *et al* (Federoff et al., 2016):

Misdiagnosed cases (M) = Clinical cases (Clinical diagnosed cases/Total) * False Positive rate (FPR)

M = ET cases misdiagnosed as PD (P) + ET cases misdiagnosed as others (O)

P = 0.4054M

O= 0.5946M

ET heritability due to misdiagnosis ($H_m$) = PD heritability * P + other disorders heritability * O

$H_m$ = (0.209 * 0.4054M) + (0.1 * 0.5946M) = (0.0847286M) + (0.05946M) = 0.1441886M

Which simplifies to:

$H_m$ = 0.1441886 * (FPR*(Clin/Total))

$H_m$ = 0.1441886 * (0.37*(1748/1748) = 0.053349782 > 5.3% of heritability is due to PD misdiagnosis rate assuming that all individuals have been clinically diagnosed

**Genetic correlations across multiple traits (LD score regression)**

Briefly, individual genotyping data was first imputed using the Sanger Imputation Services to impute the data against the UK10K + 1000 Genomes Phase 3 panel. Hapmap phase 3 markers with a R2>0.9 were extracted and used to perform association testing between the phenotype and SNPs using PLINK following an additive logistic regression adjusted for sex and the first 10 PCs. Outputted summary statistics were used as an input to LD-Hub.

### RESULTS

**Supplementary Figure 1. Essential tremor cohort with HapMap3 populations; Essential tremor cohort ancestry; Essential Tremor cohort with European CEU/TSI/MEX populations**

**Supplementary Table 1. Comparison of adjusted versus unadjusted genome-wide heritability estimates split by 10 Mb imputed segments**

**Supplementary Table 2. Linkage disequilibrium score regression analyses performed on 832 GWAS**
