## Supplementary Table 1 for "Genome-wide estimates of heritability and genetic correlations in Essential Tremor"

**Table S1. Comparison of adjusted versus unadjusted genome-wide heritability estimates split by 10 Mb imputed segments**

| Chromosome | MB10 | h² | SE | NSNP | Adjusted h² |
|---|---|---|---|---|---|
| 6 | 3 | 0,042677 | 0,00915 | 73813 | 0,042677 |
| 21 | 4 | 0,03094 | 0,00781 | 31976 | 0,03094 |
| 9 | 11 | 0,029784 | 0,007754 | 47542 | 0,029784 |
| 17 | 1 | 0,02953 | 0,00805 | 30088 | 0,02953 |
| 3 | 19 | 0,026268 | 0,007515 | 34546 | 0,026268 |
| 9 | 8 | 0,024481 | 0,007185 | 39366 | 0,024481 |
| 16 | 1 | 0,023813 | 0,007586 | 47000 | 0,023813 |
| 5 | 3 | 0,023216 | 0,006742 | 39772 | 0,023216 |
| 12 | 10 | 0,023102 | 0,007221 | 42646 | 0,023102 |
| 1 | 6 | 0,022721 | 0,007208 | 35738 | 0,022721 |
| 8 | 13 | 0,022285 | 0,007528 | 39967 | 0,022285 |
| 2 | 21 | 0,021904 | 0,007004 | 35843 | 0,021904 |
| 4 | 10 | 0,021546 | 0,006759 | 41989 | 0,021546 |
| 7 | 1 | 0,02131 | 0,007306 | 48380 | 0,02131 |
| 6 | 14 | 0,021124 | 0,006832 | 33405 | 0,021124 |
| 6 | 16 | 0,020309 | 0,00723 | 41731 | 0,020309 |
| 4 | 18 | 0,020167 | 0,007531 | 43028 | 0,020167 |
| 14 | 7 | 0,019827 | 0,006915 | 36259 | 0,019827 |
| 3 | 6 | 0,019768 | 0,007607 | 45520 | 0,019768 |
| 15 | 10 | 0,019708 | 0,006665 | 25714 | 0,019708 |
| 8 | 1 | 0,01946 | 0,007707 | 60291 | 0,01946 |
| 14 | 3 | 0,019067 | 0,006509 | 33424 | 0,019067 |
| 13 | 11 | 0,018834 | 0,006873 | 31549 | 0,018834 |
| 3 | 3 | 0,018723 | 0,006738 | 39549 | 0,018723 |
| 9 | 1 | 0,018434 | 0,007436 | 50834 | 0,018434 |
| 18 | 1 | 0,018409 | 0,006801 | 30151 | 0,018409 |
| 8 | 14 | 0,01824 | 0,006155 | 26091 | 0,01824 |
| 6 | 4 | 0,017912 | 0,006627 | 30113 | 0,017912 |
| 14 | 5 | 0,017877 | 0,00638 | 38380 | 0,017877 |
| 5 | 12 | 0,017755 | 0,006965 | 51443 | 0,017755 |
| 16 | 8 | 0,017695 | 0,008093 | 58820 | 0,017695 |
| 5 | 16 | 0,017501 | 0,006481 | 41992 | 0,017501 |
| 18 | 5 | 0,017216 | 0,00616 | 36272 | 0,017216 |
| 3 | 2 | 0,017125 | 0,006906 | 45709 | 0,017125 |
| 1 | 24 | 0,01656 | 0,006506 | 31680 | 0,01656 |
| 10 | 12 | 0,016522 | 0,006144 | 31198 | 0,016522 |
| 4 | 6 | 0,016489 | 0,006098 | 42053 | 0,016489 |
| 13 | 10 | 0,016432 | 0,006301 | 39301 | 0,016432 |
| 3 | 1 | 0,016357 | 0,006808 | 35758 | 0,016357 |
| 7 | 10 | 0,015647 | 0,005948 | 27281 | 0,015647 |
| 7 | 2 | 0,015568 | 0,007262 | 51558 | 0,015568 |
| 11 | 10 | 0,01552 | 0,006447 | 49033 | 0,01552 |
| 2 | 18 | 0,015201 | 0,00641 | 41012 | 0,015201 |
| 3 | 14 | 0,015074 | 0,005744 | 29745 | 0,015074 |
| 2 | 22 | 0,015045 | 0,006116 | 29199 | 0,015045 |
| 6 | 10 | 0,015012 | 0,005762 | 36772 | 0,015012 |
| 5 | 4 | 0,015011 | 0,005767 | 37852 | 0,015011 |
| 12 | 13 | 0,014986 | 0,006149 | 23145 | 0,014986 |
| 2 | 17 | 0,014985 | 0,006463 | 37788 | 0,014985 |
| 11 | 13 | 0,014936 | 0,006777 | 37433 | 0,014936 |
| 3 | 8 | 0,014864 | 0,005494 | 37686 | 0,014864 |
| 9 | 0 | 0,014864 | 0,006167 | 24164 | 0,014864 |
| 2 | 4 | 0,014777 | 0,006262 | 44884 | 0,014777 |

| | | | | | |
|---|---|---|---|---|---|
| 2 | 6 | 0,014708 | 0,006006 | 41378 | 0,014708 |
| 2 | 11 | 0,014674 | 0,005891 | 28431 | 0,014674 |
| 9 | 3 | 0,014637 | 0,005937 | 45885 | 0,014637 |
| 8 | 0 | 0,014635 | 0,006962 | 43098 | 0,014635 |
| 13 | 9 | 0,017593 | 0,00596 | 40540 | 0,0144535 |
| 18 | 4 | 0,011314 | 0,005684 | 40540 | 0,0144535 |
| 9 | 10 | 0,014379 | 0,005869 | 33264 | 0,014379 |
| 13 | 3 | 0,014375 | 0,005956 | 34460 | 0,014375 |
| 15 | 8 | 0,01428 | 0,005987 | 30066 | 0,01428 |
| 10 | 13 | 0,014212 | 0,006584 | 34116 | 0,014212 |
| 10 | 6 | 0,01418 | 0,005874 | 42613 | 0,01418 |
| 6 | 1 | 0,014008 | 0,006073 | 36306 | 0,014008 |
| 2 | 5 | 0,01393 | 0,006811 | 54561 | 0,01393 |
| 1 | 20 | 0,013771 | 0,005348 | 30602 | 0,013771 |
| 2 | 14 | 0,013767 | 0,005771 | 35016 | 0,013767 |
| 4 | 19 | 0,0137 | 0,006086 | 24687 | 0,0137 |
| 3 | 7 | 0,013347 | 0,00644 | 38798 | 0,013347 |
| 11 | 8 | 0,013314 | 0,005968 | 41290 | 0,013314 |
| 5 | 8 | 0,013127 | 0,006182 | 40176 | 0,013127 |
| 8 | 3 | 0,013059 | 0,006049 | 37300 | 0,013059 |
| 8 | 10 | 0,012945 | 0,005816 | 33153 | 0,012945 |
| 6 | 2 | 0,012934 | 0,006924 | 46430 | 0,012934 |
| 12 | 6 | 0,012858 | 0,005312 | 32173 | 0,012858 |
| 11 | 2 | 0,012703 | 0,005761 | 34724 | 0,012703 |
| 5 | 17 | 0,012602 | 0,00664 | 34005 | 0,012602 |
| 22 | 3 | 0,012539 | 0,006175 | 35700 | 0,012539 |
| 4 | 13 | 0,012534 | 0,005909 | 38918 | 0,012534 |
| 5 | 6 | 0,012527 | 0,0057 | 39477 | 0,012527 |
| 14 | 10 | 0,012325 | 0,005579 | 25895 | 0,012325 |
| 12 | 12 | 0,012281 | 0,005756 | 28840 | 0,012281 |
| 11 | 11 | 0,012273 | 0,005427 | 33377 | 0,012273 |
| 4 | 1 | 0,012253 | 0,005264 | 33422 | 0,012253 |
| 9 | 2 | 0,01222 | 0,006384 | 44403 | 0,01222 |
| 21 | 2 | 0,012209 | 0,006126 | 32106 | 0,012209 |
| 15 | 6 | 0,012047 | 0,005965 | 35629 | 0,012047 |
| 5 | 9 | 0,011962 | 0,005692 | 29873 | 0,011962 |
| 8 | 12 | 0,011919 | 0,005677 | 34582 | 0,011919 |
| 7 | 13 | 0,011829 | 0,005477 | 32338 | 0,011829 |
| 1 | 11 | 0,01179 | 0,005744 | 32355 | 0,01179 |
| 3 | 11 | 0,011764 | 0,005597 | 36390 | 0,011764 |
| 10 | 11 | 0,011704 | 0,005492 | 39456 | 0,011704 |
| 4 | 8 | 0,01165 | 0,005404 | 30041 | 0,01165 |
| 12 | 3 | 0,011388 | 0,005755 | 45363 | 0,011388 |
| 16 | 7 | 0,011303 | 0,005907 | 27123 | 0,011303 |
| 8 | 2 | 0,011238 | 0,006339 | 49473 | 0,011238 |
| 5 | 15 | 0,010875 | 0,00567 | 34493 | 0,010875 |
| 5 | 11 | 0,010696 | 0,005717 | 47950 | 0,010696 |
| 9 | 12 | 0,010691 | 0,005611 | 31916 | 0,010691 |
| 1 | 9 | 0,010671 | 0,005407 | 33145 | 0,010671 |
| 4 | 14 | 0,010585 | 0,005281 | 33223 | 0,010585 |
| 1 | 21 | 0,010526 | 0,005779 | 33252 | 0,010526 |
| 3 | 18 | 0,010459 | 0,005806 | 28534 | 0,010459 |
| 5 | 7 | 0,010457 | 0,005421 | 27332 | 0,010457 |
| 18 | 7 | 0,010446 | 0,006109 | 39867 | 0,010446 |
| 16 | 5 | 0,010438 | 0,005618 | 21206 | 0,010438 |
| 2 | 7 | 0,01037 | 0,005256 | 31481 | 0,01037 |

| | | | | | |
|---|---|---|---|---|---|
| 20 | 5 | 0,010364 | 0,005718 | 31290 | 0,010364 |
| 2 | 13 | 0,010291 | 0,005277 | 29210 | 0,010291 |
| 12 | 7 | 0,01029 | 0,005599 | 37470 | 0,01029 |
| 2 | 23 | 0,010263 | 0,005505 | 32375 | 0,010263 |
| 1 | 19 | 0,010199 | 0,004829 | 33607 | 0,010199 |
| 7 | 7 | 0,010145 | 0,005081 | 29208 | 0,010145 |
| 1 | 7 | 0,010044 | 0,005067 | 31708 | 0,010044 |
| 16 | 6 | 0,010007 | 0,005427 | 31092 | 0,010007 |
| 5 | 13 | 0,010007 | 0,005022 | 36294 | 0,010007 |
| 14 | 4 | 0,009909 | 0,004636 | 37929 | 0,009909 |
| 3 | 0 | 0,009879 | 0,005686 | 24606 | 0,009879 |
| 3 | 15 | 0,00987 | 0,005664 | 37374 | 0,00987 |
| 1 | 17 | 0,009693 | 0,005588 | 38655 | 0,009693 |
| 9 | 4 | 0,009683 | 0,004946 | 13433 | 0,009683 |
| 4 | 12 | 0,009665 | 0,005025 | 39801 | 0,009665 |
| 15 | 5 | 0,009536 | 0,004954 | 30954 | 0,009536 |
| 17 | 3 | 0,009486 | 0,005039 | 20225 | 0,009486 |
| 4 | 16 | 0,009407 | 0,005249 | 39545 | 0,009407 |
| 1 | 18 | 0,009361 | 0,005318 | 36324 | 0,009361 |
| 19 | 5 | 0,009336 | 0,00517 | 18571 | 0,009336 |
| 13 | 7 | 0,009335 | 0,005488 | 41982 | 0,009335 |
| 7 | 4 | 0,009301 | 0,005457 | 34071 | 0,009301 |
| 2 | 2 | 0,009203 | 0,005499 | 32605 | 0,009203 |
| 10 | 10 | 0,009142 | 0,004932 | 33098 | 0,009142 |
| 1 | 2 | 0,009129 | 0,005082 | 21200 | 0,009129 |
| 3 | 12 | 0,00907 | 0,005402 | 38872 | 0,00907 |
| 10 | 3 | 0,009065 | 0,005636 | 41783 | 0,009065 |
| 22 | 4 | 0,008896 | 0,004348 | 17591 | 0,008896 |
| 4 | 11 | 0,008844 | 0,005001 | 31736 | 0,008844 |
| 13 | 5 | 0,008775 | 0,005299 | 37150 | 0,008775 |
| 15 | 4 | 0,008658 | 0,004903 | 24492 | 0,008658 |
| 12 | 9 | 0,008434 | 0,005026 | 30366 | 0,008434 |
| 10 | 5 | 0,008397 | 0,004681 | 23324 | 0,008397 |
| 6 | 9 | 0,008251 | 0,005618 | 43905 | 0,008251 |
| 17 | 5 | 0,008189 | 0,004791 | 27295 | 0,008189 |
| 16 | 2 | 0,008101 | 0,004786 | 20028 | 0,008101 |
| 20 | 2 | 0,008091 | 0,005824 | 38519 | 0,008091 |
| 4 | 2 | 0,008035 | 0,005402 | 40563 | 0,008035 |
| 15 | 9 | 0,007995 | 0,005995 | 35880 | 0,007995 |
| 6 | 8 | 0,007994 | 0,004584 | 39646 | 0,007994 |
| 15 | 7 | 0,007784 | 0,004599 | 19699 | 0,007784 |
| 7 | 15 | 0,00753 | 0,004778 | 28347 | 0,00753 |
| 3 | 5 | 0,007518 | 0,00448 | 17289 | 0,007518 |
| 20 | 1 | 0,007511 | 0,005814 | 35181 | 0,007511 |
| 5 | 5 | 0,007501 | 0,004446 | 25642 | 0,007501 |
| 4 | 4 | 0,007424 | 0,005381 | 44603 | 0,007424 |
| 11 | 1 | 0,00737 | 0,005475 | 41831 | 0,00737 |
| 3 | 9 | 0,00735 | 0,003581 | 22541 | 0,00735 |
| 1 | 25 | 0,007239 | 0,00389 | 15193 | 0,007239 |
| 14 | 9 | 0,007217 | 0,005263 | 31254 | 0,007217 |
| 3 | 17 | 0,007214 | 0,004742 | 33632 | 0,007214 |
| 8 | 7 | 0,007187 | 0,005689 | 36691 | 0,007187 |
| 5 | 14 | 0,007089 | 0,004769 | 30920 | 0,007089 |
| 8 | 8 | 0,007048 | 0,004976 | 39148 | 0,007048 |
| 1 | 8 | 0,006924 | 0,0051 | 37446 | 0,006924 |
| 7 | 14 | 0,006909 | 0,004769 | 25245 | 0,006909 |

| | | | | | |
|---|---|---|---|---|---|
| 11 | 3 | 0,006905 | 0,004767 | 35811 | 0,006905 |
| 13 | 6 | 0,006839 | 0,00428 | 40089 | 0,006839 |
| 12 | 5 | 0,006772 | 0,00466 | 28179 | 0,006772 |
| 3 | 10 | 0,006762 | 0,004277 | 35926 | 0,006762 |
| 9 | 13 | 0,006749 | 0,004044 | 15332 | 0,006749 |
| 5 | 1 | 0,006735 | 0,005466 | 40201 | 0,006735 |
| 3 | 13 | 0,006712 | 0,004731 | 29995 | 0,006712 |
| 1 | 4 | 0,006609 | 0,004696 | 21946 | 0,006609 |
| 17 | 0 | 0,006522 | 0,003695 | 6922 | 0,006522 |
| 6 | 13 | 0,006497 | 0,005148 | 35571 | 0,006497 |
| 10 | 7 | 0,006491 | 0,004492 | 33566 | 0,006491 |
| 14 | 8 | 0,006348 | 0,005632 | 38271 | 0,006348 |
| 6 | 15 | 0,00629 | 0,005296 | 39141 | 0,00629 |
| 13 | 2 | 0,005928 | 0,004494 | 19666 | 0,005928 |
| 2 | 24 | 0,005914 | 0,00435 | 18370 | 0,005914 |
| 4 | 5 | 0,005899 | 0,004137 | 21451 | 0,005899 |
| 13 | 8 | 0,005894 | 0,004864 | 39106 | 0,005894 |
| 11 | 0 | 0,005842 | 0,003444 | 7908 | 0,005842 |
| 2 | 3 | 0,005784 | 0,004633 | 32815 | 0,005784 |
| 17 | 7 | 0,005726 | 0,004626 | 22744 | 0,005726 |
| 7 | 9 | 0,00571 | 0,004468 | 31283 | 0,00571 |
| 4 | 3 | 0,00553 | 0,004045 | 33127 | 0,00553 |
| 21 | 3 | 0,005516 | 0,00524 | 33510 | 0,005516 |
| 12 | 2 | 0,005459 | 0,004974 | 35361 | 0,005459 |
| 3 | 16 | 0,005347 | 0,004135 | 38609 | 0,005347 |
| 12 | 4 | 0,005322 | 0,003713 | 28586 | 0,005322 |
| 6 | 12 | 0,005309 | 0,004073 | 44148 | 0,005309 |
| 1 | 12 | 0,005302 | 0,003712 | 18144 | 0,005302 |
| 4 | 15 | 0,005224 | 0,004603 | 28574 | 0,005224 |
| 22 | 5 | 0,005218 | 0,003921 | 12155 | 0,005218 |
| 1 | 1 | 0,005097 | 0,00425 | 16580 | 0,005097 |
| 20 | 3 | 0,005053 | 0,003065 | 11026 | 0,005053 |
| 22 | 2 | 0,00504 | 0,003267 | 8986 | 0,00504 |
| 8 | 9 | 0,005011 | 0,00414 | 27510 | 0,005011 |
| 11 | 9 | 0,004936 | 0,004639 | 39738 | 0,004936 |
| 19 | 1 | 0,004936 | 0,004105 | 13071 | 0,004936 |
| 5 | 2 | 0,004936 | 0,003799 | 27636 | 0,004936 |
| 1 | 23 | 0,004918 | 0,005236 | 39247 | 0,004918 |
| 10 | 8 | 0,004797 | 0,00416 | 27051 | 0,004797 |
| 4 | 7 | 0,004762 | 0,004043 | 31920 | 0,004762 |
| 19 | 3 | 0,004743 | 0,003608 | 21196 | 0,004743 |
| 19 | 4 | 0,004737 | 0,0038 | 20714 | 0,004737 |
| 6 | 6 | 0,004697 | 0,003285 | 19009 | 0,004697 |
| 4 | 9 | 0,00433 | 0,004341 | 36849 | 0,00433 |
| 16 | 3 | 0,00423 | 0,004053 | 12719 | 0,00423 |
| 2 | 16 | 0,004229 | 0,004139 | 32228 | 0,004229 |
| 7 | 11 | 0,004205 | 0,004615 | 30486 | 0,004205 |
| 12 | 1 | 0,004146 | 0,005101 | 30343 | 0,004146 |
| 18 | 3 | 0,004125 | 0,004251 | 32477 | 0,004125 |
| 20 | 6 | 0,004112 | 0,004165 | 14939 | 0,004112 |
| 8 | 6 | 0,004016 | 0,004786 | 41305 | 0,004016 |
| 5 | 10 | 0,003974 | 0,003771 | 38635 | 0,003974 |
| 6 | 17 | 0,003823 | 0,004067 | 22409 | 0,003823 |
| 7 | 16 | 0,003793 | 0,002937 | 8682 | 0,003793 |
| 1 | 5 | 0,003783 | 0,00353 | 22219 | 0,003783 |
| 2 | 8 | 0,003764 | 0,004507 | 40971 | 0,003764 |

| | | | | | |
|---|---|---|---|---|---|
| 14 | 2 | 0,003688 | 0,004045 | 14111 | 0,003688 |
| 7 | 8 | 0,003622 | 0,004346 | 30903 | 0,003622 |
| 20 | 0 | 0,0036 | 0,003809 | 15064 | 0,0036 |
| 5 | 0 | 0,003434 | 0,003582 | 13014 | 0,003434 |
| 13 | 4 | 0,003357 | 0,004582 | 35891 | 0,003357 |
| 12 | 0 | 0,003329 | 0,003821 | 11567 | 0,003329 |
| 2 | 15 | 0,003314 | 0,004278 | 31894 | 0,003314 |
| 7 | 12 | 0,003272 | 0,003644 | 25142 | 0,003272 |
| 10 | 0 | 0,003224 | 0,004264 | 21179 | 0,003224 |
| 1 | 10 | 0,003012 | 0,004249 | 36505 | 0,003012 |
| 6 | 0 | 0,002926 | 0,004048 | 15171 | 0,002926 |
| 6 | 7 | 0,002869 | 0,004093 | 48516 | 0,002869 |
| 2 | 0 | 0,002805 | 0,003775 | 18860 | 0,002805 |
| 21 | 5 | 0,002688 | 0,002293 | 5012 | 0,002688 |
| 2 | 1 | 0,002557 | 0,004576 | 29563 | 0,002557 |
| 8 | 11 | 0,002556 | 0,004312 | 38050 | 0,002556 |
| 11 | 12 | 0,002312 | 0,004314 | 31285 | 0,002312 |
| 2 | 19 | 0,002298 | 0,003187 | 26766 | 0,002298 |
| 10 | 9 | 0,002223 | 0,004133 | 38221 | 0,002223 |
| 3 | 4 | 0,002145 | 0,004171 | 33268 | 0,002145 |
| 15 | 3 | 0,002026 | 0,004723 | 23874 | 0,002026 |
| 1 | 22 | 0,002011 | 0,004606 | 38449 | 0,002011 |
| 18 | 0 | 0,001978 | 0,00414 | 17008 | 0,001978 |
| 2 | 10 | 0,001965 | 0,003534 | 21122 | 0,001965 |
| 11 | 4 | 0,001855 | 0,004349 | 37436 | 0,001855 |
| 17 | 6 | 0,001796 | 0,003139 | 16982 | 0,001796 |
| 11 | 6 | 0,001786 | 0,002391 | 26515 | 0,001786 |
| 4 | 17 | 0,001633 | 0,00404 | 32413 | 0,001633 |
| 19 | 2 | 0,001567 | 0,002538 | 20506 | 0,001567 |
| 14 | 11 | 0,001512 | 0,001517 | 1553 | 0,001512 |
| 17 | 8 | 0,001689 | 0,002423 | 7923 | 0,001489 |
| 17 | 2 | 0,001289 | 0,002322 | 7923 | 0,001489 |
| 18 | 2 | 0,001465 | 0,003561 | 17911 | 0,001465 |
| 14 | 6 | 0,001427 | 0,003947 | 34162 | 0,001427 |
| 19 | 0 | 0,001363 | 0,001718 | 1399 | 0,001363 |
| 6 | 5 | 0,00129 | 0,004227 | 43132 | 0,00129 |
| 18 | 8 | 0,001251 | 0,002563 | 8701 | 0,001251 |
| 1 | 15 | 0,002036 | 0,002845 | 12889 | 0,0011215 |
| 11 | 5 | 0,000207 | 0,000966 | 12889 | 0,0011215 |
| 2 | 20 | 0,000828 | 0,003037 | 23677 | 0,000828 |
| 17 | 4 | 0,000803 | 0,001717 | 15442 | 0,000803 |
| 7 | 3 | 0,000769 | 0,00508 | 39264 | 0,000769 |
| 8 | 4 | 0,000704 | 0,003323 | 19122 | 0,000704 |
| 1 | 16 | 0,000655 | 0,004362 | 29155 | 0,000655 |
| 4 | 0 | 0,000536 | 0,002138 | 9291 | 0,000536 |
| 15 | 2 | 0,000533 | 0,001581 | 6214 | 0,000533 |
| 16 | 9 | 0,000182 | 0,002278 | 7024 | 0,000182 |
| 20 | 4 | 0,000152 | 0,004118 | 28100 | 0,000152 |
| 1 | 3 | 1,00E-06 | 0,003311 | 16671 | 0,000001 |
| 12 | 8 | 1,00E-06 | 0,003738 | 33801 | 0,000001 |
| 16 | 0 | 1,00E-06 | 0,001436 | 3317 | 0,000001 |
| 18 | 6 | 1,00E-06 | 0,004855 | 38527 | 0,000001 |
| 6 | 11 | 1,00E-06 | 0,004377 | 34378 | 0,000001 |
| 9 | 9 | 1,00E-06 | 0,004633 | 35614 | 0,000001 |
| 9 | 14 | 1,00E-06 | 0,002598 | 4566 | 0,000001 |
| 11 | 7 | 1,00E-06 | 0,002598 | 13974 | 0,000001 |

| | | | | | |
|---|---|---|---|---|---|
| 7 | 6 | 1,00E-06 | 0,001703 | 13412 | 0,000001 |
| 8 | 5 | 1,00E-06 | 0,001902 | 16150 | 0,000001 |
| 9 | 7 | 1,00E-06 | 0,002927 | 14780 | 0,000001 |
| 19 | 6 | 1,00E-06 | 0,003193 | 10129 | 0,000001 |
| 7 | 0 | 1,00E-06 | 0,002176 | 11534 | 0,000001 |
| 8 | 15 | 1,00E-06 | 0,000555 | 487 | 0,000001 |
| 10 | 4 | 1,00E-06 | 0,002638 | 22637 | 0,000001 |
| 2 | 12 | 1,00E-06 | 0,003412 | 34512 | 0,000001 |
| 3 | 20 | 1,00E-06 | 0,002319 | 5903 | 0,000001 |
| 1 | 0 | 1,00E-06 | 0,002129 | 5178 | 0,000001 |
| 21 | 1 | 1,00E-06 | 0,000981 | 64 | 0,000001 |
| 5 | 18 | 1,00E-06 | 0,002638 | 9279 | 0,000001 |
| 16 | 4 | 1,00E-06 | 0,00031 | 3 | 0,000001 |
| 7 | 5 | 1,00E-06 | 0,004671 | 52206 | 0,000001 |
| 13 | 12 | 1,00E-06 | 0,000999 | 364 | 0,000001 |
| 2 | 9 | 1,00E-06 | 0,002204 | 7576 | 0,000001 |
| 12 | 11 | 1,00E-06 | 0,003211 | 30905 | 0,000001 |

$h^2$: narrow sense heritability adjusted by sex, age, prevalence and PCs; MB10: number of segment within a particular chromosome
