## Supplementary Table 2 for "Genome-wide estimates of heritability and genetic correlations in Essential Tremor"

**Supplementary Table 2. Linkage disequilibrium score regression analyses performed on 832 GWAS**

| Trait | Source | rg | se | z | p-value | h2_obs | h2_obs_se | h2_int | h2_int_se | gcov_int | gcov_int_se |
|---|---|---|---|---|---|---|---|---|---|---|---|
| 18:2 linoleic acid (LA) | 27005778 | 0.068 | 0.1835 | 0.3703 | 0.7111 | 0.1487 | 0.0483 | 0.9928 | 0.0091 | -0.0042 | 0.0061 |
| 22:6 docosahexaenoic acid | 27005778 | 0.1221 | 0.1925 | 0.6341 | 0.526 | 0.1345 | 0.038 | 0.9901 | 0.0086 | -0.0118 | 0.0061 |
| 2hr glucose adjusted for BMI | 20081857 | -0.1136 | 0.1746 | -0.6504 | 0.5154 | 0.1063 | 0.0353 | 0.9911 | 0.0082 | 0.0038 | 0.0053 |
| Acetate | 27005778 | 0.1726 | 0.192 | 0.8986 | 0.3689 | 0.0468 | 0.0185 | 10.049 | 0.008 | -0.001 | 0.0049 |
| Acetoacetate | 27005778 | 0.0294 | 0.1643 | 0.1789 | 0.858 | 0.0829 | 0.0317 | 0.9742 | 0.0082 | -0.001 | 0.0047 |
| Adiponectin | 22479202 | 0.0306 | 0.1441 | 0.2124 | 0.8318 | 0.1128 | 0.0264 | 1.019 | 0.0137 | -0.0018 | 0.0061 |
| Adopted as a child | 0 | 0.1936 | 0.1602 | 12.088 | 0.2267 | 0.0056 | 0.0013 | 10.054 | 0.0075 | -0.0058 | 0.006 |
| Age at first live birth | 0 | -0.0088 | 0.0641 | -0.1366 | 0.8914 | 0.1687 | 0.0083 | 10.411 | 0.0109 | -0.0071 | 0.0067 |
| Age at last live birth | 0 | 0.0718 | 0.0749 | 0.9586 | 0.3378 | 0.0882 | 0.0067 | 10.178 | 0.0101 | -0.0085 | 0.0058 |
| Age at Menarche | 25231870 | -0.0703 | 0.0554 | -12.694 | 0.2043 | 0.2094 | 0.0114 | 0.9323 | 0.0135 | 0.0066 | 0.0064 |
| Age at Menopause | 26414677 | -0.1526 | 0.0855 | -17.854 | 0.0742 | 0.1392 | 0.0174 | 0.9872 | 0.0164 | 0.0032 | 0.0062 |
| Age completed full time education | 0 | 0.0461 | 0.0704 | 0.6553 | 0.5123 | 0.0856 | 0.0049 | 10.574 | 0.0123 | -0.0033 | 0.007 |
| Age of first birth | 27798627 | -0.1124 | 0.0751 | -14.962 | 0.1346 | 0.0607 | 0.0037 | 0.9588 | 0.0094 | 0.0036 | 0.0064 |
| Age of smoking initiation | 20418890 | -0.0298 | 0.2015 | -0.1478 | 0.8825 | 0.0541 | 0.0183 | 10.029 | 0.0075 | -0.0056 | 0.0058 |
| Age started oral contraceptive pill | 0 | -0.1302 | 0.0956 | -13.609 | 0.1735 | 0.047 | 0.0046 | 10.144 | 0.0093 | 0.0014 | 0.0063 |
| Age when periods started (menarche) | 0 | -0.0593 | 0.0473 | -12.526 | 0.2103 | 0.2103 | 0.0119 | 0.9932 | 0.0153 | 0.0066 | 0.0067 |
| Alanine | 27005778 | -0.1181 | 0.1579 | -0.7479 | 0.4545 | 0.0964 | 0.0303 | 10.021 | 0.0089 | 0.0013 | 0.0056 |
| Albumin | 27005778 | -0.4276 | 0.2033 | -21.033 | 0.0354 | 0.0688 | 0.0282 | 0.983 | 0.008 | 0.0076 | 0.0057 |
| Alcohol drinker status: Never | 0 | 0.066 | 0.1114 | 0.5919 | 0.5539 | 0.0141 | 0.0018 | 10.175 | 0.0086 | -0.0013 | 0.0059 |
| Alcohol drinker status: Previous | 0 | -0.0278 | 0.1238 | -0.2242 | 0.8226 | 0.014 | 0.0016 | 10.023 | 0.0078 | -0.0024 | 0.0062 |
| Alcohol intake frequency. | 0 | -0.0664 | 0.0575 | -11.546 | 0.2483 | 0.0841 | 0.0041 | 10.344 | 0.0128 | 0.0049 | 0.0068 |
| Alcohol intake versus 10 years previously | 0 | -0.1162 | 0.0806 | -14.421 | 0.1493 | 0.0358 | 0.0028 | 10.213 | 0.0109 | 0.0111 | 0.0066 |
| Alcohol usually taken with meals | 0 | -0.0112 | 0.0593 | -0.1897 | 0.8495 | 0.1136 | 0.0057 | 10.497 | 0.0115 | 0.0009 | 0.0061 |
| Alzheimers disease | 24162737 | 0.1319 | 0.1408 | 0.9369 | 0.3488 | 0.0575 | 0.013 | 10.424 | 0.0145 | -0.0008 | 0.0066 |
| Amyotrophic lateral sclerosis | 27455348 | 0.1545 | 0.2099 | 0.7361 | 0.4617 | 0.0394 | 0.0141 | 1 | 0.0074 | 0.0078 | 0.0055 |
| Anorexia Nervosa | 24514567 | 0.1666 | 0.0888 | 18.757 | 0.0607 | 0.4338 | 0.0348 | 0.9418 | 0.0087 | 0.0246 | 0.0063 |
| Apolipoprotein A-I | 27005778 | -0.067 | 0.1978 | -0.3387 | 0.7349 | 0.0774 | 0.0285 | 10.014 | 0.0116 | -0.0029 | 0.006 |
| Apolipoprotein B | 27005778 | 0.0135 | 0.1746 | 0.0773 | 0.9384 | 0.1177 | 0.036 | 0.9889 | 0.0107 | -0.0035 | 0.0064 |
| Arm fat mass (left) | 0 | 0.0178 | 0.0499 | 0.3565 | 0.7215 | 0.2339 | 0.0087 | 10.404 | 0.0216 | 0.0024 | 0.0083 |
| Arm fat mass (right) | 0 | 0.0122 | 0.0498 | 0.2447 | 0.8067 | 0.2335 | 0.0087 | 10.418 | 0.0215 | 0.0031 | 0.0083 |
| Arm fat percentage (left) | 0 | 0.0024 | 0.0513 | 0.0476 | 0.9621 | 0.2211 | 0.008 | 1.048 | 0.0208 | -0.002 | 0.0089 |
| Arm fat percentage (right) | 0 | -0.0072 | 0.0513 | -0.1405 | 0.8882 | 0.2176 | 0.0078 | 10.538 | 0.0205 | -0.0003 | 0.0088 |
| Arm fat-free mass (left) | 0 | 0.0545 | 0.0497 | 10.969 | 0.2727 | 0.2659 | 0.012 | 10.582 | 0.0322 | 0.0084 | 0.008 |
| Arm fat-free mass (right) | 0 | 0.0588 | 0.0504 | 11.672 | 0.2431 | 0.2674 | 0.0121 | 10.464 | 0.0329 | 0.0086 | 0.0083 |
| Arm predicted mass (left) | 0 | 0.0559 | 0.0498 | 11.238 | 0.2611 | 0.265 | 0.012 | 10.599 | 0.0324 | 0.0083 | 0.0081 |
| Arm predicted mass (right) | 0 | 0.07 | 0.0496 | 1.413 | 0.1577 | 0.2661 | 0.0121 | 10.474 | 0.0332 | 0.0062 | 0.0081 |
| Asthma | 17611496 | -0.0437 | 0.132 | -0.3312 | 0.7405 | 0.1258 | 0.0283 | 10.024 | 0.0103 | 0.0096 | 0.0062 |
| Attention deficit hyperactivity disorder | 20732625 | 0.1048 | 0.1937 | 0.5412 | 0.5884 | 0.2382 | 0.0987 | 10.112 | 0.0077 | -0.0083 | 0.0053 |
| Attention deficit hyperactivity disorder (GC) | 27663945 | 0.0398 | 0.1963 | 0.2026 | 0.8394 | 0.0856 | 0.0294 | 0.9876 | 0.0075 | -0.007 | 0.006 |
| Attention deficit hyperactivity disorder (No GC) | 27663945 | 0.037 | 0.1969 | 0.1877 | 0.8511 | 0.0867 | 0.0298 | 10.022 | 0.0076 | -0.0069 | 0.0061 |
| Autism spectrum disorder | 0 | -0.0823 | 0.1153 | -0.7134 | 0.4756 | 0.3942 | 0.0551 | 0.9797 | 0.0084 | -0.0016 | 0.0057 |
| Average number of double bonds in a fatty acid chain | 27005778 | 0.0846 | 0.1509 | 0.5605 | 0.5751 | 0.1857 | 0.0457 | 0.9869 | 0.0077 | -0.004 | 0.0057 |
| Average number of methylene groups per a double bond | 27005778 | -0.022 | 0.1492 | -0.1475 | 0.8827 | 0.2093 | 0.0621 | 0.9796 | 0.008 | 0.0032 | 0.0059 |
| Average weekly beer plus cider intake | 0 | -0.0129 | 0.0776 | -0.1668 | 0.8675 | 0.0521 | 0.0036 | 10.615 | 0.0116 | 9,17E-02 | 0.0061 |

| | | | | | | | | | | | |
|---|---|---|---|---|---|---|---|---|---|---|---|
| Average weekly champagne plus white wine intake | 0 | 0.0614 | 0.0892 | 0.6889 | 0.4909 | 0.0309 | 0.0029 | 10.395 | 0.0095 | 0.0044 | 0.0061 |
| Average weekly fortified wine intake | 0 | -0.049 | 0.1345 | -0.3645 | 0.7155 | 0.0119 | 0.0023 | 10.181 | 0.0077 | -0.0012 | 0.0055 |
| Average weekly intake of other alcoholic drinks | 0 | -0.0451 | 0.2585 | -0.1743 | 0.8616 | 0.0083 | 0.0063 | 10.017 | 0.0076 | 0.0008 | 0.0054 |
| Average weekly red wine intake | 0 | 0.0058 | 0.069 | 0.0842 | 0.9329 | 0.0579 | 0.0039 | 10.088 | 0.0103 | -0.0067 | 0.006 |
| Average weekly spirits intake | 0 | -0.088 | 0.0972 | -0.9058 | 0.365 | 0.0298 | 0.0028 | 10.244 | 0.0093 | 0.0084 | 0.0066 |
| Back pain for 3+ months | 0 | -0.0073 | 0.1341 | -0.0547 | 0.9564 | 0.0396 | 0.0064 | 0.9939 | 0.0081 | -0.0023 | 0.0055 |
| Basal metabolic rate | 0 | 0.0635 | 0.0487 | 13.027 | 0.1927 | 0.2869 | 0.0129 | 10.524 | 0.0347 | 0.0064 | 0.0081 |
| Bilateral oophorectomy (both ovaries removed) | 0 | -0.3948 | 0.1336 | -29.548 | 0.0031 | 0.0235 | 0.0036 | 10.048 | 0.0091 | 0.021 | 0.006 |
| Bipolar disorder | 21926972 | 0.0202 | 0.0982 | 0.2054 | 0.8373 | 0.4265 | 0.0373 | 1.02 | 0.0086 | 0.0209 | 0.0058 |
| Birth weight | 27680694 | 0.0478 | 0.0755 | 0.6339 | 0.5261 | 0.1026 | 0.0074 | 10.334 | 0.0117 | 0.0073 | 0.0062 |
| Birth weight | 0 | 0.1113 | 0.0676 | 16.454 | 0.0999 | 0.1102 | 0.0066 | 10.297 | 0.0159 | -0.0031 | 0.0068 |
| Birth weight of first child | 0 | 0.0326 | 0.0698 | 0.4679 | 0.6399 | 0.1102 | 0.0071 | 10.071 | 0.0117 | -0.0009 | 0.0065 |
| Blood clot_ DVT_ bronchitis_ emphysema_ asthma_ rhinitis_ e | 0 | -0.0556 | 0.0647 | -0.8593 | 0.3902 | 0.0573 | 0.0061 | 10.125 | 0.0189 | 0.0038 | 0.0065 |
| Blood clot_ DVT_ bronchitis_ emphysema_ asthma_ rhinitis_ e | 0 | 0.1003 | 0.1206 | 0.8314 | 0.4057 | 0.0116 | 0.0018 | 0.9994 | 0.0107 | -0.0066 | 0.006 |
| Blood clot_ DVT_ bronchitis_ emphysema_ asthma_ rhinitis_ e | 0 | 0.0753 | 0.1531 | 0.4915 | 0.6231 | 0.0063 | 0.0015 | 10.017 | 0.0083 | -7,36E-01 | 0.0057 |
| Blood clot_ DVT_ bronchitis_ emphysema_ asthma_ rhinitis_ e | 0 | 0.0792 | 0.1189 | 0.6656 | 0.5057 | 0.0126 | 0.002 | 0.9828 | 0.0092 | -0.0037 | 0.0062 |
| Blood clot_ DVT_ bronchitis_ emphysema_ asthma_ rhinitis_ e | 0 | 0.0271 | 0.0575 | 0.4711 | 0.6375 | 0.0733 | 0.0063 | 1.025 | 0.0199 | -0.0019 | 0.0066 |
| Blood clot_ DVT_ bronchitis_ emphysema_ asthma_ rhinitis_ e | 0 | -0.0334 | 0.0554 | -0.6043 | 0.5457 | 0.0774 | 0.0073 | 10.169 | 0.0211 | 0.004 | 0.0069 |
| Body fat | 26833246 | 0.0849 | 0.0713 | 11.904 | 0.2339 | 0.1106 | 0.0089 | 0.8959 | 0.0081 | -0.0054 | 0.0055 |
| Body fat percentage | 0 | -0.0063 | 0.0499 | -0.1265 | 0.8994 | 0.2183 | 0.0075 | 10.552 | 0.0203 | 0.0008 | 0.0087 |
| Body mass index | 20935630 | 0.1024 | 0.0611 | 16.758 | 0.0938 | 0.1903 | 0.0103 | 10.052 | 0.0121 | -0.0039 | 0.0066 |
| Body mass index (BMI) | 0 | -0.009 | 0.0495 | -0.1822 | 0.8554 | 0.2505 | 0.0094 | 10.399 | 0.022 | 0.0011 | 0.0086 |
| Breastfed as a baby | 0 | -0.0838 | 0.0984 | -0.8519 | 0.3943 | 0.0252 | 0.0027 | 10.259 | 0.0093 | 0.0074 | 0.0062 |
| Bring up phlegm/sputum/mucus on most days | 0 | -0.0494 | 0.1567 | -0.3154 | 0.7525 | 0.0263 | 0.0066 | 10.084 | 0.0088 | -0.0006 | 0.0058 |
| Cancer code_ self-reported: basal cell carcinoma | 0 | -0.0601 | 0.1068 | -0.563 | 0.5734 | 0.0114 | 0.002 | 0.9879 | 0.0087 | 0.0024 | 0.0054 |
| Cancer code_ self-reported: breast cancer | 0 | 0.0533 | 0.1239 | 0.4306 | 0.6667 | 0.0089 | 0.0019 | 10.248 | 0.0099 | -0.0113 | 0.0053 |
| Cancer code_ self-reported: lung cancer | 0 | -0.0821 | 0.2546 | -0.3225 | 0.7471 | 0.0018 | 0.0013 | 0.9825 | 0.0074 | -4,13E-01 | 0.0057 |
| Cancer code_ self-reported: malignant melanoma | 0 | -0.0317 | 0.1718 | -0.1844 | 0.8537 | 0.0058 | 0.0016 | 0.988 | 0.0075 | 0.0019 | 0.0057 |
| Cancer code_ self-reported: prostate cancer | 0 | -0.0916 | 0.17 | -0.5389 | 0.5899 | 0.006 | 0.0016 | 10.119 | 0.0094 | 0.0009 | 0.0059 |
| Cancer code_ self-reported: small intestine/small bowel cancer | 0 | 0.187 | 0.486 | 0.3847 | 0.7004 | 0.001 | 0.0015 | 0.9976 | 0.0075 | -0.0075 | 0.0063 |
| Cancer code_ self-reported: squamous cell carcinoma | 0 | 0.1761 | 0.2812 | 0.6262 | 0.5312 | 0.0022 | 0.0014 | 0.9979 | 0.0079 | -0.002 | 0.0058 |
| Cancer diagnosed by doctor | 0 | -0.0753 | 0.1511 | -0.4982 | 0.6183 | 0.0064 | 0.0017 | 10.115 | 0.0081 | -0.0023 | 0.0058 |
| Celiac disease | 20190752 | 0.0804 | 0.1372 | 0.5862 | 0.5577 | 0.3267 | 0.0537 | 10.555 | 0.012 | -0.0018 | 0.0079 |
| Chest pain or discomfort | 0 | -0.1141 | 0.0796 | -14.339 | 0.1516 | 0.0339 | 0.0025 | 10.093 | 0.0099 | 0.0129 | 0.0064 |
| Chest pain or discomfort walking normally | 0 | -0.1177 | 0.1153 | -10.205 | 0.3075 | 0.0812 | 0.0101 | 10.036 | 0.0082 | 0.0041 | 0.006 |
| Child birth length | 25281659 | -0.0195 | 0.1357 | -0.144 | 0.8855 | 0.1724 | 0.0264 | 0.9889 | 0.008 | 0.0126 | 0.0063 |
| Child birth weight | 23202124 | -0.0214 | 0.1273 | -0.1678 | 0.8667 | 0.114 | 0.0198 | 10.011 | 0.0074 | 0.0041 | 0.0058 |
| Childhood IQ | 23358156 | 0.1607 | 0.1289 | 12.466 | 0.2125 | 0.2551 | 0.0463 | 1.008 | 0.0104 | -0.0038 | 0.0066 |
| Childhood obesity | 22484627 | -0.0108 | 0.1019 | -0.1063 | 0.9153 | 0.3914 | 0.0497 | 0.9328 | 0.009 | 0.0021 | 0.0057 |
| Cholesterol esters in large HDL | 27005778 | -0.0816 | 0.1735 | -0.4702 | 0.6382 | 0.1184 | 0.0311 | 10.012 | 0.0149 | 0.0006 | 0.0064 |
| Cholesterol esters in large LDL | 27005778 | 0.1739 | 0.1729 | 10.059 | 0.3145 | 0.1302 | 0.0369 | 0.9852 | 0.013 | -0.0094 | 0.0063 |
| Cholesterol esters in large VLDL | 27005778 | -0.0178 | 0.1311 | -0.1357 | 0.892 | 0.1647 | 0.0355 | 0.9712 | 0.0083 | -6,28E-01 | 0.0054 |
| Cholesterol esters in medium HDL | 27005778 | -0.331 | 0.2506 | -1.321 | 0.1865 | 0.0554 | 0.0309 | 0.9988 | 0.0102 | 0.0057 | 0.0061 |
| Cholesterol esters in medium LDL | 27005778 | 0.1541 | 0.173 | 0.8907 | 0.3731 | 0.132 | 0.0368 | 0.9847 | 0.0126 | -0.0095 | 0.0063 |
| Cholesterol esters in medium VLDL | 27005778 | 0.037 | 0.151 | 0.2449 | 0.8065 | 0.1541 | 0.0394 | 0.9809 | 0.0086 | -0.0029 | 0.006 |
| Chronic Kidney Disease | 26831199 | -0.1538 | 0.1778 | -0.865 | 0.3871 | 0.0184 | 0.0059 | 10.132 | 0.0097 | 0.0036 | 0.0062 |
| Chronotype | 27494321 | 0.0653 | 0.0709 | 0.9212 | 0.3569 | 0.1049 | 0.006 | 10.081 | 0.0094 | -0.0013 | 0.0057 |

| | | | | | | | | | | | |
|---|---|---|---|---|---|---|---|---|---|---|---|
| Cigarettes smoked per day | 20418890 | -0.0026 | 0.166 | -0.0156 | 0.9875 | 0.0525 | 0.016 | 10.123 | 0.0081 | 0.0013 | 0.0056 |
| Citrate | 27005778 | -0.0844 | 0.1788 | -0.4719 | 0.637 | 0.0649 | 0.023 | 1.008 | 0.0092 | -0.0013 | 0.005 |
| College completion | 23722424 | -0.0203 | 0.0876 | -0.2314 | 0.817 | 0.0794 | 0.0061 | 10.221 | 0.0098 | 0.0022 | 0.0064 |
| Comparative body size at age 10 | 0 | -0.0997 | 0.0515 | -19.362 | 0.0528 | 0.151 | 0.0107 | 0.9879 | 0.0213 | 0.0065 | 0.0071 |
| Comparative height size at age 10 | 0 | 0.1085 | 0.0481 | 22.582 | 0.0239 | 0.246 | 0.0132 | 10.197 | 0.0353 | 0.0021 | 0.0083 |
| Concentration of chylomicrons and largest VLDL particles | 27005778 | -0.1293 | 0.1502 | -0.8607 | 0.3894 | 0.1151 | 0.0307 | 0.9843 | 0.0085 | 0.0043 | 0.0053 |
| Concentration of IDL particles | 27005778 | 0.1015 | 0.1804 | 0.563 | 0.5735 | 0.1101 | 0.0344 | 0.9981 | 0.0123 | -0.0086 | 0.0062 |
| Concentration of large HDL particles | 27005778 | -0.1221 | 0.1684 | -0.7253 | 0.4683 | 0.1271 | 0.0327 | 0.9989 | 0.0164 | 0.0002 | 0.0063 |
| Concentration of large LDL particles | 27005778 | 0.1375 | 0.1757 | 0.7829 | 0.4337 | 0.1248 | 0.0363 | 0.9905 | 0.0125 | -0.0095 | 0.0063 |
| Concentration of large VLDL particles | 27005778 | -0.0371 | 0.1383 | -0.2685 | 0.7883 | 0.1421 | 0.0362 | 0.9656 | 0.0081 | -3,13E-01 | 0.0051 |
| Concentration of medium HDL particles | 27005778 | -0.2829 | 0.2093 | -13.514 | 0.1766 | 0.072 | 0.0286 | 0.9913 | 0.0088 | 0.0032 | 0.0057 |
| Concentration of medium LDL particles | 27005778 | 0.1418 | 0.1768 | 0.8018 | 0.4226 | 0.1278 | 0.0369 | 0.988 | 0.012 | -0.0102 | 0.0064 |
| Concentration of medium VLDL particles | 27005778 | 0.0458 | 0.1394 | 0.3286 | 0.7424 | 0.1506 | 0.0372 | 0.9769 | 0.0087 | -0.002 | 0.0054 |
| Concentration of small LDL particles | 27005778 | 0.0937 | 0.1702 | 0.5506 | 0.5819 | 0.1344 | 0.0378 | 0.9877 | 0.0107 | -0.0089 | 0.0063 |
| Concentration of small VLDL particles | 27005778 | 0.0367 | 0.1479 | 0.2482 | 0.804 | 0.1617 | 0.0387 | 0.9823 | 0.0087 | -0.0042 | 0.006 |
| Concentration of very large HDL particles | 27005778 | 0.0084 | 0.214 | 0.0391 | 0.9688 | 0.0679 | 0.0302 | 1 | 0.0167 | -0.0036 | 0.0059 |
| Concentration of very large VLDL particles | 27005778 | -0.0595 | 0.1318 | -0.4513 | 0.6518 | 0.1304 | 0.0329 | 0.9775 | 0.0086 | 0.0009 | 0.0051 |
| Concentration of very small VLDL particles | 27005778 | -0.0035 | 0.1646 | -0.0216 | 0.9828 | 0.1281 | 0.0364 | 0.9952 | 0.0119 | -0.0063 | 0.0061 |
| Coronary artery disease | 26343387 | 0.0091 | 0.0665 | 0.137 | 0.8911 | 0.0808 | 0.0058 | 10.298 | 0.0114 | 0.0296 | 0.0059 |
| Cough on most days | 0 | 0.0003 | 0.113 | 0.0025 | 0.998 | 0.0514 | 0.0072 | 0.9947 | 0.0084 | -0.0021 | 0.0056 |
| Creatinine | 27005778 | -0.2893 | 0.1496 | -19.329 | 0.0532 | 0.0999 | 0.0277 | 10.167 | 0.0096 | 0.0059 | 0.0053 |
| Creatinine (enzymatic) in urine | 0 | 0.0184 | 0.0585 | 0.3144 | 0.7532 | 0.0664 | 0.0032 | 10.132 | 0.0113 | -0.0025 | 0.0066 |
| Crohns disease | 26192919 | -0.0471 | 0.0644 | -0.7303 | 0.4652 | 0.4983 | 0.0599 | 10.131 | 0.0137 | 0.032 | 0.005 |
| Current employment status: Doing unpaid or voluntary work | 0 | -0.186 | 0.1492 | -12.466 | 0.2126 | 0.0075 | 0.0016 | 10.211 | 0.008 | 0.0037 | 0.0058 |
| Current employment status: In paid employment or self-employe | 0 | 0.243 | 0.1373 | 17.694 | 0.0768 | 0.0079 | 0.0015 | 10.162 | 0.0074 | -0.0016 | 0.0054 |
| Current employment status: Looking after home and/or family | 0 | -0.1422 | 0.1717 | -0.828 | 0.4077 | 0.0052 | 0.0015 | 10.062 | 0.0082 | 0.0054 | 0.006 |
| Current employment status: Retired | 0 | -0.2158 | 0.1324 | -1.63 | 0.1031 | 0.0079 | 0.0016 | 10.179 | 0.0079 | -0.0008 | 0.0052 |
| Current employment status: Unable to work because of sickness | 0 | -0.0791 | 0.0845 | -0.9358 | 0.3494 | 0.0239 | 0.002 | 10.226 | 0.0094 | 0.0072 | 0.0055 |
| Current tobacco smoking | 0 | -0.0451 | 0.0586 | -0.7704 | 0.4411 | 0.0557 | 0.0032 | 10.113 | 0.0111 | 0.0024 | 0.0059 |
| Daytime dozing / sleeping (narcolepsy) | 0 | -0.1004 | 0.0583 | -17.216 | 0.0851 | 0.0469 | 0.0029 | 10.238 | 0.011 | 0.0059 | 0.0063 |
| Depressive symptoms | 27089181 | -0.0373 | 0.0835 | -0.4465 | 0.6552 | 0.0468 | 0.004 | 10.035 | 0.0086 | 0.0151 | 0.0059 |
| Description of average fatty acid chain length; not actual carbon | 27005778 | 0.2315 | 0.1802 | 12.848 | 0.1989 | 0.1328 | 0.037 | 0.9718 | 0.0078 | -0.0081 | 0.0051 |
| Diabetes diagnosed by doctor | 0 | -0.0044 | 0.0742 | -0.0593 | 0.9527 | 0.0424 | 0.0028 | 10.384 | 0.0138 | -0.0019 | 0.0066 |
| Diagnoses - main ICD10: B37 Candidiasis | 0 | -0.312 | 0.433 | -0.7207 | 0.4711 | 0.001 | 0.0014 | 0.997 | 0.0083 | 0.0067 | 0.0054 |
| Diagnoses - main ICD10: C44 Other malignant neoplasms of sk | 0 | 0.0663 | 0.126 | 0.5263 | 0.5987 | 0.0107 | 0.0025 | 0.9998 | 0.0103 | -0.0039 | 0.0059 |
| Diagnoses - main ICD10: C50 Malignant neoplasm of breast | 0 | -0.0062 | 0.1297 | -0.0481 | 0.9616 | 0.0087 | 0.0018 | 10.038 | 0.0092 | -0.0084 | 0.0056 |
| Diagnoses - main ICD10: C61 Malignant neoplasm of prostate | 0 | -0.1921 | 0.1897 | -10.128 | 0.3112 | 0.0044 | 0.0016 | 10.155 | 0.0086 | 0.0044 | 0.0058 |
| Diagnoses - main ICD10: D12 Benign neoplasm of colon_ rectu | 0 | -0.2165 | 0.1414 | -15.315 | 0.1257 | 0.0082 | 0.0016 | 10.053 | 0.0083 | 0.0068 | 0.0054 |
| Diagnoses - main ICD10: D25 Leiomyoma of uterus | 0 | -0.3223 | 0.1666 | -19.346 | 0.053 | 0.0052 | 0.0016 | 0.9971 | 0.0077 | 0.0159 | 0.0058 |
| Diagnoses - main ICD10: E03 Other hypothyroidism | 0 | NA | NA | NA | NA | -0.0008 | 0.0013 | 10.066 | 0.0071 | 0.0033 | 0.0058 |
| Diagnoses - main ICD10: E04 Other non-toxic goitre | 0 | 0.1889 | 0.1598 | 11.823 | 0.2371 | 0.0052 | 0.0015 | 0.9915 | 0.0083 | -0.002 | 0.0055 |
| Diagnoses - main ICD10: F31 Bipolar affective disorder | 0 | -0.2253 | 0.1693 | -13.309 | 0.1832 | 0.0054 | 0.0016 | 0.9865 | 0.0077 | 0.0059 | 0.0058 |
| Diagnoses - main ICD10: F43 Reaction to severe stress_ and ad | 0 | NA | NA | NA | NA | -5,73E-01 | 0.0014 | 10.084 | 0.0071 | -0.0028 | 0.0052 |
| Diagnoses - main ICD10: G47 Sleep disorders | 0 | -0.041 | 0.1465 | -0.2797 | 0.7797 | 0.0072 | 0.0015 | 0.9866 | 0.0078 | -0.0039 | 0.0056 |
| Diagnoses - main ICD10: G56 Mononeuropathies of upper limb | 0 | -0.066 | 0.137 | -0.4814 | 0.6302 | 0.0134 | 0.002 | 1.007 | 0.0087 | -0.0018 | 0.0066 |
| Diagnoses - main ICD10: H25 Senile cataract | 0 | -0.3442 | 0.219 | -15.717 | 0.116 | 0.0034 | 0.0015 | 1.01 | 0.0079 | 0.0054 | 0.0061 |
| Diagnoses - main ICD10: H26 Other cataract | 0 | -0.135 | 0.1283 | -10.523 | 0.2927 | 0.0095 | 0.0017 | 0.9892 | 0.0077 | 0.0085 | 0.0055 |

| | | | | | | | | | | | |
|---|---|---|---|---|---|---|---|---|---|---|---|
| Diagnoses - main ICD10: I10 Essential (primary) hypertension | 0 | -0.2594 | 0.3145 | -0.8249 | 0.4094 | 0.002 | 0.0014 | 10.026 | 0.0075 | -0.0007 | 0.0056 |
| Diagnoses - main ICD10: I20 Angina pectoris | 0 | -0.2817 | 0.1294 | -2.177 | 0.0295 | 0.0096 | 0.0017 | 0.9989 | 0.0084 | 0.0127 | 0.0056 |
| Diagnoses - main ICD10: I21 Acute myocardial infarction | 0 | 0.0097 | 0.1313 | 0.074 | 0.941 | 0.0105 | 0.0018 | 10.028 | 0.0085 | 0.0025 | 0.0063 |
| Diagnoses - main ICD10: I25 Chronic ischaemic heart disease | 0 | -0.0199 | 0.0959 | -0.2076 | 0.8355 | 0.0201 | 0.0022 | 10.146 | 0.0092 | 0.0052 | 0.0058 |
| Diagnoses - main ICD10: I30 Acute pericarditis | 0 | 0.1132 | 0.2761 | 0.41 | 0.6818 | 0.002 | 0.0015 | 0.9955 | 0.008 | -0.003 | 0.0064 |
| Diagnoses - main ICD10: I48 Atrial fibrillation and flutter | 0 | 0.0752 | 0.1089 | 0.6908 | 0.4897 | 0.0134 | 0.0021 | 10.073 | 0.011 | -0.0003 | 0.006 |
| Diagnoses - main ICD10: I80 Phlebitis and thrombophlebitis | 0 | 0.2488 | 0.1373 | 18.129 | 0.0699 | 0.0081 | 0.0017 | 0.993 | 0.0089 | -0.0115 | 0.0061 |
| Diagnoses - main ICD10: I83 Varicose veins of lower extremiti | 0 | 0.1156 | 0.0977 | 11.832 | 0.2367 | 0.0194 | 0.0023 | 10.233 | 0.0098 | -0.0043 | 0.0062 |
| Diagnoses - main ICD10: I84 Haemorrhoids | 0 | -0.1241 | 0.1259 | -0.9859 | 0.3242 | 0.0096 | 0.0019 | 0.9972 | 0.0092 | 0.0053 | 0.0057 |
| Diagnoses - main ICD10: J22 Unspecified acute lower respirato | 0 | 0.0461 | 0.2242 | 0.2056 | 0.8371 | 0.0027 | 0.0017 | 0.9987 | 0.0079 | -0.0028 | 0.0054 |
| Diagnoses - main ICD10: J33 Nasal polyp | 0 | 0.2272 | 0.1903 | 11.939 | 0.2325 | 0.0037 | 0.0017 | 10.041 | 0.0082 | -0.0016 | 0.0054 |
| Diagnoses - main ICD10: J34 Other disorders of nose and nasal | 0 | 0.2604 | 0.3408 | 0.7643 | 0.4447 | 0.0016 | 0.0014 | 10.149 | 0.0077 | -0.0048 | 0.006 |
| Diagnoses - main ICD10: J44 Other chronic obstructive pulmon | 0 | 0.3804 | 0.188 | 20.232 | 0.0431 | 0.0045 | 0.0016 | 0.999 | 0.0081 | -0.0071 | 0.0056 |
| Diagnoses - main ICD10: K20 Oesophagitis | 0 | -0.8359 | 0.4338 | -19.269 | 0.054 | 0.0017 | 0.0014 | 10.084 | 0.0079 | 0.0151 | 0.0058 |
| Diagnoses - main ICD10: K21 Gastro-oesophageal reflux diseas | 0 | 0.0223 | 0.1437 | 0.1552 | 0.8766 | 0.0067 | 0.0014 | 10.009 | 0.0072 | -0.0033 | 0.0058 |
| Diagnoses - main ICD10: K22 Other diseases of oesophagus | 0 | -0.0401 | 0.2005 | -0.2 | 0.8415 | 0.0038 | 0.0017 | 10.058 | 0.0085 | 0.0055 | 0.0058 |
| Diagnoses - main ICD10: K29 Gastritis and duodenitis | 0 | -0.2089 | 0.1407 | -14.846 | 0.1377 | 0.0088 | 0.0015 | 0.9936 | 0.0073 | 0.009 | 0.0058 |
| Diagnoses - main ICD10: K30 Dyspepsia | 0 | -0.0978 | 0.2143 | -0.4566 | 0.648 | 0.0032 | 0.0015 | 10.026 | 0.0079 | 0.0044 | 0.0057 |
| Diagnoses - main ICD10: K35 Acute appendicitis | 0 | 0.4268 | 0.2946 | 14.485 | 0.1475 | 0.0024 | 0.0014 | 10.027 | 0.0081 | -0.0067 | 0.006 |
| Diagnoses - main ICD10: K40 Inguinal hernia | 0 | 0.0662 | 0.0969 | 0.6835 | 0.4943 | 0.0186 | 0.0025 | 0.9921 | 0.0084 | -0.0066 | 0.0058 |
| Diagnoses - main ICD10: K43 Ventral hernia | 0 | -0.2992 | 0.1965 | -15.222 | 0.128 | 0.0047 | 0.0016 | 0.9958 | 0.0085 | 0.0077 | 0.006 |
| Diagnoses - main ICD10: K44 Diaphragmatic hernia | 0 | -0.4697 | 0.2182 | -21.529 | 0.0313 | 0.0049 | 0.0015 | 10.058 | 0.0083 | 0.0134 | 0.0063 |
| Diagnoses - main ICD10: K50 Crohns disease [regional enteriti | 0 | 0.275 | 0.1839 | 14.954 | 0.1348 | 0.0045 | 0.0016 | 10.008 | 0.0083 | -0.0078 | 0.0058 |
| Diagnoses - main ICD10: K51 Ulcerative colitis | 0 | -0.0606 | 0.1813 | -0.3345 | 0.738 | 0.0043 | 0.0017 | 10.119 | 0.0086 | 0.0048 | 0.0056 |
| Diagnoses - main ICD10: K52 Other non-infective gastro-enteri | 0 | 0.0722 | 0.1754 | 0.4117 | 0.6805 | 0.0045 | 0.0014 | 10.055 | 0.0075 | 0.0008 | 0.0059 |
| Diagnoses - main ICD10: K57 Diverticular disease of intestine | 0 | -0.0337 | 0.1064 | -0.3162 | 0.7519 | 0.0143 | 0.0016 | 0.9945 | 0.008 | 0.0006 | 0.0058 |
| Diagnoses - main ICD10: K60 Fissure and fistula of anal and re | 0 | 0.27 | 0.1781 | 15.162 | 0.1295 | 0.005 | 0.0015 | 0.9855 | 0.0079 | -0.0081 | 0.0053 |
| Diagnoses - main ICD10: K62 Other diseases of anus and rectu | 0 | 0.1135 | 0.1503 | 0.7551 | 0.4502 | 0.0058 | 0.0015 | 0.9936 | 0.008 | -0.0036 | 0.0057 |
| Diagnoses - main ICD10: K76 Other diseases of liver | 0 | -0.1061 | 0.2351 | -0.4513 | 0.6517 | 0.0036 | 0.0014 | 0.9924 | 0.0079 | 0.0038 | 0.0065 |
| Diagnoses - main ICD10: K80 Cholelithiasis | 0 | -0.0219 | 0.1291 | -0.1693 | 0.8656 | 0.0092 | 0.002 | 10.216 | 0.0099 | -0.0023 | 0.0059 |
| Diagnoses - main ICD10: L03 Cellulitis | 0 | 0.009 | 0.1943 | 0.0464 | 0.963 | 0.0034 | 0.0017 | 10.063 | 0.008 | -0.0008 | 0.0053 |
| Diagnoses - main ICD10: M10 Gout | 0 | 0.6063 | 0.3182 | 19.052 | 0.0568 | 0.0025 | 0.0015 | 0.9994 | 0.0079 | -0.0133 | 0.0053 |
| Diagnoses - main ICD10: M16 Coxarthrosis [arthrosis of hip] | 0 | -0.0715 | 0.1181 | -0.6055 | 0.5449 | 0.0111 | 0.0018 | 10.055 | 0.0094 | -0.0002 | 0.0061 |
| Diagnoses - main ICD10: M17 Gonarthrosis [arthrosis of knee] | 0 | -0.2627 | 0.1262 | -20.821 | 0.0373 | 0.0134 | 0.0018 | 1.005 | 0.0086 | 0.011 | 0.0063 |
| Diagnoses - main ICD10: M20 Acquired deformities of fingers | 0 | 0.0624 | 0.1383 | 0.4517 | 0.6515 | 0.0103 | 0.0016 | 0.9942 | 0.0091 | 0.0028 | 0.0064 |
| Diagnoses - main ICD10: M21 Other acquired deformities of lir | 0 | NA | NA | NA | NA | -4,17E-01 | 0.0012 | 10.109 | 0.0071 | -0.0057 | 0.0059 |
| Diagnoses - main ICD10: M23 Internal derangement of knee | 0 | 0.1342 | 0.1601 | 0.8382 | 0.4019 | 0.008 | 0.002 | 1.004 | 0.0089 | -0.0013 | 0.0064 |
| Diagnoses - main ICD10: M24 Other specific joint derangemen | 0 | 0.1066 | 0.1992 | 0.5354 | 0.5924 | 0.0034 | 0.0015 | 0.9877 | 0.008 | 0.0057 | 0.0055 |
| Diagnoses - main ICD10: M25 Other joint disorders_ not elsew | 0 | -0.1983 | 0.3339 | -0.594 | 0.5525 | 0.0016 | 0.0014 | 10.116 | 0.0076 | 0.0035 | 0.0059 |
| Diagnoses - main ICD10: M54 Dorsalgia | 0 | 0.2142 | 0.1412 | 15.166 | 0.1294 | 0.0096 | 0.0017 | 10.063 | 0.0088 | -0.0039 | 0.006 |
| Diagnoses - main ICD10: M67 Other disorders of synovium and | 0 | -0.1833 | 0.2339 | -0.7836 | 0.4333 | 0.0028 | 0.0014 | 0.9965 | 0.0081 | 0.0033 | 0.0053 |
| Diagnoses - main ICD10: M70 Soft tissue disorders related to u | 0 | 0.0835 | 0.1912 | 0.4368 | 0.6623 | 0.0043 | 0.0015 | 0.9808 | 0.0076 | -0.0046 | 0.0061 |
| Diagnoses - main ICD10: M72 Fibroblastic disorders | 0 | -0.073 | 0.1248 | -0.5847 | 0.5588 | 0.0116 | 0.0023 | 1.009 | 0.0123 | 0.0031 | 0.006 |
| Diagnoses - main ICD10: N19 Unspecified renal failure | 0 | 0.2472 | 0.2064 | 11.975 | 0.2311 | 0.0031 | 0.0015 | 0.9984 | 0.0074 | -0.0013 | 0.0056 |
| Diagnoses - main ICD10: N20 Calculus of kidney and ureter | 0 | 0.0861 | 0.1523 | 0.5652 | 0.5719 | 0.0076 | 0.0015 | 1.011 | 0.0075 | 0.0021 | 0.0061 |
| Diagnoses - main ICD10: N32 Other disorders of bladder | 0 | -0.3059 | 0.1985 | -15.412 | 0.1233 | 0.0041 | 0.0014 | 0.9871 | 0.0076 | 0.0113 | 0.0055 |
| Diagnoses - main ICD10: N40 Hyperplasia of prostate | 0 | -0.0556 | 0.1541 | -0.3608 | 0.7183 | 0.0065 | 0.0015 | 10.032 | 0.0076 | 0.0047 | 0.0055 |

| | | | | | | | | | | | |
|---|---|---|---|---|---|---|---|---|---|---|---|
| Diagnoses - main ICD10: N81 Female genital prolapse | 0 | -0.1136 | 0.1594 | -0.7128 | 0.476 | 0.0063 | 0.0017 | 10.267 | 0.0083 | -0.0006 | 0.0061 |
| Diagnoses - main ICD10: N92 Excessive_ frequent and irregula | 0 | -0.0616 | 0.1358 | -0.4536 | 0.6501 | 0.0074 | 0.0018 | 0.9988 | 0.0091 | 0.0096 | 0.0054 |
| Diagnoses - main ICD10: O75 Other complications of labour an | 0 | 0.2477 | 0.2558 | 0.9682 | 0.3329 | 0.003 | 0.0013 | 0.9876 | 0.0062 | -0.0011 | 0.0062 |
| Diagnoses - main ICD10: R04 Haemorrhage from respiratory pa | 0 | -0.4954 | 0.2692 | -18.405 | 0.0657 | 0.0026 | 0.0016 | 0.9968 | 0.0076 | 0.0105 | 0.0056 |
| Diagnoses - main ICD10: R07 Pain in throat and chest | 0 | -0.0124 | 0.1101 | -0.1128 | 0.9102 | 0.0164 | 0.0018 | 10.038 | 0.0082 | 0.0022 | 0.0061 |
| Diagnoses - main ICD10: R10 Abdominal and pelvic pain | 0 | -0.0498 | 0.1162 | -0.4288 | 0.6681 | 0.0123 | 0.0017 | 0.9956 | 0.0085 | 0.0052 | 0.0059 |
| Diagnoses - main ICD10: R11 Nausea and vomiting | 0 | -0.404 | 0.3626 | -11.143 | 0.2651 | 0.0013 | 0.0014 | 0.9947 | 0.0079 | 0.0036 | 0.006 |
| Diagnoses - main ICD10: R14 Flatulence and related conditions | 0 | 0.1377 | 0.2203 | 0.625 | 0.532 | 0.0035 | 0.0013 | 0.989 | 0.0076 | -0.0022 | 0.0056 |
| Diagnoses - main ICD10: R31 Unspecified haematuria | 0 | -0.01 | 0.2351 | -0.0423 | 0.9662 | 0.0027 | 0.0016 | 10.095 | 0.0082 | 0.0036 | 0.0052 |
| Diagnoses - main ICD10: R35 Polyuria | 0 | -0.1236 | 0.2182 | -0.5666 | 0.571 | 0.0036 | 0.0016 | 0.9948 | 0.0083 | 0.0052 | 0.0058 |
| Diagnoses - main ICD10: R55 Syncope and collapse | 0 | -0.2233 | 0.1694 | -13.186 | 0.1873 | 0.0056 | 0.0016 | 1.007 | 0.0081 | 0.005 | 0.0056 |
| Diagnoses - main ICD10: R69 Unknown and unspecified causes | 0 | 0.0401 | 0.168 | 0.2385 | 0.8115 | 0.0047 | 0.0014 | 10.043 | 0.008 | 0.0027 | 0.0058 |
| Diagnoses - main ICD10: S09 Other and unspecified injuries of | 0 | -0.4688 | 0.2019 | -23.223 | 0.0202 | 0.0048 | 0.0015 | 0.9859 | 0.0074 | 0.0156 | 0.0063 |
| Diagnoses - main ICD10: S52 Fracture of forearm | 0 | -0.0238 | 0.147 | -0.1617 | 0.8715 | 0.0066 | 0.0018 | 0.9965 | 0.0085 | -0.0058 | 0.0059 |
| Diagnoses - main ICD10: S66 Injury of muscle and tendon at w | 0 | -0.256 | 0.3146 | -0.8138 | 0.4158 | 0.002 | 0.0013 | 1.001 | 0.0078 | 0.0049 | 0.0057 |
| Diagnoses - main ICD10: S76 Injury of muscle and tendon at hi | 0 | 0.2843 | 0.3572 | 0.7957 | 0.4262 | 0.0015 | 0.0013 | 0.9976 | 0.0073 | 0.0004 | 0.006 |
| Diagnoses - main ICD10: T84 Complications of internal orthopa | 0 | -0.044 | 0.1719 | -0.2556 | 0.7982 | 0.0056 | 0.0018 | 0.9844 | 0.0086 | 0.008 | 0.0059 |
| Diagnoses - main ICD10: Z09 Follow-up examination after trea | 0 | -0.2087 | 0.2237 | -0.9329 | 0.3509 | 0.0035 | 0.0014 | 0.9976 | 0.0078 | 0.006 | 0.0059 |
| Diagnoses - main ICD10: Z47 Other orthopaedic follow-up care | 0 | 0.0042 | 0.2234 | 0.0188 | 0.985 | 0.0026 | 0.0014 | 10.035 | 0.0079 | -0.0054 | 0.005 |
| Diagnoses - main ICD10: Z80 Family history of malignant neop | 0 | 0.1967 | 0.4892 | 0.4021 | 0.6876 | 0.0007 | 0.0014 | 0.9951 | 0.0075 | -0.0026 | 0.0055 |
| Diastolic blood pressure_ automated reading | 0 | -0.1054 | 0.0561 | -18.798 | 0.0601 | 0.1395 | 0.0062 | 10.304 | 0.0182 | 0.0146 | 0.0074 |
| Difference in height between adolescence and adulthood; age 14 | 23449627 | -0.0759 | 0.1579 | -0.481 | 0.6305 | 0.528 | 0.1197 | 0.9759 | 0.0079 | 0.0007 | 0.0058 |
| Difference in height between childhood and adulthood; age 8 | 23449627 | -0.0504 | 0.1341 | -0.3761 | 0.7068 | 0.3393 | 0.0585 | 0.9722 | 0.0089 | 0.0055 | 0.0067 |
| Difficulty not smoking for 1 day | 0 | -0.2575 | 0.1423 | -18.095 | 0.0704 | 0.1068 | 0.0247 | 0.9889 | 0.0089 | 0.0078 | 0.0053 |
| Distance between home and job workplace | 0 | 0.0387 | 0.1724 | 0.2245 | 0.8224 | 0.0113 | 0.0034 | 10.087 | 0.0085 | 0.003 | 0.0064 |
| Doctor diagnosed asthma | 0 | 0.071 | 0.0977 | 0.7266 | 0.4675 | 0.0697 | 0.0106 | 0.9986 | 0.0101 | -0.0091 | 0.0057 |
| Doctor diagnosed hayfever or allergic rhinitis | 0 | 0.0255 | 0.1061 | 0.2407 | 0.8098 | 0.0815 | 0.0107 | 10.017 | 0.0103 | -0.0042 | 0.0062 |
| Drive faster than motorway speed limit | 0 | 0.0314 | 0.0666 | 0.4709 | 0.6377 | 0.0576 | 0.0032 | 1.026 | 0.0108 | -0.0003 | 0.0068 |
| Duration of fitness test | 0 | 0.1652 | 0.231 | 0.7152 | 0.4745 | 0.021 | 0.01 | 1.003 | 0.0077 | -0.0078 | 0.0057 |
| Duration of heavy DIY | 0 | -0.2085 | 0.122 | -17.092 | 0.0874 | 0.0339 | 0.004 | 1.004 | 0.0083 | 0.01 | 0.0063 |
| Duration of light DIY | 0 | -0.0751 | 0.0965 | -0.7782 | 0.4364 | 0.0367 | 0.0037 | 0.9972 | 0.0089 | 0.0069 | 0.0056 |
| Duration of moderate activity | 0 | -0.1398 | 0.0891 | -15.689 | 0.1167 | 0.0318 | 0.0025 | 10.115 | 0.01 | 0.0043 | 0.0061 |
| Duration of other exercises | 0 | -0.07 | 0.1392 | -0.503 | 0.615 | 0.0178 | 0.0033 | 10.121 | 0.0088 | 0.0043 | 0.0057 |
| Duration of strenuous sports | 0 | -0.0727 | 0.2377 | -0.3057 | 0.7598 | 0.0256 | 0.0154 | 0.997 | 0.0087 | 0.0008 | 0.0052 |
| Duration of vigorous activity | 0 | -0.0236 | 0.0954 | -0.2471 | 0.8048 | 0.0361 | 0.0035 | 0.9933 | 0.0085 | -0.0008 | 0.0063 |
| Duration of walks | 0 | 0.0412 | 0.0798 | 0.517 | 0.6052 | 0.0461 | 0.0031 | 10.116 | 0.0105 | 0.0005 | 0.0065 |
| Duration walking for pleasure | 0 | -0.0583 | 0.0923 | -0.6311 | 0.528 | 0.0252 | 0.0026 | 10.145 | 0.0081 | 0.0015 | 0.0061 |
| Eczema | 26482879 | 0.2246 | 0.1435 | 15.657 | 0.1174 | 0.071 | 0.0148 | 10.164 | 0.0075 | 0.0137 | 0.0059 |
| Ever depressed for a whole week | 0 | 0.2217 | 0.1123 | 19.732 | 0.0485 | 0.0607 | 0.0063 | 0.9989 | 0.0094 | -0.0101 | 0.0062 |
| Ever had bowel cancer screening | 0 | -0.0545 | 0.1037 | -0.5253 | 0.5994 | 0.0127 | 0.0017 | 10.204 | 0.0084 | -0.0057 | 0.0058 |
| Ever had hysterectomy (womb removed) | 0 | -0.1167 | 0.1202 | -0.9709 | 0.3316 | 0.0223 | 0.0039 | 10.023 | 0.0093 | 0.0061 | 0.006 |
| Ever had prostate specific antigen (PSA) test | 0 | -0.0511 | 0.1117 | -0.4571 | 0.6476 | 0.0352 | 0.0039 | 10.009 | 0.0086 | -0.0008 | 0.006 |
| Ever had stillbirth_ spontaneous miscarriage or termination | 0 | 0.277 | 0.1235 | 22.431 | 0.0249 | 0.021 | 0.0032 | 0.9961 | 0.0085 | -0.0069 | 0.0055 |
| Ever highly irritable/argumentative for 2 days | 0 | 0.0314 | 0.0973 | 0.323 | 0.7467 | 0.0555 | 0.006 | 0.9921 | 0.009 | -0.0067 | 0.0055 |
| Ever manic/hyper for 2 days | 0 | 0.0423 | 0.1612 | 0.2621 | 0.7932 | 0.0203 | 0.0051 | 0.992 | 0.0081 | -0.0029 | 0.0058 |
| Ever smoked | 0 | -0.0481 | 0.0608 | -0.7917 | 0.4285 | 0.0752 | 0.0037 | 1.02 | 0.0134 | 0.0119 | 0.0063 |
| Ever stopped smoking for 6+ months | 0 | -0.2732 | 0.1896 | -14.405 | 0.1497 | 0.0186 | 0.006 | 10.031 | 0.008 | 0.006 | 0.006 |

| | | | | | | | | | | | |
|---|---|---|---|---|---|---|---|---|---|---|---|
| Ever taken oral contraceptive pill | 0 | 0.2659 | 0.1106 | 24.046 | 0.0162 | 0.0203 | 0.0031 | 10.072 | 0.0078 | -0.0106 | 0.0051 |
| Ever unenthusiastic/disinterested for a whole week | 0 | 0.0904 | 0.1058 | 0.8547 | 0.3927 | 0.0578 | 0.0065 | 0.9998 | 0.0096 | -0.006 | 0.006 |
| Ever used hormone-replacement therapy (HRT) | 0 | -0.1249 | 0.0897 | -13.925 | 0.1638 | 0.0411 | 0.0035 | 0.9959 | 0.0085 | 0.021 | 0.0062 |
| Ever vs never smoked | 20418890 | -0.1438 | 0.0931 | -15.438 | 0.1226 | 0.0718 | 0.0072 | 10.021 | 0.0073 | 0.0107 | 0.0053 |
| Excessive daytime sleepiness | 27992416 | -0.199 | 0.0924 | -21.545 | 0.0312 | 0.0539 | 0.0049 | 10.061 | 0.0086 | 0.0118 | 0.0058 |
| Exposure to tobacco smoke at home | 0 | -0.094 | 0.1181 | -0.7961 | 0.426 | 0.0121 | 0.0017 | 0.9968 | 0.0075 | -0.0022 | 0.0057 |
| Exposure to tobacco smoke outside home | 0 | 0.0528 | 0.0852 | 0.6195 | 0.5356 | 0.0329 | 0.0026 | 10.104 | 0.0098 | -0.0084 | 0.0063 |
| Extreme bmi | 23563607 | 0.1798 | 0.0955 | 18.822 | 0.0598 | 0.6878 | 0.0595 | 10.276 | 0.0125 | -0.0135 | 0.0081 |
| Extreme height | 23563607 | 0.1308 | 0.0787 | 16.609 | 0.0967 | 12.476 | 0.1052 | 10.132 | 0.0189 | 0.0049 | 0.0075 |
| Extreme waist-to-hip ratio | 23563607 | -0.2467 | 0.1517 | -16.268 | 0.1038 | 0.3516 | 0.0653 | 0.977 | 0.0096 | 0.0099 | 0.0066 |
| Eye problems/disorders: Cataract | 0 | 0.2254 | 0.1401 | 16.093 | 0.1075 | 0.0217 | 0.0049 | 10.162 | 0.0083 | -0.0104 | 0.0054 |
| Eye problems/disorders: Diabetes related eye disease | 0 | 0.0784 | 0.176 | 0.4455 | 0.656 | 0.0184 | 0.0047 | 10.095 | 0.008 | -0.0033 | 0.0059 |
| Eye problems/disorders: Glaucoma | 0 | -0.0408 | 0.1171 | -0.3482 | 0.7277 | 0.0385 | 0.0061 | 10.168 | 0.01 | 0.005 | 0.0061 |
| Eye problems/disorders: Injury or trauma resulting in loss of vis | 0 | NA | NA | NA | NA | NA | NA | NA | NA | NA | NA |
| Falls in the last year | 0 | -0.1051 | 0.0792 | -13.263 | 0.1847 | 0.0343 | 0.0022 | 0.9942 | 0.0096 | 0.0034 | 0.0062 |
| Family relationship satisfaction | 0 | 0.0313 | 0.1013 | 0.3094 | 0.757 | 0.0544 | 0.0055 | 10.103 | 0.009 | 0.0041 | 0.0058 |
| Fasting glucose main effect | 22581228 | 0.1163 | 0.1165 | 0.9978 | 0.3184 | 0.0842 | 0.0153 | 0.9997 | 0.0104 | -0.0038 | 0.0058 |
| Fasting insulin main effect | 22581228 | -0.0212 | 0.1291 | -0.1644 | 0.8694 | 0.075 | 0.0111 | 10.075 | 0.0083 | 5,04E-01 | 0.006 |
| Fasting proinsulin | 20081858 | -0.0176 | 0.1934 | -0.0908 | 0.9276 | 0.1617 | 0.0957 | 0.9833 | 0.0125 | 0.0019 | 0.0057 |
| Father still alive | 0 | 0.4961 | 0.1772 | 28.004 | 0.0051 | 0.0068 | 0.0016 | 10.088 | 0.0079 | -0.011 | 0.0058 |
| Fathers age at death | 27015805 | 0.0902 | 0.1298 | 0.6949 | 0.4871 | 0.0423 | 0.0072 | 10.149 | 0.0079 | -0.004 | 0.0061 |
| Fathers age at death | 0 | 0.0402 | 0.0792 | 0.5081 | 0.6114 | 0.0343 | 0.0033 | 10.204 | 0.0097 | -0.0051 | 0.0057 |
| Fed-up feelings | 0 | -0.0541 | 0.0614 | -0.8809 | 0.3784 | 0.0713 | 0.0035 | 10.279 | 0.012 | 0.004 | 0.0066 |
| Femoral Neck bone mineral density | 26367794 | 0.0156 | 0.0893 | 0.1741 | 0.8618 | 0.1206 | 0.0153 | 0.9796 | 0.0097 | -0.0028 | 0.0056 |
| Femoral neck bone mineral density | 22504420 | 0.1287 | 0.0777 | 16.551 | 0.0979 | 0.302 | 0.0264 | 0.9822 | 0.0107 | -0.0126 | 0.0054 |
| Ferritin | 25352340 | 0.0975 | 0.1744 | 0.5591 | 0.5761 | 0.0939 | 0.0295 | 10.269 | 0.0112 | -0.0031 | 0.0073 |
| Financial situation satisfaction | 0 | -0.1547 | 0.0919 | -16.835 | 0.0923 | 0.0582 | 0.0061 | 10.044 | 0.0091 | 0.0095 | 0.0058 |
| Fluid intelligence score | 0 | 0.081 | 0.0728 | 11.135 | 0.2655 | 0.2397 | 0.0111 | 10.363 | 0.013 | -0.0002 | 0.0073 |
| Forced expiratory volume in 1 second (FEV1) | 28166213 | 0.1349 | 0.0708 | 19.064 | 0.0566 | 0.2614 | 0.018 | 0.9693 | 0.0095 | -0.0076 | 0.0057 |
| Forced expiratory volume in 1 second (FEV1) | 26635082 | 0.105 | 0.1081 | 0.9717 | 0.3312 | 0.1403 | 0.018 | 0.9875 | 0.0091 | 0.0001 | 0.0057 |
| Forced expiratory volume in 1 second (FEV1) | 21946350 | 0.1838 | 0.1517 | 12.117 | 0.2256 | 0.1716 | 0.0228 | 0.878 | 0.0135 | -0.0034 | 0.0096 |
| Forced expiratory volume in 1 second (FEV1)/Forced Vital capa | 28166213 | 0.0972 | 0.0792 | 1.228 | 0.2195 | 0.2644 | 0.0224 | 0.9641 | 0.0108 | -0.0086 | 0.0064 |
| Forced expiratory volume in 1 second (FEV1)/Forced Vital capa | 26635082 | 0.0904 | 0.1202 | 0.7523 | 0.4519 | 0.1087 | 0.0149 | 0.9815 | 0.0083 | 0.0002 | 0.0059 |
| Forced expiratory volume in 1 second (FEV1)/Forced Vital capa | 21946350 | 0.1176 | 0.118 | 0.9964 | 0.319 | 0.1187 | 0.0148 | 0.9315 | 0.0091 | -0.0018 | 0.006 |
| Forced expiratory volume in 1-second (FEV1) | 0 | 0.1143 | 0.0527 | 21.677 | 0.0302 | 0.1527 | 0.0071 | 10.182 | 0.0167 | -0.0037 | 0.0078 |
| Forced expiratory volume in 1-second (FEV1)_ Best measure | 0 | 0.1357 | 0.0556 | 24.422 | 0.0146 | 0.1659 | 0.008 | 10.225 | 0.0158 | -0.0069 | 0.0075 |
| Forced expiratory volume in 1-second (FEV1)_ predicted | 0 | 0.1332 | 0.0706 | 18.858 | 0.0593 | 0.1755 | 0.0134 | 10.158 | 0.0142 | -0.0009 | 0.007 |
| Forced expiratory volume in 1-second (FEV1)_ predicted percer | 0 | 0.1251 | 0.0596 | 21.007 | 0.0357 | 0.2051 | 0.0107 | 1.02 | 0.0122 | -0.0079 | 0.0063 |
| Forced vital capacity (FVC) | 0 | 0.0939 | 0.0506 | 18.573 | 0.0633 | 0.1765 | 0.0083 | 1.036 | 0.0195 | 0.0014 | 0.0079 |
| Forced vital capacity (FVC)_ Best measure | 0 | 0.1183 | 0.0529 | 22.364 | 0.0253 | 0.1935 | 0.0091 | 10.336 | 0.0179 | -0.0033 | 0.0078 |
| Forced Vital capacity(FVC) | 28166213 | 0.1253 | 0.0712 | 17.612 | 0.0782 | 0.2605 | 0.0168 | 0.9729 | 0.0097 | -0.0057 | 0.0056 |
| Forced Vital capacity(FVC) | 26635082 | 0.0509 | 0.1009 | 0.5047 | 0.6138 | 0.1519 | 0.0169 | 0.978 | 0.0087 | 0.0008 | 0.0058 |
| Forearm Bone mineral density | 26367794 | 0.2141 | 0.2552 | 0.839 | 0.4015 | 0.0792 | 0.0459 | 10.116 | 0.0079 | -0.0078 | 0.0059 |
| Former alcohol drinker | 0 | -0.0552 | 0.1675 | -0.3293 | 0.7419 | 0.1312 | 0.0245 | 0.9948 | 0.0075 | -0.002 | 0.006 |
| Former vs Current smoker | 20418890 | 0.2785 | 0.1645 | 16.929 | 0.0905 | 0.0579 | 0.0112 | 10.071 | 0.0075 | -0.0059 | 0.0057 |
| Fracture resulting from simple fall | 0 | 0.1597 | 0.1834 | 0.8707 | 0.3839 | 0.0518 | 0.0154 | 0.9945 | 0.0077 | 0.001 | 0.0062 |
| Fractured bone site(s): Ankle | 0 | -0.0082 | 0.1541 | -0.0529 | 0.9578 | 0.006 | 0.0016 | 0.9971 | 0.0077 | 0.0016 | 0.0052 |

| | | | | | | | | | | | |
|---|---|---|---|---|---|---|---|---|---|---|---|
| Fractured bone site(s): Arm | 0 | 0.0354 | 0.1653 | 0.2142 | 0.8304 | 0.0047 | 0.0015 | 0.9977 | 0.0075 | 0.0019 | 0.0057 |
| Fractured bone site(s): Other bones | 0 | 0.1388 | 0.1424 | 0.9744 | 0.3299 | 0.0085 | 0.0018 | 10.036 | 0.0082 | -0.0088 | 0.0065 |
| Fractured bone site(s): Wrist | 0 | -0.1608 | 0.148 | -1.086 | 0.2775 | 0.0068 | 0.0016 | 10.062 | 0.008 | -0.0008 | 0.006 |
| Fractured/broken bones in last 5 years | 0 | 0.0584 | 0.1039 | 0.562 | 0.5741 | 0.0188 | 0.0023 | 10.067 | 0.0097 | -0.0067 | 0.0067 |
| Free cholesterol | 27005778 | 0.1423 | 0.2138 | 0.6654 | 0.5058 | 0.1016 | 0.0438 | 10.112 | 0.0112 | -0.0051 | 0.0061 |
| Free cholesterol in IDL | 27005778 | 0.0402 | 0.1642 | 0.2449 | 0.8065 | 0.1144 | 0.0319 | 0.9941 | 0.0124 | -0.0077 | 0.0061 |
| Free cholesterol in large HDL | 27005778 | -0.0687 | 0.1675 | -0.4102 | 0.6817 | 0.1056 | 0.0286 | 10.072 | 0.0156 | -0.0017 | 0.006 |
| Free cholesterol in large LDL | 27005778 | 0.1233 | 0.1652 | 0.7463 | 0.4555 | 0.1155 | 0.0307 | 0.9868 | 0.0126 | -0.0097 | 0.0061 |
| Free cholesterol in large VLDL | 27005778 | -0.149 | 0.1412 | -10.557 | 0.2911 | 0.1352 | 0.0325 | 0.978 | 0.0077 | 0.0026 | 0.0053 |
| Free cholesterol in medium HDL | 27005778 | -0.1397 | 0.1859 | -0.7514 | 0.4524 | 0.0714 | 0.0259 | 0.9947 | 0.0097 | -0.0019 | 0.0056 |
| Free cholesterol in medium VLDL | 27005778 | -0.0108 | 0.1508 | -0.0715 | 0.943 | 0.1192 | 0.0367 | 0.9871 | 0.0081 | -0.001 | 0.0054 |
| Free cholesterol in small VLDL | 27005778 | -0.0298 | 0.1627 | -0.1832 | 0.8546 | 0.1244 | 0.0351 | 0.9898 | 0.0089 | -0.0033 | 0.006 |
| Free cholesterol in very large HDL | 27005778 | 0.0457 | 0.224 | 0.2039 | 0.8384 | 0.0541 | 0.0265 | 10.125 | 0.0154 | -0.0049 | 0.0058 |
| Free cholesterol to esterified cholesterol ratio | 27005778 | 0.1306 | 0.2208 | 0.5914 | 0.5543 | 0.1048 | 0.0466 | 0.9964 | 0.01 | -0.0078 | 0.0062 |
| Frequency of depressed mood in last 2 weeks | 0 | 0.0761 | 0.0742 | 10.257 | 0.305 | 0.0436 | 0.0029 | 10.044 | 0.0101 | -0.0027 | 0.0061 |
| Frequency of heavy DIY in last 4 weeks | 0 | -0.0093 | 0.1534 | -0.0609 | 0.9515 | 0.0206 | 0.0036 | 10.053 | 0.0076 | 0.003 | 0.0061 |
| Frequency of light DIY in last 4 weeks | 0 | 0.1398 | 0.119 | 11.747 | 0.2401 | 0.0224 | 0.0032 | 10.041 | 0.0074 | -0.0053 | 0.0057 |
| Frequency of other exercises in last 4 weeks | 0 | -0.0485 | 0.1163 | -0.4173 | 0.6765 | 0.0214 | 0.0036 | 0.9981 | 0.0083 | 0.0065 | 0.0058 |
| Frequency of stair climbing in last 4 weeks | 0 | 0.0224 | 0.0782 | 0.2864 | 0.7746 | 0.035 | 0.0024 | 10.138 | 0.0099 | -0.0026 | 0.0063 |
| Frequency of tenseness / restlessness in last 2 weeks | 0 | 0.0568 | 0.075 | 0.7574 | 0.4488 | 0.0432 | 0.0027 | 10.199 | 0.0101 | -0.0011 | 0.0059 |
| Frequency of tiredness / lethargy in last 2 weeks | 0 | 0.024 | 0.0669 | 0.3582 | 0.7202 | 0.0602 | 0.0032 | 10.206 | 0.0108 | 3,19E-01 | 0.0069 |
| Frequency of travelling from home to job workplace | 0 | 0.0698 | 0.1059 | 0.6594 | 0.5096 | 0.0225 | 0.0032 | 10.176 | 0.0086 | -0.0046 | 0.0055 |
| Frequency of unenthusiasm / disinterest in last 2 weeks | 0 | 0.028 | 0.0702 | 0.3988 | 0.6901 | 0.0396 | 0.0028 | 10.137 | 0.0103 | -0.0051 | 0.0061 |
| Frequency of walking for pleasure in last 4 weeks | 0 | 0.1144 | 0.0843 | 13.572 | 0.1747 | 0.0348 | 0.0027 | 10.074 | 0.0089 | -0.0001 | 0.0064 |
| Friendships satisfaction | 0 | -0.0095 | 0.1033 | -0.0916 | 0.927 | 0.0584 | 0.006 | 10.035 | 0.0087 | 0.0036 | 0.0063 |
| Getting up in morning | 0 | 0.0308 | 0.0605 | 0.5091 | 0.6107 | 0.0698 | 0.0034 | 10.315 | 0.0135 | 0.0015 | 0.0058 |
| Glucose | 27005778 | -0.2086 | 0.1764 | -11.825 | 0.237 | 0.0815 | 0.0219 | 0.9922 | 0.0076 | 0.0063 | 0.0059 |
| Glutamine | 27005778 | 0.161 | 0.2033 | 0.792 | 0.4284 | 0.0562 | 0.0228 | 10.211 | 0.0098 | -0.0032 | 0.0055 |
| Glycoprotein acetyls; mainly a1-acid glycoprotein | 27005778 | -0.3094 | 0.1572 | -19.675 | 0.0491 | 0.1149 | 0.0312 | 0.98 | 0.0074 | 0.0046 | 0.0054 |
| Guilty feelings | 0 | 0.0038 | 0.0657 | 0.0573 | 0.9543 | 0.0513 | 0.0028 | 10.102 | 0.0103 | -0.0016 | 0.0065 |
| Had major operations | 0 | -0.0905 | 0.1171 | -0.7732 | 0.4394 | 0.0249 | 0.0039 | 10.053 | 0.0083 | 0.0037 | 0.0055 |
| Had menopause | 0 | 0.0023 | 0.1307 | 0.0179 | 0.9857 | 0.0201 | 0.0045 | 10.032 | 0.0087 | 0.0044 | 0.006 |
| Had other major operations | 0 | 0.088 | 0.1043 | 0.8445 | 0.3984 | 0.0292 | 0.0032 | 10.146 | 0.008 | -0.0048 | 0.006 |
| Hair/balding pattern: Pattern 2 | 0 | 0.0483 | 0.086 | 0.5622 | 0.574 | 0.0515 | 0.0073 | 1.042 | 0.0179 | 0.0032 | 0.0063 |
| Hair/balding pattern: Pattern 3 | 0 | -0.157 | 0.0947 | -16.569 | 0.0975 | 0.0579 | 0.0165 | 10.152 | 0.0269 | 0.0032 | 0.0066 |
| Hair/balding pattern: Pattern 4 | 0 | -0.014 | 0.063 | -0.2222 | 0.8242 | 0.1627 | 0.0227 | 1.108 | 0.0347 | -0.0006 | 0.0075 |
| Hand grip strength (left) | 0 | 0.0989 | 0.0569 | 17.388 | 0.0821 | 0.1076 | 0.0043 | 10.241 | 0.014 | -0.006 | 0.0079 |
| Hand grip strength (right) | 0 | 0.0897 | 0.0587 | 1.527 | 0.1268 | 0.1067 | 0.0045 | 10.304 | 0.0147 | 0.0022 | 0.0082 |
| Handedness (chirality/laterality): Left-handed | 0 | 0.0484 | 0.135 | 0.3586 | 0.7199 | 0.0095 | 0.0018 | 10.068 | 0.0087 | -0.0074 | 0.0059 |
| Handedness (chirality/laterality): Use both right and left hands e | 0 | -0.3627 | 0.1994 | -18.191 | 0.0689 | 0.0049 | 0.0014 | 10.005 | 0.0074 | 0.007 | 0.0055 |
| Happiness | 0 | 0.0612 | 0.0904 | 0.6764 | 0.4988 | 0.0602 | 0.0055 | 10.091 | 0.0085 | -0.0055 | 0.0062 |
| HbA1C | 20858683 | -0.0535 | 0.1249 | -0.4285 | 0.6683 | 0.0628 | 0.0124 | 0.9955 | 0.0085 | 0.0024 | 0.0053 |
| HDL cholesterol | 20686565 | -0.0417 | 0.08 | -0.5212 | 0.6023 | 0.1287 | 0.0243 | 10.407 | 0.0516 | 0.0074 | 0.0065 |
| Headaches for 3+ months | 0 | 0.0186 | 0.13 | 0.1428 | 0.8864 | 0.0582 | 0.0082 | 10.041 | 0.0075 | 0.0037 | 0.0058 |
| Health satisfaction | 0 | -0.008 | 0.0872 | -0.0923 | 0.9265 | 0.0831 | 0.0056 | 10.051 | 0.009 | 0.0084 | 0.0064 |
| Hearing aid user | 0 | -0.2122 | 0.124 | -17.107 | 0.0871 | 0.0187 | 0.003 | 0.995 | 0.0085 | 0.0103 | 0.0062 |
| Hearing difficulty/problems with background noise | 0 | -0.0148 | 0.0707 | -0.2087 | 0.8347 | 0.0535 | 0.0028 | 10.085 | 0.0098 | 0.0066 | 0.0065 |

| | | | | | | | | | | | |
|---|---|---|---|---|---|---|---|---|---|---|---|
| Hearing difficulty/problems: Yes | 0 | -0.0246 | 0.0724 | -0.34 | 0.7339 | 0.0432 | 0.0026 | 10.013 | 0.0098 | 0.0071 | 0.0063 |
| Heart rate | 23583979 | 0.0848 | 0.0868 | 0.9773 | 0.3284 | 0.0835 | 0.0111 | 10.103 | 0.0099 | -0.0035 | 0.0054 |
| Heel bone mineral density (BMD) T-score_ automated | 0 | 0.0503 | 0.0453 | 11.116 | 0.2663 | 0.3015 | 0.0379 | 10.638 | 0.0455 | -0.0034 | 0.0079 |
| Heel bone mineral density (BMD) T-score_ automated (left) | 0 | -0.0048 | 0.0528 | -0.0905 | 0.9279 | 0.3152 | 0.0402 | 10.058 | 0.0289 | 0.0065 | 0.0068 |
| Heel bone mineral density (BMD) T-score_ automated (right) | 0 | 0.0344 | 0.0549 | 0.6276 | 0.5303 | 0.315 | 0.0407 | 10.071 | 0.03 | 0.0016 | 0.0071 |
| Height; Females at age 10 and males at age 12 | 23449627 | 0.1673 | 0.0972 | 17.219 | 0.0851 | 0.4286 | 0.0481 | 0.9487 | 0.0089 | -0.0018 | 0.0058 |
| Height_2010 | 20881960 | 0.1365 | 0.0609 | 22.408 | 0.025 | 0.286 | 0.018 | 0.9862 | 0.0199 | -0.0055 | 0.0078 |
| Hip circumference | 25673412 | 0.1164 | 0.0665 | 17.497 | 0.0802 | 0.1296 | 0.0062 | 0.8492 | 0.0095 | -0.0065 | 0.0059 |
| Hip circumference | 0 | 0.0274 | 0.0496 | 0.552 | 0.5809 | 0.2167 | 0.0093 | 10.543 | 0.023 | 0.0014 | 0.0081 |
| HOMA-B | 20081858 | -0.1119 | 0.1402 | -0.798 | 0.4249 | 0.0783 | 0.0135 | 0.9946 | 0.0082 | -0.0002 | 0.0061 |
| HOMA-IR | 20081858 | -0.0992 | 0.1524 | -0.6509 | 0.5151 | 0.072 | 0.0138 | 0.9998 | 0.0074 | 0.0003 | 0.006 |
| Home area population density - urban or rural: England/Wales - | 0 | -0.1751 | 0.2519 | -0.6952 | 0.4869 | 0.0019 | 0.0015 | 10.272 | 0.0073 | 0.0032 | 0.0052 |
| Home area population density - urban or rural: Scotland - Large | 0 | -0.038 | 0.1773 | -0.2144 | 0.8302 | 0.0113 | 0.0034 | 21.341 | 0.0176 | 0.0074 | 0.0088 |
| ICV | 25607358 | 0.1564 | 0.1578 | 0.9915 | 0.3214 | 0.1991 | 0.0482 | 0.9932 | 0.0081 | -0.0029 | 0.0055 |
| Illness_ injury_ bereavement_ stress in last 2 years: Death of a c | 0 | 0.0371 | 0.1609 | 0.2303 | 0.8178 | 0.0059 | 0.0017 | 10.092 | 0.0082 | -0.002 | 0.0064 |
| Illness_ injury_ bereavement_ stress in last 2 years: Financial di | 0 | 0.1324 | 0.0661 | 20.037 | 0.0451 | 0.0329 | 0.0024 | 10.307 | 0.01 | -0.0061 | 0.0056 |
| Illness_ injury_ bereavement_ stress in last 2 years: Marital sepa | 0 | 0.2488 | 0.1613 | 15.425 | 0.1229 | 0.0043 | 0.0015 | 10.102 | 0.0079 | -0.0052 | 0.0052 |
| Illness_ injury_ bereavement_ stress in last 2 years: None of the | 0 | -0.1937 | 0.0829 | -23.352 | 0.0195 | 0.0223 | 0.0019 | 1.018 | 0.008 | 0.0107 | 0.0059 |
| Illness_ injury_ bereavement_ stress in last 2 years: Serious illn | 0 | 0.0787 | 0.1337 | 0.5886 | 0.5562 | 0.0103 | 0.0017 | 10.132 | 0.0078 | 0.0035 | 0.006 |
| Illness_ injury_ bereavement_ stress in last 2 years: Serious illn | 0 | 0.1738 | 0.101 | 17.215 | 0.0852 | 0.0125 | 0.0018 | 10.227 | 0.009 | -0.008 | 0.0055 |
| Illnesses of father: Chronic bronchitis/emphysema | 0 | -0.2731 | 0.1048 | -26.062 | 0.0092 | 0.0153 | 0.002 | 1.02 | 0.0082 | 0.0133 | 0.0054 |
| Illnesses of father: Diabetes | 0 | -0.0833 | 0.0981 | -0.8498 | 0.3954 | 0.0185 | 0.0025 | 10.219 | 0.0099 | 0.0012 | 0.0056 |
| Illnesses of father: Heart disease | 0 | 0.0433 | 0.0843 | 0.514 | 0.6073 | 0.0313 | 0.0027 | 0.9896 | 0.0096 | -0.008 | 0.006 |
| Illnesses of father: High blood pressure | 0 | -0.0961 | 0.1113 | -0.8641 | 0.3875 | 0.0202 | 0.0022 | 10.072 | 0.0094 | 0.0067 | 0.0062 |
| Illnesses of father: Lung cancer | 0 | 0.0681 | 0.1154 | 0.5905 | 0.5549 | 0.0156 | 0.0021 | 0.988 | 0.0081 | -0.0037 | 0.0064 |
| Illnesses of father: None of the above (group 1) | 0 | 0.0527 | 0.1041 | 0.5058 | 0.613 | 0.0154 | 0.0021 | 10.164 | 0.009 | -0.0015 | 0.0056 |
| Illnesses of father: None of the above (group 2) | 0 | 0.0086 | 0.1284 | 0.0668 | 0.9467 | 0.0106 | 0.0018 | 0.99 | 0.0084 | -0.0068 | 0.0058 |
| Illnesses of father: Prostate cancer | 0 | 0.1187 | 0.1499 | 0.7918 | 0.4285 | 0.0091 | 0.0019 | 10.066 | 0.0097 | -0.0042 | 0.006 |
| Illnesses of mother: Alzheimers disease/dementia | 0 | 0.0169 | 0.151 | 0.1117 | 0.9111 | 0.0072 | 0.0018 | 10.109 | 0.0103 | 0.0031 | 0.006 |
| Illnesses of mother: Breast cancer | 0 | -0.0437 | 0.1304 | -0.3352 | 0.7375 | 0.0098 | 0.002 | 10.027 | 0.0097 | -0.0005 | 0.006 |
| Illnesses of mother: Chronic bronchitis/emphysema | 0 | -0.1456 | 0.1239 | -11.752 | 0.2399 | 0.0114 | 0.002 | 10.095 | 0.0077 | 0.0035 | 0.0059 |
| Illnesses of mother: Diabetes | 0 | -0.0993 | 0.1013 | -0.9806 | 0.3268 | 0.0208 | 0.0021 | 10.081 | 0.0099 | 0.0071 | 0.0064 |
| Illnesses of mother: Heart disease | 0 | -0.0055 | 0.1151 | -0.0479 | 0.9618 | 0.0158 | 0.002 | 10.071 | 0.0085 | 0.0048 | 0.0059 |
| Illnesses of mother: High blood pressure | 0 | -0.1145 | 0.085 | -13.479 | 0.1777 | 0.0301 | 0.0025 | 10.089 | 0.0101 | 0.0075 | 0.0063 |
| Illnesses of mother: None of the above (group 1) | 0 | 0.2103 | 0.1106 | 19.008 | 0.0573 | 0.0171 | 0.0019 | 10.077 | 0.0085 | -0.0099 | 0.0059 |
| Illnesses of mother: None of the above (group 2) | 0 | -0.0426 | 0.1326 | -0.3214 | 0.7479 | 0.0088 | 0.0019 | 10.063 | 0.0086 | 0.0014 | 0.0059 |
| Illnesses of siblings: Diabetes | 0 | -0.0009 | 0.0939 | -0.0091 | 0.9927 | 0.0212 | 0.0024 | 1.002 | 0.0086 | 0.0004 | 0.0056 |
| Illnesses of siblings: Heart disease | 0 | -0.1174 | 0.1178 | -0.9965 | 0.319 | 0.0165 | 0.0024 | 10.132 | 0.0088 | 0.0049 | 0.0058 |
| Illnesses of siblings: High blood pressure | 0 | 0.0298 | 0.0836 | 0.3566 | 0.7214 | 0.0372 | 0.0029 | 0.9968 | 0.0101 | -0.0054 | 0.0063 |
| Illnesses of siblings: None of the above (group 1) | 0 | 0.0413 | 0.0857 | 0.4822 | 0.6297 | 0.0367 | 0.003 | 10.085 | 0.0096 | -0.0013 | 0.0063 |
| Illnesses of siblings: None of the above (group 2) | 0 | -0.0057 | 0.1441 | -0.0396 | 0.9684 | 0.0102 | 0.002 | 0.9968 | 0.008 | -0.01 | 0.0056 |
| Illnesses of siblings: Severe depression | 0 | 0.1322 | 0.1331 | 0.9931 | 0.3207 | 0.0137 | 0.0021 | 10.078 | 0.0081 | -0.0025 | 0.0063 |
| Illnesses of siblings: Stroke | 0 | -0.073 | 0.2028 | -0.3601 | 0.7188 | 0.0045 | 0.002 | 10.062 | 0.0075 | 0.0001 | 0.0061 |
| Impedance of arm (left) | 0 | 0.0167 | 0.0531 | 0.3149 | 0.7528 | 0.2328 | 0.0096 | 1.06 | 0.0253 | -0.0057 | 0.0084 |
| Impedance of arm (right) | 0 | 0.0159 | 0.0531 | 0.2985 | 0.7653 | 0.2369 | 0.0095 | 10.393 | 0.0249 | -0.0049 | 0.0086 |
| Impedance of leg (left) | 0 | -0.0299 | 0.0517 | -0.5782 | 0.5632 | 0.2453 | 0.01 | 10.239 | 0.0232 | 0.0027 | 0.0078 |
| Impedance of leg (right) | 0 | -0.0385 | 0.0523 | -0.7357 | 0.4619 | 0.2442 | 0.0098 | 10.334 | 0.0225 | 0.0037 | 0.0079 |

| | | | | | | | | | | | |
|---|---|---|---|---|---|---|---|---|---|---|---|
| Impedance of whole body | 0 | -0.0088 | 0.0511 | -0.1714 | 0.8639 | 0.2603 | 0.0107 | 10.359 | 0.025 | -0.0008 | 0.0083 |
| Infant head circumference | 22504419 | -0.1811 | 0.1491 | -12.143 | 0.2246 | 0.2413 | 0.0475 | 0.9878 | 0.008 | 0.0046 | 0.006 |
| Inflammatory Bowel Disease (Euro) | 26192919 | 0.0257 | 0.0701 | 0.3668 | 0.7138 | 0.3263 | 0.0334 | 10.428 | 0.0129 | 0.0586 | 0.0059 |
| Insomnia | 28604731 | -0.0081 | 0.0995 | -0.0811 | 0.9354 | 0.0472 | 0.0051 | 10.068 | 0.0082 | 0.0031 | 0.0053 |
| Insomnia | 27992416 | -0.0867 | 0.0904 | -0.9594 | 0.3374 | 0.1371 | 0.0125 | 10.001 | 0.0092 | 0.0044 | 0.0055 |
| Intelligence | 28530673 | 0.061 | 0.0734 | 0.8303 | 0.4064 | 0.189 | 0.0112 | 10.095 | 0.0096 | 0.0007 | 0.006 |
| Irritability | 0 | -0.0368 | 0.0633 | -0.5811 | 0.5612 | 0.0691 | 0.0042 | 0.9895 | 0.014 | 0.0036 | 0.0067 |
| Ischemic stroke | 26935894 | NA | NA | NA | NA | NA | NA | NA | NA | NA | NA |
| Isoleucine | 27005778 | -0.1781 | 0.1733 | -10.276 | 0.3041 | 0.0797 | 0.0261 | 0.987 | 0.008 | 0.0021 | 0.0056 |
| Job involves heavy manual or physical work | 0 | -0.0838 | 0.0713 | -11.752 | 0.2399 | 0.0857 | 0.0049 | 1.039 | 0.0108 | 0.0065 | 0.0067 |
| Job involves mainly walking or standing | 0 | -0.081 | 0.0772 | -10.492 | 0.2941 | 0.0801 | 0.0049 | 10.301 | 0.011 | 0.0064 | 0.0069 |
| Job involves shift work | 0 | -0.0103 | 0.0878 | -0.1177 | 0.9063 | 0.0297 | 0.0032 | 10.012 | 0.0083 | 0.0031 | 0.005 |
| Knee pain for 3+ months | 0 | -0.1911 | 0.1812 | -10.547 | 0.2916 | 0.029 | 0.007 | 0.994 | 0.0077 | 0.0071 | 0.0062 |
| LDL cholesterol | 20686565 | -0.0469 | 0.0817 | -0.5737 | 0.5661 | 0.1231 | 0.0184 | 0.9991 | 0.0207 | -0.0011 | 0.0069 |
| Leg fat mass (left) | 0 | 0.0042 | 0.0477 | 0.0885 | 0.9295 | 0.2346 | 0.0084 | 10.393 | 0.021 | 0.0045 | 0.008 |
| Leg fat mass (right) | 0 | -0.0012 | 0.0475 | -0.0243 | 0.9806 | 0.2338 | 0.0083 | 10.368 | 0.0209 | 0.0052 | 0.0081 |
| Leg fat percentage (left) | 0 | -0.0348 | 0.0498 | -0.6994 | 0.4843 | 0.21 | 0.0069 | 10.586 | 0.0192 | 0.0044 | 0.0086 |
| Leg fat percentage (right) | 0 | -0.042 | 0.0493 | -0.8526 | 0.3939 | 0.2107 | 0.0069 | 10.514 | 0.0189 | 0.0044 | 0.0088 |
| Leg fat-free mass (left) | 0 | 0.0666 | 0.0495 | 13.459 | 0.1783 | 0.2701 | 0.0118 | 10.432 | 0.0311 | 0.0048 | 0.008 |
| Leg fat-free mass (right) | 0 | 0.0732 | 0.05 | 14.633 | 0.1434 | 0.2678 | 0.0117 | 10.493 | 0.0312 | 0.0043 | 0.0081 |
| Leg pain on walking | 0 | -0.0486 | 0.0954 | -0.5095 | 0.6104 | 0.055 | 0.0057 | 1.012 | 0.0089 | 0.0008 | 0.0057 |
| Leg predicted mass (left) | 0 | 0.0671 | 0.0493 | 13.593 | 0.174 | 0.2695 | 0.0117 | 10.441 | 0.031 | 0.0051 | 0.008 |
| Leg predicted mass (right) | 0 | 0.0746 | 0.05 | 14.906 | 0.1361 | 0.2675 | 0.0117 | 1.049 | 0.0311 | 0.0041 | 0.0081 |
| Length of menstrual cycle | 0 | -0.1442 | 0.1387 | -10.398 | 0.2984 | 0.1135 | 0.0221 | 0.9948 | 0.0082 | 0.0015 | 0.0061 |
| Length of working week for main job | 0 | 0.1463 | 0.1059 | 1.382 | 0.167 | 0.0203 | 0.003 | 10.073 | 0.0076 | -0.0007 | 0.0052 |
| Leptin_adjBMI | 26833098 | -0.0424 | 0.1333 | -0.3179 | 0.7505 | 0.0932 | 0.0189 | 1.002 | 0.0087 | 0.0032 | 0.0056 |
| Leptin_not_adjBMI | 26833098 | 0.111 | 0.1266 | 0.8768 | 0.3806 | 0.1114 | 0.0155 | 0.9847 | 0.0078 | -0.0023 | 0.0056 |
| Leucine | 27005778 | -0.0766 | 0.206 | -0.3716 | 0.7102 | 0.0567 | 0.023 | 1.001 | 0.0083 | 0.0014 | 0.0055 |
| Light smokers_ at least 100 smokes in lifetime | 0 | -0.007 | 0.0949 | -0.0736 | 0.9414 | 0.0733 | 0.0085 | 10.094 | 0.009 | 0.0048 | 0.0059 |
| Loneliness_ isolation | 0 | 0.0225 | 0.072 | 0.313 | 0.7543 | 0.0386 | 0.0025 | 10.196 | 0.0091 | -0.0034 | 0.006 |
| Long-standing illness_ disability or infirmity | 0 | -0.0199 | 0.0687 | -0.2893 | 0.7724 | 0.0516 | 0.0026 | 10.163 | 0.0106 | 1,63E-02 | 0.0065 |
| Loud music exposure frequency | 0 | -0.1105 | 0.1387 | -0.7964 | 0.4258 | 0.0274 | 0.005 | 10.132 | 0.0083 | 0.0053 | 0.0062 |
| Lumbar Spine bone mineral density | 26367794 | 0.0344 | 0.1027 | 0.3351 | 0.7376 | 0.1202 | 0.0153 | 0.9844 | 0.01 | -0.0024 | 0.0055 |
| Lumbar spine bone mineral density | 22504420 | 0.1254 | 0.0828 | 15.142 | 0.13 | 0.2712 | 0.0253 | 10.114 | 0.0118 | -0.009 | 0.006 |
| Lung adenocarcinoma | 27488534 | -0.016 | 0.1651 | -0.0966 | 0.9231 | 0.0308 | 0.012 | 10.189 | 0.0076 | 0.0193 | 0.0049 |
| Lung cancer | 27488534 | 0.0753 | 0.0991 | 0.7597 | 0.4474 | 0.3329 | 0.067 | 10.075 | 0.0097 | 0.0184 | 0.0055 |
| Lung cancer (all) | 24880342 | 0.1242 | 0.109 | 11.393 | 0.2546 | 0.1369 | 0.0299 | 0.9989 | 0.0094 | 0.0011 | 0.0051 |
| Lung cancer (squamous cell) | 24880342 | 0.0033 | 0.1773 | 0.0184 | 0.9853 | 0.0499 | 0.0197 | 1.005 | 0.008 | -0.0007 | 0.0054 |
| Major depressive disorder | 22472876 | 0.0369 | 0.137 | 0.2694 | 0.7876 | 0.1517 | 0.0263 | 10.144 | 0.0076 | 0.0302 | 0.0058 |
| Maternal smoking around birth | 0 | 0.0711 | 0.0704 | 10.103 | 0.3123 | 0.0513 | 0.0031 | 10.263 | 0.0109 | -0.0017 | 0.006 |
| Maximum heart rate during fitness test | 0 | -0.2286 | 0.1266 | -18.059 | 0.0709 | 0.0683 | 0.0135 | 10.142 | 0.0094 | 0.0116 | 0.0058 |
| Maximum workload during fitness test | 0 | 0.0397 | 0.1651 | 0.2407 | 0.8098 | 0.0457 | 0.0111 | 1.004 | 0.008 | 0.001 | 0.0067 |
| Mean Accumbens | 25607358 | -0.014 | 0.2041 | -0.0685 | 0.9454 | 0.0837 | 0.0401 | 0.9816 | 0.0079 | -0.0008 | 0.0054 |
| Mean Caudate | 25607358 | -0.1999 | 0.1429 | -13.989 | 0.1618 | 0.2455 | 0.0385 | 0.9692 | 0.0075 | 0.0046 | 0.0061 |
| Mean diameter for HDL particles | 27005778 | -0.1444 | 0.1791 | -0.8061 | 0.4202 | 0.1156 | 0.0335 | 10.089 | 0.0209 | 0.0005 | 0.0062 |
| Mean diameter for LDL particles | 27005778 | 0.068 | 0.2566 | 0.265 | 0.791 | 0.0438 | 0.0265 | 0.9976 | 0.0126 | 0.0005 | 0.0058 |
| Mean diameter for VLDL particles | 27005778 | 0.0107 | 0.1493 | 0.0713 | 0.9431 | 0.1298 | 0.0351 | 0.9856 | 0.0094 | 0.0042 | 0.0051 |

| | | | | | | | | | | | |
|---|---|---|---|---|---|---|---|---|---|---|---|
| Mean Hippocampus | 25607358 | -0.2926 | 0.1991 | -14.695 | 0.1417 | 0.1397 | 0.0448 | 0.9904 | 0.0086 | 0.0125 | 0.0064 |
| Mean Pallidum | 25607358 | -0.0188 | 0.1802 | -0.1043 | 0.9169 | 0.1678 | 0.0452 | 0.9764 | 0.008 | 0.0004 | 0.0064 |
| Mean platelet volume | 22139419 | -0.1115 | 0.1014 | -10.992 | 0.2717 | 0.3276 | 0.0546 | 0.9734 | 0.0124 | 0.0053 | 0.0059 |
| Mean Putamen | 25607358 | -0.1805 | 0.1249 | -14.451 | 0.1484 | 0.2867 | 0.0494 | 0.9513 | 0.0088 | 0.0058 | 0.0061 |
| Mean Thalamus | 25607358 | -0.1457 | 0.1826 | -0.7978 | 0.425 | 0.1281 | 0.038 | 0.9844 | 0.0076 | 0.0086 | 0.006 |
| Mean time to correctly identify matches | 0 | 0.0796 | 0.0601 | 13.232 | 0.1858 | 0.0732 | 0.0032 | 1.031 | 0.011 | 0.0029 | 0.0066 |
| Medication for cholesterol_ blood pressure or diabetes: Blood p | 0 | -0.0999 | 0.0719 | -13.892 | 0.1648 | 0.106 | 0.0061 | 10.218 | 0.0118 | 0.0117 | 0.0066 |
| Medication for cholesterol_ blood pressure or diabetes: Cholest | 0 | -0.0449 | 0.0728 | -0.6164 | 0.5376 | 0.0736 | 0.0061 | 10.089 | 0.0113 | -0.0007 | 0.0061 |
| Medication for cholesterol_ blood pressure or diabetes: Insulin | 0 | 0.0537 | 0.1736 | 0.3097 | 0.7568 | 0.0093 | 0.0035 | 10.078 | 0.0086 | -0.0081 | 0.0052 |
| Medication for cholesterol_ blood pressure or diabetes: None of | 0 | 0.0738 | 0.0707 | 10.444 | 0.2963 | 0.0903 | 0.0057 | 10.233 | 0.0117 | -0.0053 | 0.0062 |
| Medication for cholesterol_ blood pressure_ diabetes_ or take e | 0 | -0.0667 | 0.065 | -1.027 | 0.3044 | 0.1067 | 0.0072 | 1.013 | 0.0134 | 0.003 | 0.0068 |
| Medication for cholesterol_ blood pressure_ diabetes_ or take e | 0 | 0.0376 | 0.0871 | 0.4313 | 0.6662 | 0.0543 | 0.005 | 10.007 | 0.0102 | -0.0073 | 0.006 |
| Medication for cholesterol_ blood pressure_ diabetes_ or take e | 0 | -0.1305 | 0.1431 | -0.9116 | 0.362 | 0.0152 | 0.0029 | 10.004 | 0.008 | 0.0117 | 0.0055 |
| Medication for cholesterol_ blood pressure_ diabetes_ or take e | 0 | 0.0534 | 0.0788 | 0.6782 | 0.4976 | 0.0686 | 0.005 | 10.003 | 0.0106 | -0.0033 | 0.0065 |
| Medication for pain relief_ constipation_ heartburn: Aspirin | 0 | 0.0062 | 0.0802 | 0.0772 | 0.9384 | 0.0221 | 0.0019 | 1.006 | 0.008 | 0.0016 | 0.0058 |
| Medication for pain relief_ constipation_ heartburn: Ibuprofen ( | 0 | -0.052 | 0.0868 | -0.5992 | 0.5491 | 0.0228 | 0.0019 | 0.9951 | 0.0084 | 0.0008 | 0.0058 |
| Medication for pain relief_ constipation_ heartburn: Laxatives ( | 0 | 0.0053 | 0.1125 | 0.0471 | 0.9624 | 0.0115 | 0.0017 | 1.005 | 0.0079 | -0.0031 | 0.0056 |
| Medication for pain relief_ constipation_ heartburn: None of the | 0 | -0.0005 | 0.0687 | -0.0069 | 0.9945 | 0.055 | 0.0029 | 10.199 | 0.0116 | 0.0024 | 0.0065 |
| Medication for pain relief_ constipation_ heartburn: Omeprazol | 0 | -0.0628 | 0.1074 | -0.585 | 0.5585 | 0.018 | 0.0016 | 10.214 | 0.0085 | 0.0061 | 0.0066 |
| Medication for pain relief_ constipation_ heartburn: Paracetamo | 0 | 0.0085 | 0.0818 | 0.104 | 0.9171 | 0.0431 | 0.0024 | 10.222 | 0.0091 | -0.002 | 0.007 |
| Mineral and other dietary supplements: Calcium | 0 | -0.0151 | 0.123 | -0.1229 | 0.9022 | 0.0123 | 0.0016 | 10.007 | 0.0076 | 0.0005 | 0.0062 |
| Mineral and other dietary supplements: Fish oil (including cod l | 0 | -0.0978 | 0.0787 | -12.435 | 0.2137 | 0.0269 | 0.002 | 10.069 | 0.0091 | 0.0062 | 0.0063 |
| Mineral and other dietary supplements: Glucosamine | 0 | -0.0005 | 0.081 | -0.006 | 0.9952 | 0.024 | 0.0022 | 10.111 | 0.0091 | 0.0034 | 0.006 |
| Mineral and other dietary supplements: None of the above | 0 | 0.0215 | 0.0801 | 0.2681 | 0.7886 | 0.0298 | 0.0023 | 10.122 | 0.0095 | -0.0044 | 0.0064 |
| Mineral and other dietary supplements: Selenium | 0 | -0.003 | 0.1514 | -0.0199 | 0.9841 | 0.0056 | 0.0016 | 10.059 | 0.0085 | 0.0037 | 0.0055 |
| Mineral and other dietary supplements: Zinc | 0 | -0.1534 | 0.1208 | -1.27 | 0.2041 | 0.0109 | 0.0017 | 0.9916 | 0.0079 | 0.0152 | 0.0055 |
| Miserableness | 0 | -0.0198 | 0.0642 | -0.3077 | 0.7583 | 0.0654 | 0.0037 | 10.112 | 0.0122 | 0.0048 | 0.0062 |
| Mono-unsaturated fatty acids | 27005778 | 0.1154 | 0.2096 | 0.5508 | 0.5818 | 0.1131 | 0.043 | 0.985 | 0.0083 | -0.0064 | 0.0058 |
| Mood swings | 0 | -0.0684 | 0.063 | -10.862 | 0.2774 | 0.0724 | 0.0035 | 10.128 | 0.0115 | 0.0047 | 0.0066 |
| Morning/evening person (chronotype) | 0 | -0.0218 | 0.0566 | -0.3845 | 0.7006 | 0.1161 | 0.0046 | 10.424 | 0.015 | -0.0012 | 0.0073 |
| Most recent bowel cancer screening | 0 | 0.161 | 0.2712 | 0.5936 | 0.5528 | 0.0091 | 0.0064 | 10.127 | 0.0078 | 0.0018 | 0.0059 |
| Mothers age at death | 27015805 | -0.1431 | 0.1329 | -10.764 | 0.2817 | 0.0419 | 0.0078 | 10.019 | 0.0083 | 0.0009 | 0.0054 |
| Mothers age at death | 0 | 0.0551 | 0.1077 | 0.5118 | 0.6088 | 0.0254 | 0.0031 | 10.218 | 0.0084 | -0.0036 | 0.0057 |
| Mouth/teeth dental problems: Bleeding gums | 0 | -0.0649 | 0.0874 | -0.7422 | 0.4579 | 0.0225 | 0.0021 | 10.068 | 0.0096 | 0.0043 | 0.0056 |
| Mouth/teeth dental problems: Dentures | 0 | -0.0697 | 0.0701 | -0.9943 | 0.3201 | 0.0483 | 0.0031 | 10.304 | 0.012 | 0.0057 | 0.0061 |
| Mouth/teeth dental problems: Loose teeth | 0 | -0.0173 | 0.0962 | -0.1801 | 0.8571 | 0.0137 | 0.0018 | 10.123 | 0.0083 | 0.0053 | 0.0053 |
| Mouth/teeth dental problems: Mouth ulcers | 0 | -0.1491 | 0.0752 | -19.819 | 0.0475 | 0.0303 | 0.0036 | 10.168 | 0.011 | 0.0134 | 0.006 |
| Mouth/teeth dental problems: None of the above | 0 | 0.0315 | 0.0735 | 0.4289 | 0.668 | 0.0367 | 0.0021 | 1.018 | 0.0099 | -0.004 | 0.0057 |
| Mouth/teeth dental problems: Painful gums | 0 | -0.0838 | 0.1443 | -0.5806 | 0.5615 | 0.0081 | 0.0017 | 0.9954 | 0.0084 | -0.0006 | 0.0057 |
| Mouth/teeth dental problems: Toothache | 0 | 0.1817 | 0.1528 | 11.891 | 0.2344 | 0.0087 | 0.0017 | 10.035 | 0.0085 | -0.0044 | 0.0062 |
| Multiple sclerosis | 21833088 | -0.047 | 0.1987 | -0.2363 | 0.8132 | 0.0543 | 0.029 | 10.488 | 0.0102 | 0.0322 | 0.0065 |
| Nap during day | 0 | -0.0005 | 0.0611 | -0.0075 | 0.994 | 0.0804 | 0.0037 | 10.264 | 0.0131 | -0.0041 | 0.0076 |
| Neck/shoulder pain for 3+ months | 0 | -0.1853 | 0.1896 | -0.9774 | 0.3284 | 0.0188 | 0.0063 | 10.119 | 0.0078 | 0.0035 | 0.0056 |
| Neo-conscientiousness | 21173776 | 0.0463 | 0.2039 | 0.2273 | 0.8202 | 0.0799 | 0.0331 | 0.9973 | 0.0083 | 0.0016 | 0.0057 |
| Neo-openness to experience | 21173776 | -0.0202 | 0.1538 | -0.1313 | 0.8955 | 0.1141 | 0.029 | 0.9901 | 0.0079 | -0.0033 | 0.0056 |
| Nervous feelings | 0 | 0.0872 | 0.0606 | 14.393 | 0.1501 | 0.067 | 0.004 | 10.074 | 0.0136 | 0.0036 | 0.0064 |
| Neuroticism | 27089181 | 0.0071 | 0.0701 | 0.1006 | 0.9198 | 0.0891 | 0.0077 | 0.9895 | 0.0136 | 0.0041 | 0.0065 |

| | | | | | | | | | | | |
|---|---|---|---|---|---|---|---|---|---|---|---|
| Neuroticism | 24828478 | 0.139 | 0.1531 | 0.9085 | 0.3636 | 0.0119 | 0.0038 | 10.169 | 0.0083 | -0.0029 | 0.0055 |
| Neuroticism score | 0 | 0.002 | 0.0588 | 0.0343 | 0.9727 | 0.1212 | 0.0061 | 10.049 | 0.0149 | 0.0039 | 0.0069 |
| Noisy workplace | 0 | 0.0493 | 0.0989 | 0.4983 | 0.6182 | 0.0611 | 0.0059 | 10.296 | 0.0089 | 0.001 | 0.0061 |
| Non-accidental death in close genetic family | 0 | -0.2135 | 0.1787 | -11.947 | 0.2322 | 0.0182 | 0.0048 | 0.9942 | 0.0078 | 0.0035 | 0.0058 |
| Non-cancer illness code_ self-reported: allergy or anaphylactic r | 0 | -0.4644 | 0.8184 | -0.5675 | 0.5704 | 0.0006 | 0.0014 | 10.117 | 0.008 | 0.0034 | 0.006 |
| Non-cancer illness code_ self-reported: angina | 0 | -0.0557 | 0.0976 | -0.5703 | 0.5685 | 0.0212 | 0.002 | 10.293 | 0.0092 | 0.0041 | 0.006 |
| Non-cancer illness code_ self-reported: ankylosing spondylitis | 0 | 0.3927 | 0.381 | 10.308 | 0.3026 | 0.0015 | 0.0015 | 10.026 | 0.0079 | -0.0026 | 0.0061 |
| Non-cancer illness code_ self-reported: anxiety/panic attacks | 0 | -0.1491 | 0.157 | -0.9496 | 0.3423 | 0.006 | 0.0013 | 0.9991 | 0.0071 | 0.0079 | 0.0057 |
| Non-cancer illness code_ self-reported: arthritis (nos) | 0 | 0.1889 | 0.3604 | 0.5243 | 0.6001 | 0.0015 | 0.0014 | 10.159 | 0.0081 | -0.0055 | 0.0059 |
| Non-cancer illness code_ self-reported: asthma | 0 | -0.0463 | 0.0644 | -0.7193 | 0.4719 | 0.0577 | 0.0062 | 10.142 | 0.0189 | 0.0041 | 0.0065 |
| Non-cancer illness code_ self-reported: back problem | 0 | -0.0927 | 0.162 | -0.5725 | 0.567 | 0.0058 | 0.0015 | 0.9999 | 0.0074 | 0.0017 | 0.006 |
| Non-cancer illness code_ self-reported: bladder problem (not ca | 0 | 0.3738 | 0.2532 | 14.764 | 0.1398 | 0.0025 | 0.0014 | 0.996 | 0.0084 | -0.0192 | 0.0061 |
| Non-cancer illness code_ self-reported: bone disorder | 0 | 0.2196 | 0.4438 | 0.4948 | 0.6207 | 0.0008 | 0.0014 | 10.096 | 0.0074 | -0.0114 | 0.006 |
| Non-cancer illness code_ self-reported: cholelithiasis/gall stones | 0 | 0.0115 | 0.1444 | 0.0793 | 0.9368 | 0.0095 | 0.0021 | 10.127 | 0.0096 | -0.004 | 0.0065 |
| Non-cancer illness code_ self-reported: chronic obstructive airw | 0 | -0.1916 | 0.158 | -12.128 | 0.2252 | 0.0058 | 0.0016 | 0.9936 | 0.0076 | 0.0092 | 0.0058 |
| Non-cancer illness code_ self-reported: crohns disease | 0 | 0.0586 | 0.1581 | 0.3707 | 0.7109 | 0.0051 | 0.0016 | 10.058 | 0.0079 | -0.0042 | 0.0057 |
| Non-cancer illness code_ self-reported: deep venous thrombosis | 0 | 0.0671 | 0.1174 | 0.5718 | 0.5674 | 0.0122 | 0.0019 | 0.9952 | 0.0106 | -0.0065 | 0.0059 |
| Non-cancer illness code_ self-reported: depression | 0 | 0.0436 | 0.1039 | 0.4192 | 0.6751 | 0.0184 | 0.0018 | 10.004 | 0.0082 | 0.0028 | 0.0061 |
| Non-cancer illness code_ self-reported: diabetes | 0 | 0.0368 | 0.0776 | 0.4738 | 0.6357 | 0.0352 | 0.0026 | 10.347 | 0.0129 | -0.0052 | 0.0064 |
| Non-cancer illness code_ self-reported: diverticular disease/dive | 0 | 0.2061 | 0.1166 | 17.674 | 0.0772 | 0.0136 | 0.0018 | 0.9901 | 0.0082 | -0.0078 | 0.0059 |
| Non-cancer illness code_ self-reported: eczema/dermatitis | 0 | 0.1687 | 0.1375 | 1.227 | 0.2198 | 0.0088 | 0.0024 | 10.082 | 0.0083 | -0.0109 | 0.0055 |
| Non-cancer illness code_ self-reported: emphysema/chronic bro | 0 | 0.096 | 0.1282 | 0.749 | 0.4539 | 0.0101 | 0.0018 | 0.9847 | 0.0084 | -0.0044 | 0.006 |
| Non-cancer illness code_ self-reported: enlarged prostate | 0 | -0.3417 | 0.1647 | -20.743 | 0.0381 | 0.006 | 0.0015 | 10.063 | 0.0079 | 0.011 | 0.0065 |
| Non-cancer illness code_ self-reported: gastro-oesophageal reflu | 0 | -0.0936 | 0.1237 | -0.7563 | 0.4494 | 0.0116 | 0.0018 | 10.125 | 0.0087 | 0.0041 | 0.0064 |
| Non-cancer illness code_ self-reported: glaucoma | 0 | 0.0554 | 0.1189 | 0.4658 | 0.6414 | 0.0113 | 0.0018 | 10.119 | 0.0094 | 0.0025 | 0.0062 |
| Non-cancer illness code_ self-reported: gout | 0 | -0.0428 | 0.0869 | -0.4926 | 0.6223 | 0.0236 | 0.0071 | 0.985 | 0.0259 | 0.0026 | 0.0053 |
| Non-cancer illness code_ self-reported: hayfever/allergic rhinitis | 0 | 0.0818 | 0.0927 | 0.8817 | 0.3779 | 0.0247 | 0.0025 | 10.028 | 0.0103 | -0.013 | 0.0062 |
| Non-cancer illness code_ self-reported: heart attack/myocardial | 0 | -0.0294 | 0.0872 | -0.3369 | 0.7362 | 0.0197 | 0.0019 | 10.045 | 0.0085 | -0.0013 | 0.0056 |
| Non-cancer illness code_ self-reported: hiatus hernia | 0 | -0.1467 | 0.1596 | -0.9191 | 0.358 | 0.007 | 0.0015 | 10.383 | 0.0082 | 0.0064 | 0.0058 |
| Non-cancer illness code_ self-reported: high cholesterol | 0 | -0.0183 | 0.0736 | -0.2492 | 0.8032 | 0.0445 | 0.0044 | 10.152 | 0.0138 | -0.0051 | 0.0066 |
| Non-cancer illness code_ self-reported: hypertension | 0 | -0.0804 | 0.0543 | -14.793 | 0.1391 | 0.1182 | 0.0053 | 10.158 | 0.0175 | 0.0112 | 0.008 |
| Non-cancer illness code_ self-reported: hyperthyroidism/thyroto | 0 | 0.1495 | 0.129 | 1.159 | 0.2465 | 0.0083 | 0.002 | 0.9995 | 0.0087 | -0.0065 | 0.0056 |
| Non-cancer illness code_ self-reported: hypertrophic cardiomyo | 0 | -0.04 | 0.3319 | -0.1205 | 0.9041 | 0.0014 | 0.0014 | 1.003 | 0.0076 | -0.0027 | 0.0057 |
| Non-cancer illness code_ self-reported: hypopituitarism | 0 | -0.1624 | 0.3485 | -0.4662 | 0.6411 | 0.0013 | 0.0013 | 0.9953 | 0.0074 | 0.0004 | 0.006 |
| Non-cancer illness code_ self-reported: hypothyroidism/myxoed | 0 | 0.0998 | 0.0692 | 1.443 | 0.149 | 0.0513 | 0.0062 | 10.213 | 0.0182 | -0.0116 | 0.0066 |
| Non-cancer illness code_ self-reported: iron deficiency anaemia | 0 | 0.4653 | 0.2205 | 21.097 | 0.0349 | 0.0037 | 0.0014 | 0.996 | 0.0074 | -0.008 | 0.0053 |
| Non-cancer illness code_ self-reported: joint disorder | 0 | 0.2741 | 0.2659 | 10.306 | 0.3027 | 0.0025 | 0.0014 | 0.9879 | 0.0069 | -0.0044 | 0.006 |
| Non-cancer illness code_ self-reported: kidney stone/ureter ston | 0 | 0.237 | 0.1433 | 1.654 | 0.0981 | 0.0087 | 0.0017 | 10.048 | 0.0089 | -0.006 | 0.006 |
| Non-cancer illness code_ self-reported: malabsorption/coeliac d | 0 | 0.1009 | 0.1506 | 0.6703 | 0.5027 | 0.0093 | 0.0054 | 0.9997 | 0.0183 | 0.0027 | 0.0055 |
| Non-cancer illness code_ self-reported: mania/bipolar disorder/r | 0 | 0.0742 | 0.1691 | 0.439 | 0.6607 | 0.0055 | 0.0014 | 10.015 | 0.0073 | -0.0089 | 0.0059 |
| Non-cancer illness code_ self-reported: migraine | 0 | -0.0513 | 0.0981 | -0.5224 | 0.6014 | 0.0163 | 0.0019 | 1.012 | 0.0098 | 0.001 | 0.0061 |
| Non-cancer illness code_ self-reported: muscle or soft tissue inj | 0 | 0.0333 | 0.2606 | 0.1278 | 0.8983 | 0.002 | 0.0015 | 1.002 | 0.008 | 0.0011 | 0.0059 |
| Non-cancer illness code_ self-reported: nasal polyps | 0 | -0.0134 | 0.163 | -0.0821 | 0.9345 | 0.0049 | 0.0017 | 10.091 | 0.0082 | 0.0024 | 0.0054 |
| Non-cancer illness code_ self-reported: osteoarthritis | 0 | 0.0157 | 0.1025 | 0.1535 | 0.878 | 0.0197 | 0.0018 | 10.175 | 0.0085 | -0.0065 | 0.0068 |
| Non-cancer illness code_ self-reported: osteoporosis | 0 | -0.071 | 0.109 | -0.6512 | 0.5149 | 0.014 | 0.002 | 10.059 | 0.0092 | 0.0003 | 0.0057 |
| Non-cancer illness code_ self-reported: pernicious anaemia | 0 | 0.3386 | 0.2284 | 14.825 | 0.1382 | 0.0029 | 0.0014 | 0.9938 | 0.007 | -0.0067 | 0.0054 |
| Non-cancer illness code_ self-reported: pneumothorax | 0 | 0.2133 | 0.1902 | 11.216 | 0.262 | 0.004 | 0.0015 | 0.9816 | 0.0076 | -0.0048 | 0.0058 |

| | | | | | | | | | | | |
|---|---|---|---|---|---|---|---|---|---|---|---|
| Non-cancer illness code_ self-reported: polio / poliomyelitis | 0 | 0.1877 | 0.2826 | 0.6641 | 0.5066 | 0.0017 | 0.0013 | 0.9986 | 0.0073 | -0.0079 | 0.0057 |
| Non-cancer illness code_ self-reported: psoriasis | 0 | -0.0743 | 0.1291 | -0.5757 | 0.5648 | 0.0077 | 0.0017 | 10.087 | 0.0101 | 0.0066 | 0.0053 |
| Non-cancer illness code_ self-reported: pulmonary embolism +/ | 0 | 0.1002 | 0.159 | 0.6302 | 0.5285 | 0.0058 | 0.0015 | 10.024 | 0.0084 | -0.0009 | 0.0057 |
| Non-cancer illness code_ self-reported: retinal detachment | 0 | -0.2856 | 0.2198 | -12.989 | 0.194 | 0.003 | 0.0015 | 10.067 | 0.0082 | 0.0083 | 0.0058 |
| Non-cancer illness code_ self-reported: rheumatoid arthritis | 0 | -0.0289 | 0.1819 | -0.1588 | 0.8738 | 0.0045 | 0.0015 | 10.048 | 0.0074 | 0.005 | 0.0061 |
| Non-cancer illness code_ self-reported: sleep apnoea | 0 | -0.1096 | 0.1917 | -0.572 | 0.5673 | 0.0038 | 0.0015 | 10.026 | 0.0086 | 0.0042 | 0.0056 |
| Non-cancer illness code_ self-reported: type 2 diabetes | 0 | -0.0835 | 0.1508 | -0.5538 | 0.5797 | 0.0061 | 0.0014 | 10.113 | 0.0083 | 0.007 | 0.005 |
| Non-cancer illness code_ self-reported: ulcerative colitis | 0 | -0.1096 | 0.1438 | -0.7626 | 0.4457 | 0.0063 | 0.0018 | 10.062 | 0.0087 | 0.0038 | 0.0054 |
| Non-cancer illness code_ self-reported: uterine fibroids | 0 | 0.1931 | 0.164 | 11.776 | 0.239 | 0.0067 | 0.0017 | 10.113 | 0.0087 | -0.0129 | 0.0058 |
| Non-cancer illness code_ self-reported: vaginal prolapse/uterine | 0 | -0.0208 | 0.1804 | -0.1154 | 0.9082 | 0.0054 | 0.0015 | 0.9995 | 0.0074 | 0.002 | 0.0058 |
| Non-cancer illness code_ self-reported: varicose veins | 0 | -0.1456 | 0.2122 | -0.686 | 0.4927 | 0.0042 | 0.0015 | 10.127 | 0.0077 | 0.0029 | 0.0066 |
| Non-cancer illness code_ self-reported: vitiligo | 0 | 0.1459 | 0.3323 | 0.439 | 0.6607 | 0.0013 | 0.0014 | 0.9956 | 0.0079 | -0.0007 | 0.0057 |
| No-wear time bias adjusted acceleration standard deviation | 0 | -0.1381 | 0.0942 | -14.656 | 0.1428 | 0.0885 | 0.0087 | 10.031 | 0.0093 | 0.0042 | 0.0054 |
| Number of children ever born | 27798627 | 0.1914 | 0.092 | 20.792 | 0.0376 | 0.0247 | 0.0018 | 0.9786 | 0.0082 | -0.0083 | 0.0063 |
| Number of children fathered | 0 | 0.0006 | 0.112 | 0.0051 | 0.9959 | 0.031 | 0.0042 | 10.152 | 0.0083 | 0.0028 | 0.0063 |
| Number of cigarettes currently smoked daily (current cigarette s | 0 | 0.0697 | 0.1636 | 0.426 | 0.6701 | 0.1092 | 0.0266 | 0.9952 | 0.0079 | -0.0036 | 0.0064 |
| Number of cigarettes previously smoked daily | 0 | -0.0102 | 0.0898 | -0.1131 | 0.91 | 0.0968 | 0.0136 | 10.029 | 0.0101 | -0.0049 | 0.0064 |
| Number of days/week of moderate physical activity 10+ minute | 0 | -0.0644 | 0.0756 | -0.851 | 0.3947 | 0.0423 | 0.0027 | 10.118 | 0.0104 | 0.0074 | 0.0064 |
| Number of days/week of vigorous physical activity 10+ minutes | 0 | 0.0403 | 0.0811 | 0.4967 | 0.6194 | 0.0376 | 0.0026 | 10.173 | 0.0097 | -0.003 | 0.0064 |
| Number of days/week walked 10+ minutes | 0 | 0.0854 | 0.0682 | 12.528 | 0.2103 | 0.0428 | 0.0025 | 10.093 | 0.0088 | -0.0032 | 0.006 |
| Number of depression episodes | 0 | 0.3254 | 0.1719 | 18.935 | 0.0583 | 0.0413 | 0.0129 | 0.9955 | 0.0084 | -0.0013 | 0.0055 |
| Number of full brothers | 0 | 0.0869 | 0.1019 | 0.8525 | 0.3939 | 0.0196 | 0.0019 | 10.211 | 0.0081 | 6,45E-01 | 0.0064 |
| Number of full sisters | 0 | 0.1241 | 0.0989 | 1.254 | 0.2099 | 0.0186 | 0.002 | 10.379 | 0.009 | 0.0045 | 0.0058 |
| Number of incorrect matches in round | 0 | -0.0206 | 0.0646 | -0.319 | 0.7498 | 0.0553 | 0.0028 | 0.9977 | 0.0106 | 0.0015 | 0.0062 |
| Number of live births | 0 | 0.1178 | 0.0857 | 13.754 | 0.169 | 0.0635 | 0.0044 | 0.9975 | 0.0092 | 0.0019 | 0.0067 |
| Number of older siblings | 0 | -0.2967 | 0.4456 | -0.6659 | 0.5055 | 0.0037 | 0.0052 | 10.109 | 0.0072 | 0.0073 | 0.0056 |
| Number of operations_ self-reported | 0 | -0.0091 | 0.0795 | -0.1139 | 0.9093 | 0.0384 | 0.0023 | 10.296 | 0.0091 | 0.0082 | 0.0066 |
| Number of pregnancy terminations | 0 | 0.0498 | 0.1371 | 0.3635 | 0.7162 | 0.0572 | 0.0107 | 10.089 | 0.0088 | 0.0013 | 0.0058 |
| Number of self-reported cancers | 0 | -0.0889 | 0.1485 | -0.5989 | 0.5493 | 0.0072 | 0.0017 | 10.075 | 0.0087 | -0.0048 | 0.0058 |
| Number of self-reported non-cancer illnesses | 0 | 0.0252 | 0.0591 | 0.427 | 0.6694 | 0.064 | 0.0027 | 10.064 | 0.0111 | -0.0048 | 0.0068 |
| Number of treatments/medications taken | 0 | -0.0474 | 0.0631 | -0.7525 | 0.4518 | 0.0619 | 0.0027 | 10.203 | 0.0106 | 0.0031 | 0.007 |
| Number of trend entries | 0 | 0.2118 | 0.2041 | 10.381 | 0.2992 | 0.0329 | 0.0104 | 10.054 | 0.0078 | -0.0044 | 0.0066 |
| Number of unsuccessful stop-smoking attempts | 0 | -0.0425 | 0.1198 | -0.3548 | 0.7227 | 0.0488 | 0.008 | 10.077 | 0.0081 | 0.007 | 0.006 |
| Obesity class 1 | 23563607 | 0.0544 | 0.0632 | 0.8611 | 0.3892 | 0.2212 | 0.0118 | 10.056 | 0.0123 | 0.0006 | 0.007 |
| Obesity class 2 | 23563607 | 0.1357 | 0.0844 | 16.076 | 0.1079 | 0.185 | 0.0139 | 0.9984 | 0.0116 | -0.005 | 0.0076 |
| Obesity class 3 | 23563607 | 0.215 | 0.1145 | 18.772 | 0.0605 | 0.1206 | 0.0148 | 0.9804 | 0.0096 | -0.0043 | 0.007 |
| Omega-3 fatty acids | 27005778 | 0.0124 | 0.1914 | 0.065 | 0.9482 | 0.1407 | 0.0425 | 0.9914 | 0.0093 | -0.0059 | 0.0059 |
| Omega-9 and saturated fatty acids | 27005778 | 0.157 | 0.2404 | 0.653 | 0.5137 | 0.094 | 0.0447 | 0.9937 | 0.0086 | -0.0078 | 0.0062 |
| Other eye problems | 0 | 0.0713 | 0.1252 | 0.5696 | 0.5689 | 0.0094 | 0.0017 | 10.221 | 0.0082 | 0.0073 | 0.0059 |
| Other serious medical condition/disability diagnosed by doctor | 0 | 0.0699 | 0.0793 | 0.8811 | 0.3782 | 0.0252 | 0.0019 | 10.108 | 0.0088 | -0.0011 | 0.0055 |
| Overall health rating | 0 | -0.0346 | 0.0544 | -0.6358 | 0.5249 | 0.0991 | 0.0034 | 10.619 | 0.0129 | 0.0047 | 0.007 |
| Overweight | 23563607 | 0.0578 | 0.0658 | 0.8789 | 0.3795 | 0.1145 | 0.0069 | 10.055 | 0.0109 | 0.0039 | 0.0065 |
| Pack years adult smoking as proportion of life span exposed to s | 0 | -0.0608 | 0.0777 | -0.7826 | 0.4339 | 0.1263 | 0.0119 | 10.146 | 0.0108 | -0.0033 | 0.0064 |
| Pack years of smoking PREVIEW ONLY | 0 | -0.0849 | 0.0812 | -10.452 | 0.2959 | 0.11 | 0.0104 | 10.129 | 0.0103 | -0.0017 | 0.0065 |
| Pain type(s) experienced in last month: Back pain | 0 | -0.0618 | 0.0755 | -0.8185 | 0.4131 | 0.0409 | 0.0023 | 10.141 | 0.0096 | 0.0046 | 0.0063 |
| Pain type(s) experienced in last month: Facial pain | 0 | -0.0371 | 0.1615 | -0.2298 | 0.8182 | 0.0064 | 0.0016 | 0.9982 | 0.0076 | 0.0081 | 0.0061 |
| Pain type(s) experienced in last month: Headache | 0 | -0.0376 | 0.0748 | -0.5024 | 0.6154 | 0.0451 | 0.003 | 0.9978 | 0.0109 | 0.0002 | 0.0068 |

| | | | | | | | | | | | |
|---|---|---|---|---|---|---|---|---|---|---|---|
| Pain type(s) experienced in last month: Hip pain | 0 | -0.0012 | 0.1007 | -0.0121 | 0.9903 | 0.0234 | 0.0021 | 10.157 | 0.0092 | 0.0026 | 0.0065 |
| Pain type(s) experienced in last month: Knee pain | 0 | -0.0177 | 0.0752 | -0.2358 | 0.8136 | 0.0413 | 0.0026 | 10.072 | 0.0107 | 0.002 | 0.0063 |
| Pain type(s) experienced in last month: Neck or shoulder pain | 0 | 0.0638 | 0.0815 | 0.7828 | 0.4337 | 0.0331 | 0.0023 | 10.225 | 0.01 | -0.0037 | 0.0064 |
| Pain type(s) experienced in last month: None of the above | 0 | -0.0112 | 0.0683 | -0.1648 | 0.8691 | 0.0557 | 0.0029 | 10.313 | 0.0106 | -0.0014 | 0.0069 |
| Pain type(s) experienced in last month: Pain all over the body | 0 | -0.1476 | 0.1526 | -0.9672 | 0.3335 | 0.0096 | 0.0017 | 10.109 | 0.0085 | 0.0058 | 0.0064 |
| Pain type(s) experienced in last month: Stomach or abdominal p | 0 | -0.0626 | 0.0853 | -0.7338 | 0.4631 | 0.0213 | 0.0018 | 10.079 | 0.0083 | 0.0036 | 0.0061 |
| Parents age at death | 27015805 | -0.0491 | 0.1721 | -0.2856 | 0.7752 | 0.0284 | 0.0077 | 10.186 | 0.0081 | -0.0021 | 0.0065 |
| Parkinsons disease | 19915575 | 0.1862 | 0.1242 | 14.989 | 0.1339 | 0.4497 | 0.1254 | 11.094 | 0.0101 | -0.0253 | 0.0053 |
| Past tobacco smoking | 0 | 0.0626 | 0.0613 | 10.212 | 0.3072 | 0.0912 | 0.0042 | 10.185 | 0.0133 | -0.0168 | 0.0066 |
| Peak expiratory flow (PEF) | 0 | 0.0609 | 0.0575 | 10.596 | 0.2893 | 0.0924 | 0.0054 | 10.345 | 0.0152 | -0.0019 | 0.007 |
| PGC cross-disorder analysis | 23453885 | -0.0294 | 0.0875 | -0.3363 | 0.7367 | 0.1623 | 0.0129 | 1.025 | 0.0125 | 0.0255 | 0.0079 |
| Phenylalanine | 27005778 | -0.1408 | 0.2015 | -0.6984 | 0.485 | 0.0649 | 0.025 | 0.9995 | 0.008 | 0.0025 | 0.0058 |
| Phospholipids in chylomicrons and largest VLDL particles | 27005778 | -0.2041 | 0.153 | -13.337 | 0.1823 | 0.1062 | 0.0278 | 0.9779 | 0.0078 | 0.0026 | 0.0054 |
| Phospholipids in IDL | 27005778 | 0.0287 | 0.1756 | 0.1636 | 0.8701 | 0.1018 | 0.0321 | 0.9971 | 0.0118 | -0.0071 | 0.0061 |
| Phospholipids in large HDL | 27005778 | -0.1333 | 0.1701 | -0.7837 | 0.4332 | 0.1223 | 0.033 | 0.9972 | 0.0153 | 0.0003 | 0.0063 |
| Phospholipids in large LDL | 27005778 | 0.1245 | 0.1725 | 0.7218 | 0.4704 | 0.1072 | 0.0298 | 0.9912 | 0.012 | -0.0086 | 0.0062 |
| Phospholipids in large VLDL | 27005778 | -0.1561 | 0.1327 | -11.768 | 0.2393 | 0.1283 | 0.0344 | 0.9795 | 0.0081 | 0.0053 | 0.0049 |
| Phospholipids in medium HDL | 27005778 | -0.1919 | 0.1882 | -10.198 | 0.3078 | 0.0737 | 0.026 | 0.9909 | 0.0092 | 0.0004 | 0.0056 |
| Phospholipids in medium LDL | 27005778 | 0.1592 | 0.1741 | 0.9144 | 0.3605 | 0.1136 | 0.0309 | 0.9864 | 0.0117 | -0.0092 | 0.0063 |
| Phospholipids in medium VLDL | 27005778 | -0.0267 | 0.1499 | -0.1781 | 0.8586 | 0.1203 | 0.036 | 0.9879 | 0.0078 | -0.0005 | 0.0054 |
| Phospholipids in small VLDL | 27005778 | -0.0015 | 0.1554 | -0.0094 | 0.9925 | 0.1261 | 0.0371 | 0.9903 | 0.0093 | -0.0036 | 0.0059 |
| Phospholipids in very large HDL | 27005778 | -0.0634 | 0.1914 | -0.3311 | 0.7405 | 0.0919 | 0.0326 | 10.082 | 0.0188 | -0.0023 | 0.0061 |
| Phospholipids in very large VLDL | 27005778 | -0.2228 | 0.1535 | -14.514 | 0.1467 | 0.1151 | 0.0308 | 0.9777 | 0.0078 | 0.0039 | 0.0055 |
| Phospholipids in very small VLDL | 27005778 | 0.0466 | 0.1786 | 0.2606 | 0.7944 | 0.1061 | 0.0354 | 10.009 | 0.0128 | -0.0075 | 0.0062 |
| Platelet count | 22139419 | 0.0974 | 0.0781 | 12.477 | 0.2121 | 0.1146 | 0.013 | 0.9861 | 0.0107 | -0.0035 | 0.0051 |
| Potassium in urine | 0 | -0.0245 | 0.0685 | -0.3571 | 0.721 | 0.0431 | 0.0025 | 10.114 | 0.0101 | -0.0037 | 0.0061 |
| Primary biliary cirrhosis | 26394269 | -0.0288 | 0.1156 | -0.2494 | 0.803 | 0.4018 | 0.0675 | 0.9874 | 0.0113 | 0.0018 | 0.0068 |
| Primary sclerosing cholangitis | 27992413 | 0.093 | 0.1254 | 0.741 | 0.4587 | 0.3221 | 0.0825 | 0.992 | 0.0145 | 0.0313 | 0.0065 |
| Prospective memory result | 0 | 0.1015 | 0.1 | 1.015 | 0.3101 | 0.0585 | 0.0059 | 10.113 | 0.0095 | -0.0052 | 0.0065 |
| Pulse rate | 0 | 0.0326 | 0.0686 | 0.4752 | 0.6347 | 0.1379 | 0.0149 | 1.015 | 0.0145 | -0.0036 | 0.0062 |
| Pulse rate_ automated reading | 0 | 0.0412 | 0.0514 | 0.8025 | 0.4223 | 0.1573 | 0.0135 | 1.014 | 0.0313 | -0.0076 | 0.007 |
| Pulse wave Arterial Stiffness index | 0 | -0.3185 | 0.121 | -26.331 | 0.0085 | 0.0396 | 0.0054 | 10.024 | 0.0086 | 0.0096 | 0.0062 |
| Pulse wave peak to peak time | 0 | 0.3463 | 0.1221 | 2.837 | 0.0046 | 0.0398 | 0.0052 | 10.041 | 0.0084 | -0.0088 | 0.0062 |
| Pulse wave reflection index | 0 | -0.1552 | 0.1235 | -12.563 | 0.209 | 0.0539 | 0.0058 | 10.073 | 0.0084 | 0.0022 | 0.0068 |
| Qualifications: A levels/AS levels or equivalent | 0 | 0.0537 | 0.0597 | 0.8988 | 0.3687 | 0.0951 | 0.0041 | 10.602 | 0.0146 | -0.0078 | 0.0076 |
| Qualifications: College or University degree | 0 | 0.0966 | 0.0537 | 17.997 | 0.0719 | 0.1687 | 0.006 | 11.145 | 0.0177 | -0.0141 | 0.0086 |
| Qualifications: CSEs or equivalent | 0 | -0.0888 | 0.0891 | -0.9967 | 0.3189 | 0.0202 | 0.002 | 10.614 | 0.0088 | 0.0024 | 0.0057 |
| Qualifications: None of the above | 0 | -0.0682 | 0.0609 | -11.187 | 0.2633 | 0.0984 | 0.0043 | 10.638 | 0.0138 | 0.0089 | 0.0077 |
| Qualifications: NVQ or HND or HNC or equivalent | 0 | -0.0682 | 0.1112 | -0.6137 | 0.5394 | 0.0148 | 0.0018 | 10.111 | 0.0086 | -0.0012 | 0.0061 |
| Qualifications: O levels/GCSEs or equivalent | 0 | -0.0101 | 0.0675 | -0.1498 | 0.8809 | 0.0497 | 0.0026 | 10.091 | 0.0106 | 0.0006 | 0.0065 |
| Qualifications: Other professional qualifications eg: nursing_ te | 0 | -0.0137 | 0.0678 | -0.2019 | 0.84 | 0.0479 | 0.0025 | 10.253 | 0.0099 | 0.0035 | 0.0066 |
| Ratio of bisallylic groups to double bonds | 27005778 | -0.1037 | 0.1284 | -0.8074 | 0.4194 | 0.2706 | 0.0761 | 0.967 | 0.0079 | 0.0008 | 0.0056 |
| Ratio of bisallylic groups to total fatty acids | 27005778 | -0.0234 | 0.1323 | -0.1765 | 0.8599 | 0.2714 | 0.0728 | 0.9708 | 0.0075 | -0.0025 | 0.0056 |
| Reason for glasses/contact lenses: For astigmatism | 0 | 0.1245 | 0.1309 | 0.9512 | 0.3415 | 0.0092 | 0.0016 | 1.027 | 0.0079 | 0.0057 | 0.0059 |
| Reason for glasses/contact lenses: For just reading/near work as | 0 | 0.0986 | 0.2183 | 0.4516 | 0.6515 | 0.0034 | 0.0018 | 11.367 | 0.0093 | -0.0033 | 0.0054 |
| Reason for glasses/contact lenses: For long-sightedness_ i.e. for | 0 | -0.0844 | 0.1386 | -0.6092 | 0.5424 | 0.009 | 0.0016 | 10.697 | 0.0092 | 0.0068 | 0.006 |
| Reason for glasses/contact lenses: For short-sightedness_ i.e. on | 0 | 0.0601 | 0.0917 | 0.6549 | 0.5125 | 0.0242 | 0.0021 | 11.221 | 0.0103 | -0.0045 | 0.0064 |

| | | | | | | | | | | | |
|---|---|---|---|---|---|---|---|---|---|---|---|
| Reason for reducing amount of alcohol drunk: Health precautior | 0 | -0.0681 | 0.0963 | -0.7073 | 0.4794 | 0.0474 | 0.0049 | 0.996 | 0.0086 | 0.0021 | 0.0065 |
| Relative age of first facial hair | 0 | 0.0195 | 0.0675 | 0.2883 | 0.7731 | 0.1192 | 0.0098 | 10.493 | 0.0219 | 0.0051 | 0.0067 |
| Relative age voice broke | 0 | 0.0352 | 0.0777 | 0.4535 | 0.6502 | 0.0741 | 0.0059 | 10.036 | 0.0107 | 8,92E-02 | 0.0058 |
| Reproduciblity of spirometry measurement using ERS/ATS crit | 0 | -0.1322 | 0.1258 | -10.508 | 0.2933 | 0.0147 | 0.0023 | 10.055 | 0.0086 | 0.0023 | 0.0058 |
| Rheumatoid Arthritis | 24390342 | -0.0497 | 0.0806 | -0.6161 | 0.5378 | 0.1683 | 0.0337 | 10.032 | 0.0185 | 0.0088 | 0.0061 |
| Risk taking | 0 | 0.0313 | 0.0633 | 0.4941 | 0.6213 | 0.0569 | 0.0031 | 0.9971 | 0.0112 | 0.0049 | 0.0062 |
| Schizophrenia | 25056061 | 0.0566 | 0.0651 | 0.8694 | 0.3846 | 0.4516 | 0.0189 | 10.553 | 0.0149 | 0.0116 | 0.0084 |
| Seen a psychiatrist for nerves_ anxiety_ tension or depression | 0 | 0.1189 | 0.0822 | 14.466 | 0.148 | 0.0305 | 0.0023 | 10.098 | 0.0097 | -0.002 | 0.0057 |
| Seen doctor (GP) for nerves_ anxiety_ tension or depression | 0 | 0.0257 | 0.0636 | 0.4036 | 0.6865 | 0.0618 | 0.0034 | 10.141 | 0.0115 | 0.0073 | 0.0062 |
| Sensitivity / hurt feelings | 0 | -0.0745 | 0.072 | -10.352 | 0.3006 | 0.0635 | 0.0034 | 1.018 | 0.0113 | 0.0077 | 0.0074 |
| Serum creatinine | 26831199 | 0.0683 | 0.0826 | 0.8272 | 0.4081 | 0.1105 | 0.0102 | 0.9625 | 0.0156 | -0.0012 | 0.0067 |
| Serum creatinine (non-diabetes) | 26831199 | 0.0774 | 0.0871 | 0.8885 | 0.3743 | 0.1186 | 0.0117 | 0.9645 | 0.0158 | -0.0014 | 0.0066 |
| Serum cystatin c | 26831199 | -0.061 | 0.0998 | -0.6118 | 0.5407 | 0.1742 | 0.0675 | 0.9558 | 0.0155 | 0.0025 | 0.0057 |
| Serum total cholesterol | 27005778 | 0.0132 | 0.1744 | 0.0755 | 0.9399 | 0.1101 | 0.0349 | 0.9965 | 0.0114 | -0.0059 | 0.0062 |
| Serum total triglycerides | 27005778 | -0.0389 | 0.1442 | -0.2698 | 0.7873 | 0.141 | 0.0376 | 0.9815 | 0.0078 | -0.0013 | 0.0057 |
| Serumurate overweight | 25811787 | -0.067 | 0.1053 | -0.6363 | 0.5246 | 0.5411 | 0.3209 | 0.9606 | 0.0327 | 0.0017 | 0.0057 |
| Shortness of breath walking on level ground | 0 | -0.0686 | 0.0934 | -0.7347 | 0.4625 | 0.0506 | 0.0054 | 10.043 | 0.0085 | 0.002 | 0.0059 |
| Sitting height | 0 | 0.0724 | 0.0485 | 14.936 | 0.1353 | 0.3284 | 0.0195 | 10.355 | 0.0471 | 0.0109 | 0.0094 |
| Sitting height ratio | 25865494 | -0.0998 | 0.1152 | -0.8659 | 0.3866 | 0.2099 | 0.0274 | 0.9812 | 0.0084 | 0.0048 | 0.0061 |
| Sleep duration | 27494321 | 0.0633 | 0.1039 | 0.6093 | 0.5423 | 0.0565 | 0.0053 | 10.155 | 0.0092 | -0.01 | 0.006 |
| Sleep duration | 0 | 0.0112 | 0.0677 | 0.1657 | 0.8684 | 0.0724 | 0.0036 | 10.161 | 0.0114 | -0.002 | 0.0068 |
| Sleeplessness / insomnia | 0 | -0.0137 | 0.071 | -0.1925 | 0.8473 | 0.0624 | 0.0031 | 10.181 | 0.0113 | 0.0073 | 0.0072 |
| Smoking status: Current | 0 | -0.037 | 0.0582 | -0.6358 | 0.5249 | 0.0521 | 0.0031 | 10.144 | 0.0111 | 0.0014 | 0.0057 |
| Smoking status: Previous | 0 | -0.0674 | 0.0732 | -0.9205 | 0.3573 | 0.0524 | 0.0031 | 10.153 | 0.0118 | 0.016 | 0.0065 |
| Smoking/smokers in household | 0 | -0.1564 | 0.1326 | -11.794 | 0.2382 | 0.0092 | 0.0022 | 10.176 | 0.0095 | 0.0047 | 0.0059 |
| Snoring | 0 | 0.0463 | 0.0689 | 0.6723 | 0.5014 | 0.0615 | 0.003 | 10.117 | 0.0104 | -0.0011 | 0.0072 |
| Sodium in urine | 0 | -0.0731 | 0.0597 | -12.257 | 0.2203 | 0.0725 | 0.0034 | 10.183 | 0.0113 | 0.0093 | 0.0066 |
| Squamous cell lung cancer | 27488534 | -0.0238 | 0.1511 | -0.1575 | 0.8748 | 0.0402 | 0.0122 | 10.072 | 0.0077 | 0.0058 | 0.0054 |
| Standing height | 0 | 0.1 | 0.0439 | 22.803 | 0.0226 | 0.4561 | 0.0243 | 1.094 | 0.0597 | 0.0065 | 0.0105 |
| Started insulin within one year diagnosis of diabetes | 0 | 0.246 | 0.1391 | 17.678 | 0.0771 | 0.1541 | 0.0459 | 10.035 | 0.0088 | -0.0147 | 0.0054 |
| Subjective well being | 27089181 | -0.0516 | 0.098 | -0.5263 | 0.5987 | 0.0252 | 0.0022 | 1.001 | 0.0081 | 0.0042 | 0.0059 |
| Suffer from nerves | 0 | 0.1861 | 0.0697 | 26.719 | 0.0075 | 0.047 | 0.0032 | 0.9898 | 0.0114 | -0.0003 | 0.0059 |
| Systemic lupus erythematosus | 26502338 | -0.019 | 0.1088 | -0.1744 | 0.8615 | 0.4156 | 0.0721 | 10.842 | 0.0126 | 0.0137 | 0.006 |
| Systolic blood pressure_ automated reading | 0 | -0.0767 | 0.057 | -13.447 | 0.1787 | 0.1304 | 0.0059 | 10.367 | 0.0173 | 0.0104 | 0.0075 |
| Taking other prescription medications | 0 | 0.0803 | 0.0697 | 11.522 | 0.2492 | 0.0448 | 0.0025 | 10.066 | 0.0103 | -0.0031 | 0.0064 |
| Target heart rate achieved | 0 | -0.3358 | 0.1725 | -19.468 | 0.0516 | 0.0357 | 0.0112 | 1.002 | 0.0077 | 0.0142 | 0.006 |
| Tense / highly strung | 0 | 0.1312 | 0.0651 | 20.145 | 0.044 | 0.0597 | 0.0032 | 0.9834 | 0.011 | -0.0078 | 0.006 |
| Time from waking to first cigarette | 0 | 0.1058 | 0.1544 | 0.6853 | 0.4931 | 0.0983 | 0.0267 | 10.088 | 0.0088 | 0.0004 | 0.0057 |
| Time spent driving | 0 | 0.0468 | 0.0843 | 0.5559 | 0.5783 | 0.0444 | 0.003 | 0.9949 | 0.0086 | 0.0017 | 0.0063 |
| Time spent using computer | 0 | 0.1495 | 0.0721 | 20.727 | 0.0382 | 0.0969 | 0.0043 | 10.403 | 0.0121 | -0.0061 | 0.008 |
| Time spent watching television (TV) | 0 | -0.1168 | 0.0609 | -19.178 | 0.0551 | 0.0998 | 0.0042 | 10.512 | 0.0131 | 0.0052 | 0.0076 |
| Tinnitus: Yes_ but not now_ but have in the past | 0 | -0.1978 | 0.1545 | -1.28 | 0.2005 | 0.0185 | 0.0047 | 0.9899 | 0.0078 | 0.0031 | 0.0054 |
| Tinnitus: Yes_ now most or all of the time | 0 | -0.0686 | 0.1216 | -0.5643 | 0.5725 | 0.0311 | 0.005 | 10.061 | 0.0077 | 0.0045 | 0.0053 |
| Tinnitus: Yes_ now some of the time | 0 | -0.1432 | 0.2157 | -0.664 | 0.5067 | 0.009 | 0.0043 | 0.9987 | 0.0073 | 0.005 | 0.0056 |
| Tobacco smoking: Ex-smoker | 0 | -0.1307 | 0.1062 | -12.305 | 0.2185 | 0.0758 | 0.0089 | 10.096 | 0.0099 | 0.0126 | 0.0062 |
| Total Cholesterol | 20686565 | -0.07 | 0.0723 | -0.9684 | 0.3328 | 0.1453 | 0.0195 | 0.9815 | 0.0192 | 0.0027 | 0.0068 |
| Total cholesterol in HDL | 27005778 | -0.101 | 0.1737 | -0.5811 | 0.5611 | 0.0905 | 0.0271 | 10.052 | 0.013 | -0.0009 | 0.006 |

| | | | | | | | | | | | |
|---|---|---|---|---|---|---|---|---|---|---|---|
| Total cholesterol in IDL | 27005778 | 0.1676 | 0.1766 | 0.9495 | 0.3424 | 0.1162 | 0.0357 | 0.9941 | 0.013 | -0.0093 | 0.0062 |
| Total cholesterol in large HDL | 27005778 | -0.0457 | 0.1772 | -0.2581 | 0.7963 | 0.1006 | 0.028 | 10.061 | 0.0155 | -0.0009 | 0.0062 |
| Total cholesterol in large LDL | 27005778 | 0.135 | 0.1678 | 0.8041 | 0.4213 | 0.1143 | 0.0308 | 0.9864 | 0.0127 | -0.009 | 0.0061 |
| Total cholesterol in large VLDL | 27005778 | -0.1332 | 0.1456 | -0.9153 | 0.36 | 0.129 | 0.0307 | 0.9799 | 0.0075 | 0.0017 | 0.0054 |
| Total cholesterol in LDL | 27005778 | 0.1333 | 0.1692 | 0.7878 | 0.4308 | 0.1194 | 0.0335 | 0.9856 | 0.0129 | -0.0093 | 0.0062 |
| Total cholesterol in medium HDL | 27005778 | -0.2233 | 0.2182 | -10.234 | 0.3061 | 0.0564 | 0.0279 | 0.9968 | 0.0106 | 0.0027 | 0.0058 |
| Total cholesterol in medium LDL | 27005778 | 0.1454 | 0.1731 | 0.8401 | 0.4009 | 0.1149 | 0.0313 | 0.9873 | 0.0124 | -0.0096 | 0.0063 |
| Total cholesterol in medium VLDL | 27005778 | -0.0192 | 0.1498 | -0.1285 | 0.8978 | 0.1293 | 0.0378 | 0.9853 | 0.0084 | -0.0013 | 0.0057 |
| Total cholesterol in small LDL | 27005778 | 0.0942 | 0.1783 | 0.5286 | 0.5971 | 0.114 | 0.0358 | 0.9923 | 0.0118 | -0.0091 | 0.0064 |
| Total cholesterol in small VLDL | 27005778 | 0.006 | 0.1799 | 0.0335 | 0.9733 | 0.1012 | 0.0331 | 0.9965 | 0.0095 | -0.0044 | 0.0061 |
| Total cholesterol in very large HDL | 27005778 | 0.1732 | 0.25 | 0.6929 | 0.4884 | 0.0408 | 0.0241 | 10.098 | 0.0131 | -0.0056 | 0.0059 |
| Total lipids in chylomicrons and largest VLDL particles | 27005778 | -0.1296 | 0.1481 | -0.8756 | 0.3813 | 0.1267 | 0.0281 | 0.9831 | 0.0081 | -8,00E-01 | 0.0055 |
| Total lipids in IDL | 27005778 | 0.1234 | 0.1795 | 0.6875 | 0.4918 | 0.1123 | 0.035 | 0.9973 | 0.0126 | -0.0089 | 0.0062 |
| Total lipids in large HDL | 27005778 | -0.1122 | 0.1705 | -0.6581 | 0.5105 | 0.1245 | 0.0326 | 10.006 | 0.0161 | 0.0002 | 0.0063 |
| Total lipids in large LDL | 27005778 | 0.1498 | 0.174 | 0.8609 | 0.3893 | 0.1274 | 0.0367 | 0.988 | 0.0127 | -0.0096 | 0.0063 |
| Total lipids in large VLDL | 27005778 | -0.0631 | 0.1276 | -0.4945 | 0.621 | 0.1485 | 0.0324 | 0.9757 | 0.0078 | 0.0036 | 0.005 |
| Total lipids in medium HDL | 27005778 | -0.2925 | 0.2186 | -1.338 | 0.1809 | 0.0662 | 0.0288 | 0.9936 | 0.0092 | 0.0036 | 0.0058 |
| Total lipids in medium LDL | 27005778 | 0.1467 | 0.1754 | 0.8363 | 0.403 | 0.1299 | 0.0371 | 0.9859 | 0.0123 | -0.0097 | 0.0064 |
| Total lipids in medium VLDL | 27005778 | 0.0308 | 0.1407 | 0.2187 | 0.8269 | 0.1468 | 0.0365 | 0.9792 | 0.0078 | -0.0005 | 0.0053 |
| Total lipids in small HDL | 27005778 | -0.0213 | 0.2058 | -0.1034 | 0.9176 | 0.0668 | 0.0276 | 0.9994 | 0.0085 | -0.0019 | 0.0056 |
| Total lipids in small LDL | 27005778 | 0.098 | 0.1752 | 0.5595 | 0.5758 | 0.1282 | 0.0375 | 0.9882 | 0.0111 | -0.009 | 0.0063 |
| Total lipids in small VLDL | 27005778 | 0.0389 | 0.153 | 0.254 | 0.7995 | 0.1553 | 0.0386 | 0.984 | 0.0089 | -0.0045 | 0.0061 |
| Total lipids in very large HDL | 27005778 | 0.0203 | 0.2188 | 0.0928 | 0.9261 | 0.0662 | 0.0302 | 10.147 | 0.0167 | -0.0039 | 0.0063 |
| Total lipids in very large VLDL | 27005778 | -0.1377 | 0.1414 | -0.9735 | 0.3303 | 0.1447 | 0.031 | 0.9708 | 0.0078 | 0.0023 | 0.0054 |
| Total lipids in very small VLDL | 27005778 | 0.0364 | 0.1695 | 0.2147 | 0.83 | 0.1183 | 0.0366 | 10.004 | 0.0124 | -0.0074 | 0.0062 |
| Townsend deprivation index at recruitment | 0 | -0.0161 | 0.0695 | -0.2311 | 0.8172 | 0.0344 | 0.0022 | 10.323 | 0.0093 | 0.0063 | 0.0059 |
| Transferrin | 25352340 | -0.0304 | 0.1383 | -0.2196 | 0.8262 | 0.167 | 0.0789 | 10.613 | 0.0287 | 0.0015 | 0.0068 |
| Transport type for commuting to job workplace: Car/motor vehi | 0 | 0.0179 | 0.116 | 0.1544 | 0.8773 | 0.0251 | 0.0033 | 10.221 | 0.0088 | -0.002 | 0.0062 |
| Transport type for commuting to job workplace: Cycle | 0 | 0.2486 | 0.1196 | 2.078 | 0.0377 | 0.0306 | 0.0035 | 10.174 | 0.0086 | -0.0145 | 0.0062 |
| Transport type for commuting to job workplace: Public transpor | 0 | -0.0188 | 0.1081 | -0.174 | 0.8619 | 0.0331 | 0.0038 | 10.189 | 0.0094 | 0.0029 | 0.0062 |
| Transport type for commuting to job workplace: Walk | 0 | -0.0253 | 0.1319 | -0.192 | 0.8477 | 0.0169 | 0.003 | 10.035 | 0.0079 | 0.0032 | 0.0059 |
| Treatment/medication code: adcal-d3 1.5g/10micrograms chewa | 0 | NA | NA | NA | NA | NA | NA | NA | NA | NA | NA |
| Treatment/medication code: alendronate sodium | 0 | NA | NA | NA | NA | NA | NA | NA | NA | NA | NA |
| Treatment/medication code: allopurinol | 0 | NA | NA | NA | NA | NA | NA | NA | NA | NA | NA |
| Treatment/medication code: amitriptyline | 0 | NA | NA | NA | NA | NA | NA | NA | NA | NA | NA |
| Treatment/medication code: amlodipine | 0 | NA | NA | NA | NA | NA | NA | NA | NA | NA | NA |
| Treatment/medication code: arcoxia 90mg tablet | 0 | NA | NA | NA | NA | NA | NA | NA | NA | NA | NA |
| Treatment/medication code: aspirin | 0 | NA | NA | NA | NA | NA | NA | NA | NA | NA | NA |
| Treatment/medication code: atenolol | 0 | NA | NA | NA | NA | NA | NA | NA | NA | NA | NA |
| Treatment/medication code: atorvastatin | 0 | NA | NA | NA | NA | NA | NA | NA | NA | NA | NA |
| Treatment/medication code: atrovent 20micrograms inhaler | 0 | NA | NA | NA | NA | NA | NA | NA | NA | NA | NA |
| Treatment/medication code: beconase 50micrograms nasal spray | 0 | NA | NA | NA | NA | NA | NA | NA | NA | NA | NA |
| Treatment/medication code: becotide 50 inhaler | 0 | NA | NA | NA | NA | NA | NA | NA | NA | NA | NA |
| Treatment/medication code: bendroflumethiazide | 0 | NA | NA | NA | NA | NA | NA | NA | NA | NA | NA |
| Treatment/medication code: bisoprolol | 0 | NA | NA | NA | NA | NA | NA | NA | NA | NA | NA |
| Treatment/medication code: candesartan cilexetil | 0 | NA | NA | NA | NA | NA | NA | NA | NA | NA | NA |
| Treatment/medication code: carbimazole | 0 | NA | NA | NA | NA | NA | NA | NA | NA | NA | NA |

| | | | | | | | | | | | |
|---|---|---|---|---|---|---|---|---|---|---|---|
| Treatment/medication code: cetirizine | 0 | NA | NA | NA | NA | NA | NA | NA | NA | NA | NA |
| Treatment/medication code: cipralex 5mg tablet | 0 | NA | NA | NA | NA | NA | NA | NA | NA | NA | NA |
| Treatment/medication code: citalopram | 0 | NA | NA | NA | NA | NA | NA | NA | NA | NA | NA |
| Treatment/medication code: clopidogrel | 0 | NA | NA | NA | NA | NA | NA | NA | NA | NA | NA |
| Treatment/medication code: co-amilofruse | 0 | NA | NA | NA | NA | NA | NA | NA | NA | NA | NA |
| Treatment/medication code: co-codamol | 0 | NA | NA | NA | NA | NA | NA | NA | NA | NA | NA |
| Treatment/medication code: codeine | 0 | NA | NA | NA | NA | NA | NA | NA | NA | NA | NA |
| Treatment/medication code: diclofenac | 0 | NA | NA | NA | NA | NA | NA | NA | NA | NA | NA |
| Treatment/medication code: dothiepin | 0 | NA | NA | NA | NA | NA | NA | NA | NA | NA | NA |
| Treatment/medication code: doxazosin | 0 | NA | NA | NA | NA | NA | NA | NA | NA | NA | NA |
| Treatment/medication code: enalapril | 0 | NA | NA | NA | NA | NA | NA | NA | NA | NA | NA |
| Treatment/medication code: eumovate cream | 0 | NA | NA | NA | NA | NA | NA | NA | NA | NA | NA |
| Treatment/medication code: evening primrose oil | 0 | NA | NA | NA | NA | NA | NA | NA | NA | NA | NA |
| Treatment/medication code: ezetimibe | 0 | NA | NA | NA | NA | NA | NA | NA | NA | NA | NA |
| Treatment/medication code: flax oil tablet | 0 | NA | NA | NA | NA | NA | NA | NA | NA | NA | NA |
| Treatment/medication code: flecainide | 0 | NA | NA | NA | NA | NA | NA | NA | NA | NA | NA |
| Treatment/medication code: furosemide | 0 | NA | NA | NA | NA | NA | NA | NA | NA | NA | NA |
| Treatment/medication code: gliclazide | 0 | NA | NA | NA | NA | NA | NA | NA | NA | NA | NA |
| Treatment/medication code: glucosamine product | 0 | NA | NA | NA | NA | NA | NA | NA | NA | NA | NA |
| Treatment/medication code: hydroxocobalamin product | 0 | NA | NA | NA | NA | NA | NA | NA | NA | NA | NA |
| Treatment/medication code: ibuprofen | 0 | NA | NA | NA | NA | NA | NA | NA | NA | NA | NA |
| Treatment/medication code: indivina 1mg/2.5mg tablet | 0 | NA | NA | NA | NA | NA | NA | NA | NA | NA | NA |
| Treatment/medication code: insulin product | 0 | NA | NA | NA | NA | NA | NA | NA | NA | NA | NA |
| Treatment/medication code: isosorbide dinitrate | 0 | NA | NA | NA | NA | NA | NA | NA | NA | NA | NA |
| Treatment/medication code: isosorbide mononitrate | 0 | NA | NA | NA | NA | NA | NA | NA | NA | NA | NA |
| Treatment/medication code: kapake tablet | 0 | NA | NA | NA | NA | NA | NA | NA | NA | NA | NA |
| Treatment/medication code: lansoprazole | 0 | NA | NA | NA | NA | NA | NA | NA | NA | NA | NA |
| Treatment/medication code: letrozole | 0 | NA | NA | NA | NA | NA | NA | NA | NA | NA | NA |
| Treatment/medication code: levothyroxine sodium | 0 | NA | NA | NA | NA | NA | NA | NA | NA | NA | NA |
| Treatment/medication code: liothyronine | 0 | NA | NA | NA | NA | NA | NA | NA | NA | NA | NA |
| Treatment/medication code: lisinopril | 0 | NA | NA | NA | NA | NA | NA | NA | NA | NA | NA |
| Treatment/medication code: lisinopril+hydrochlorothiazide 10m | 0 | NA | NA | NA | NA | NA | NA | NA | NA | NA | NA |
| Treatment/medication code: logynon tablet | 0 | NA | NA | NA | NA | NA | NA | NA | NA | NA | NA |
| Treatment/medication code: losartan | 0 | NA | NA | NA | NA | NA | NA | NA | NA | NA | NA |
| Treatment/medication code: metformin | 0 | NA | NA | NA | NA | NA | NA | NA | NA | NA | NA |
| Treatment/medication code: morphine | 0 | NA | NA | NA | NA | NA | NA | NA | NA | NA | NA |
| Treatment/medication code: nicorandil | 0 | NA | NA | NA | NA | NA | NA | NA | NA | NA | NA |
| Treatment/medication code: omacor 1g capsule | 0 | NA | NA | NA | NA | NA | NA | NA | NA | NA | NA |
| Treatment/medication code: omeprazole | 0 | NA | NA | NA | NA | NA | NA | NA | NA | NA | NA |
| Treatment/medication code: paracetamol | 0 | NA | NA | NA | NA | NA | NA | NA | NA | NA | NA |
| Treatment/medication code: perindopril | 0 | NA | NA | NA | NA | NA | NA | NA | NA | NA | NA |
| Treatment/medication code: prednisolone | 0 | NA | NA | NA | NA | NA | NA | NA | NA | NA | NA |
| Treatment/medication code: quinine | 0 | NA | NA | NA | NA | NA | NA | NA | NA | NA | NA |
| Treatment/medication code: ramipril | 0 | NA | NA | NA | NA | NA | NA | NA | NA | NA | NA |
| Treatment/medication code: ranitidine | 0 | NA | NA | NA | NA | NA | NA | NA | NA | NA | NA |
| Treatment/medication code: rhinocort 50micrograms nasal spray | 0 | NA | NA | NA | NA | NA | NA | NA | NA | NA | NA |
| Treatment/medication code: rosiglitazone | 0 | NA | NA | NA | NA | NA | NA | NA | NA | NA | NA |

| | | | | | | | | | | | |
|---|---|---|---|---|---|---|---|---|---|---|---|
| Treatment/medication code: salbutamol | 0 | NA | NA | NA | NA | NA | NA | NA | NA | NA | NA |
| Treatment/medication code: salmeterol product | 0 | NA | NA | NA | NA | NA | NA | NA | NA | NA | NA |
| Treatment/medication code: senna | 0 | NA | NA | NA | NA | NA | NA | NA | NA | NA | NA |
| Treatment/medication code: seretide 50 evohaler | 0 | NA | NA | NA | NA | NA | NA | NA | NA | NA | NA |
| Treatment/medication code: simvastatin | 0 | NA | NA | NA | NA | NA | NA | NA | NA | NA | NA |
| Treatment/medication code: spiriva 18micrograms inhalation ca | 0 | NA | NA | NA | NA | NA | NA | NA | NA | NA | NA |
| Treatment/medication code: symbicort 100/6 turbohaler | 0 | NA | NA | NA | NA | NA | NA | NA | NA | NA | NA |
| Treatment/medication code: testogel 50mg gel 5g sachet | 0 | NA | NA | NA | NA | NA | NA | NA | NA | NA | NA |
| Treatment/medication code: thyroxine product | 0 | NA | NA | NA | NA | NA | NA | NA | NA | NA | NA |
| Treatment/medication code: tramadol | 0 | NA | NA | NA | NA | NA | NA | NA | NA | NA | NA |
| Treatment/medication code: tranexamic acid | 0 | NA | NA | NA | NA | NA | NA | NA | NA | NA | NA |
| Treatment/medication code: venlafaxine | 0 | NA | NA | NA | NA | NA | NA | NA | NA | NA | NA |
| Treatment/medication code: ventolin 100micrograms inhaler | 0 | NA | NA | NA | NA | NA | NA | NA | NA | NA | NA |
| Treatment/medication code: vitamin b12 preparation | 0 | NA | NA | NA | NA | NA | NA | NA | NA | NA | NA |
| Treatment/medication code: warfarin | 0 | NA | NA | NA | NA | NA | NA | NA | NA | NA | NA |
| Treatment/medication code: xalatan 0.005% eye drops | 0 | NA | NA | NA | NA | NA | NA | NA | NA | NA | NA |
| Triglycerides | 20686565 | 0.0002 | 0.0811 | 0.0023 | 0.9982 | 0.1691 | 0.0295 | 0.9491 | 0.0189 | -0.002 | 0.0066 |
| Triglycerides in chylomicrons and largest VLDL particles | 27005778 | -0.2308 | 0.1551 | -14.878 | 0.1368 | 0.097 | 0.0301 | 0.9862 | 0.0081 | 0.0057 | 0.0052 |
| Triglycerides in IDL | 27005778 | 0.0107 | 0.1666 | 0.0641 | 0.9489 | 0.1221 | 0.0342 | 0.9989 | 0.0145 | -0.0073 | 0.0061 |
| Triglycerides in large VLDL | 27005778 | -0.091 | 0.1319 | -0.6902 | 0.4901 | 0.1231 | 0.0313 | 0.9783 | 0.0078 | 0.0056 | 0.0049 |
| Triglycerides in medium VLDL | 27005778 | -0.0222 | 0.1464 | -0.1517 | 0.8794 | 0.1063 | 0.0328 | 0.9898 | 0.0077 | 0.0018 | 0.005 |
| Triglycerides in small HDL | 27005778 | -0.1062 | 0.1806 | -0.588 | 0.5565 | 0.0792 | 0.0265 | 0.991 | 0.0083 | -0.0004 | 0.006 |
| Triglycerides in small VLDL | 27005778 | -0.0057 | 0.1485 | -0.0385 | 0.9693 | 0.1362 | 0.0374 | 0.9838 | 0.0081 | -0.0029 | 0.0057 |
| Triglycerides in very large HDL | 27005778 | -0.0864 | 0.1906 | -0.4533 | 0.6503 | 0.0791 | 0.0301 | 10.105 | 0.0302 | -0.0034 | 0.0061 |
| Triglycerides in very large VLDL | 27005778 | -0.1884 | 0.1439 | -13.098 | 0.1903 | 0.1283 | 0.0312 | 0.973 | 0.0078 | 0.0043 | 0.0055 |
| Triglycerides in very small VLDL | 27005778 | 0.0178 | 0.1475 | 0.1206 | 0.904 | 0.1557 | 0.0372 | 0.9873 | 0.0099 | -0.0062 | 0.0061 |
| Trunk fat mass | 0 | 0.0324 | 0.0483 | 0.672 | 0.5016 | 0.2335 | 0.0083 | 10.435 | 0.0215 | 0.0017 | 0.0082 |
| Trunk fat percentage | 0 | 0.0084 | 0.0504 | 0.166 | 0.8681 | 0.2077 | 0.0075 | 10.562 | 0.0202 | 1,95E-01 | 0.0086 |
| Trunk fat-free mass | 0 | 0.0701 | 0.0495 | 14.163 | 0.1567 | 0.2885 | 0.0144 | 10.666 | 0.0397 | 0.0068 | 0.0085 |
| Trunk predicted mass | 0 | 0.0712 | 0.0496 | 14.341 | 0.1515 | 0.287 | 0.0143 | 10.666 | 0.0395 | 0.0065 | 0.0085 |
| Type 2 Diabetes | 22885922 | -0.0247 | 0.1006 | -0.2457 | 0.8059 | 0.0828 | 0.0093 | 10.114 | 0.0088 | -0.0058 | 0.0053 |
| Type of tobacco previously smoked: Cigars or pipes | 0 | 0.5365 | 0.2934 | 18.285 | 0.0675 | 0.0102 | 0.0055 | 10.024 | 0.0073 | -0.0122 | 0.0059 |
| Types of physical activity in last 4 weeks: Heavy DIY (eg: weed | 0 | -0.0914 | 0.076 | -1.202 | 0.2294 | 0.0347 | 0.002 | 10.058 | 0.0093 | 0.0089 | 0.0056 |
| Types of physical activity in last 4 weeks: Light DIY (eg: pruni | 0 | -0.0547 | 0.0738 | -0.7418 | 0.4582 | 0.0385 | 0.0023 | 10.063 | 0.0098 | 0.0058 | 0.0062 |
| Types of physical activity in last 4 weeks: None of the above | 0 | -0.1666 | 0.085 | -19.597 | 0.05 | 0.0239 | 0.0019 | 10.146 | 0.0084 | 0.0094 | 0.0057 |
| Types of physical activity in last 4 weeks: Other exercises (eg: s | 0 | 0.0635 | 0.0729 | 0.8715 | 0.3835 | 0.0453 | 0.0028 | 10.175 | 0.0113 | -0.0056 | 0.0062 |
| Types of physical activity in last 4 weeks: Strenuous sports | 0 | 0.2106 | 0.1025 | 20.557 | 0.0398 | 0.0208 | 0.0019 | 10.181 | 0.0087 | -0.0126 | 0.0058 |
| Types of physical activity in last 4 weeks: Walking for pleasure | 0 | 0.1122 | 0.0738 | 15.209 | 0.1283 | 0.0365 | 0.0023 | 10.419 | 0.0096 | -0.0006 | 0.0061 |
| Types of transport used (excluding work): Car/motor vehicle | 0 | -0.1305 | 0.0895 | -14.582 | 0.1448 | 0.0259 | 0.002 | 10.035 | 0.0086 | 0.0094 | 0.0065 |
| Types of transport used (excluding work): Cycle | 0 | 0.0504 | 0.0923 | 0.5464 | 0.5848 | 0.0251 | 0.0018 | 10.228 | 0.0083 | -0.003 | 0.0062 |
| Types of transport used (excluding work): Public transport | 0 | -0.0664 | 0.0967 | -0.6868 | 0.4922 | 0.0223 | 0.0019 | 10.279 | 0.0079 | 0.0013 | 0.006 |
| Types of transport used (excluding work): Walk | 0 | -0.0162 | 0.0779 | -0.2073 | 0.8358 | 0.0345 | 0.0026 | 10.239 | 0.0109 | 0.0075 | 0.0065 |
| Tyrosine | 27005778 | -0.1587 | 0.1571 | -10.101 | 0.3125 | 0.083 | 0.0313 | 0.9962 | 0.0085 | 0.0038 | 0.0052 |
| Ulcerative colitis | 26192919 | 0.0831 | 0.0953 | 0.8718 | 0.3833 | 0.2426 | 0.0321 | 10.462 | 0.0123 | 0.0571 | 0.0061 |
| Underlying (primary) cause of death: ICD10: E85.4 Organ-limit | 0 | NA | NA | NA | NA | NA | NA | NA | NA | NA | NA |
| Underlying (primary) cause of death: ICD10: J84.1 Other inters | 0 | -0.1752 | 0.2539 | -0.6901 | 0.4901 | 0.1153 | 0.0707 | 0.9922 | 0.0078 | 0.0056 | 0.0061 |
| Urate | 23263486 | -0.0407 | 0.0697 | -0.5844 | 0.559 | 0.1773 | 0.0611 | 0.9341 | 0.0596 | 0.0048 | 0.0067 |

| | | | | | | | | | | | |
|---|---|---|---|---|---|---|---|---|---|---|---|
| Urinary albumin-to-creatinine ratio | 26631737 | -0.2327 | 0.1592 | -14.621 | 0.1437 | 0.0466 | 0.0091 | 0.9914 | 0.0075 | 0.0033 | 0.0056 |
| Urinary albumin-to-creatinine ratio (non-diabetes) | 26631737 | -0.2421 | 0.1519 | -1.594 | 0.1109 | 0.0572 | 0.0107 | 0.9891 | 0.0075 | 0.0035 | 0.0058 |
| Used an inhaler for chest within last hour | 0 | -0.3048 | 0.2139 | -14.249 | 0.1542 | 0.0045 | 0.0017 | 10.012 | 0.0079 | 0.006 | 0.0061 |
| Usual walking pace | 0 | 0.1572 | 0.0621 | 25.323 | 0.0113 | 0.0778 | 0.0032 | 1.026 | 0.0131 | -0.0121 | 0.0075 |
| Valine | 27005778 | 0.0917 | 0.1796 | 0.511 | 0.6094 | 0.0794 | 0.0219 | 10.046 | 0.0087 | -0.0006 | 0.0059 |
| Vascular/heart problems diagnosed by doctor: Angina | 0 | -0.0512 | 0.0983 | -0.5211 | 0.6023 | 0.0209 | 0.0021 | 10.306 | 0.0091 | 0.0025 | 0.006 |
| Vascular/heart problems diagnosed by doctor: Heart attack | 0 | -0.0402 | 0.088 | -0.4567 | 0.6479 | 0.0191 | 0.0019 | 10.071 | 0.0086 | -0.0013 | 0.0056 |
| Vascular/heart problems diagnosed by doctor: High blood press | 0 | -0.0632 | 0.0532 | -11.875 | 0.235 | 0.12 | 0.0054 | 1.022 | 0.0173 | 0.0083 | 0.0081 |
| Vascular/heart problems diagnosed by doctor: None of the abov | 0 | 0.0422 | 0.0545 | 0.7744 | 0.4387 | 0.1164 | 0.0052 | 10.242 | 0.0167 | -0.0043 | 0.0081 |
| Vitamin and mineral supplements: Multivitamins +/- minerals | 0 | 0.0117 | 0.0909 | 0.1286 | 0.8977 | 0.0238 | 0.002 | 10.103 | 0.0092 | 0.0016 | 0.0058 |
| Vitamin and mineral supplements: None of the above | 0 | -0.0056 | 0.0806 | -0.0698 | 0.9444 | 0.0296 | 0.002 | 10.001 | 0.0083 | -0.0025 | 0.0058 |
| Vitamin and mineral supplements: Vitamin A | 0 | -0.1985 | 0.1927 | -10.298 | 0.3031 | 0.004 | 0.0015 | 10.006 | 0.0077 | 0.0018 | 0.006 |
| Vitamin and mineral supplements: Vitamin B | 0 | -0.1244 | 0.152 | -0.8189 | 0.4129 | 0.0076 | 0.0016 | 0.9989 | 0.0075 | -0.0064 | 0.0059 |
| Vitamin and mineral supplements: Vitamin C | 0 | -0.0631 | 0.0978 | -0.6453 | 0.5187 | 0.0137 | 0.0018 | 0.9967 | 0.0085 | 0.0057 | 0.0059 |
| Vitamin and mineral supplements: Vitamin D | 0 | 0.001 | 0.1507 | 0.0069 | 0.9945 | 0.006 | 0.0017 | 0.9976 | 0.0081 | 0.002 | 0.0059 |
| Vitamin and mineral supplements: Vitamin E | 0 | 0.0434 | 0.1403 | 0.3096 | 0.7569 | 0.007 | 0.0014 | 0.9941 | 0.0082 | -0.0031 | 0.0053 |
| Waist circumference | 25673412 | 0.0854 | 0.0615 | 1.388 | 0.1651 | 0.1244 | 0.0053 | 0.8331 | 0.0091 | -0.0047 | 0.0054 |
| Waist circumference | 0 | 0.004 | 0.0487 | 0.0818 | 0.9348 | 0.2015 | 0.0078 | 10.399 | 0.0203 | 0.0016 | 0.0079 |
| Waist-to-hip ratio | 25673412 | -0.0138 | 0.0663 | -0.2089 | 0.8345 | 0.1162 | 0.0076 | 0.9073 | 0.0105 | 0.0009 | 0.0058 |
| Wears glasses or contact lenses | 0 | 0.006 | 0.0971 | 0.0614 | 0.9511 | 0.0163 | 0.0017 | 10.016 | 0.0089 | 0.0008 | 0.0055 |
| Weight | 0 | 0.0476 | 0.0479 | 0.9929 | 0.3207 | 0.2676 | 0.0106 | 10.394 | 0.0266 | 0.0042 | 0.008 |
| Weight change compared with 1 year ago | 0 | -0.0411 | 0.1394 | -0.2946 | 0.7683 | 0.0078 | 0.0016 | 10.289 | 0.0075 | 0.0037 | 0.0057 |
| Wheeze or whistling in the chest in last year | 0 | -0.0679 | 0.0623 | -10.894 | 0.276 | 0.0619 | 0.0036 | 10.146 | 0.0127 | 0.0034 | 0.0069 |
| Whole body fat mass | 0 | 0.0207 | 0.0485 | 0.4276 | 0.669 | 0.2374 | 0.0084 | 10.392 | 0.0213 | 0.0019 | 0.0082 |
| Whole body fat-free mass | 0 | 0.0697 | 0.0489 | 14.237 | 0.1545 | 0.2916 | 0.0136 | 10.596 | 0.0372 | 0.0065 | 0.0083 |
| Whole body water mass | 0 | 0.07 | 0.0492 | 14.219 | 0.1551 | 0.2914 | 0.0136 | 10.607 | 0.037 | 0.0064 | 0.0083 |
| Why reduced smoking: Illness or ill health | 0 | -0.0855 | 0.2206 | -0.3875 | 0.6984 | 0.0948 | 0.0495 | 10.032 | 0.0075 | 0.0044 | 0.0057 |
| Why stopped smoking: Doctors advice | 0 | -0.0464 | 0.161 | -0.2883 | 0.7731 | 0.0206 | 0.0059 | 0.9932 | 0.0075 | -0.003 | 0.0054 |
| Why stopped smoking: Health precaution | 0 | 0.0414 | 0.1191 | 0.3474 | 0.7283 | 0.0567 | 0.0072 | 0.994 | 0.0085 | -0.0047 | 0.0067 |
| Why stopped smoking: None of the above | 0 | -0.0295 | 0.1303 | -0.2262 | 0.8211 | 0.0427 | 0.0062 | 0.995 | 0.0077 | 4,18E-01 | 0.0063 |
| Work/job satisfaction | 0 | -0.1182 | 0.136 | -0.8687 | 0.385 | 0.0491 | 0.0074 | 0.9968 | 0.0084 | 0.008 | 0.0058 |
| Worrier / anxious feelings | 0 | -0.0156 | 0.0579 | -0.2697 | 0.7874 | 0.0797 | 0.0051 | 10.009 | 0.0155 | 0.006 | 0.0063 |
| Worry too long after embarrassment | 0 | 0.019 | 0.0621 | 0.3054 | 0.7601 | 0.066 | 0.0036 | 10.128 | 0.0115 | 0.0066 | 0.0065 |
| Years of schooling (proxy cognitive performance) | 25201988 | -0.0453 | 0.0844 | -0.5368 | 0.5914 | 0.1057 | 0.0076 | 10.281 | 0.0111 | 0.0012 | 0.0063 |
| Years of schooling 2013 | 23722424 | -0.0688 | 0.0834 | -0.8254 | 0.4092 | 0.0831 | 0.0064 | 10.206 | 0.0104 | 0.0047 | 0.006 |
| Years of schooling 2016 | 27225129 | 0.0738 | 0.0518 | 14.245 | 0.1543 | 0.126 | 0.005 | 0.9341 | 0.0131 | -0.0096 | 0.0068 |

Source: number denotes PMID; UKBB, UK Biobank; rg, genetic correlation (regression); se, standard error; FDR, false discovery rate adjusted p-value; h2 obs, shared heritability observed; h2 int, sha
