## Supplementary figures and images for "Genome-wide estimates of heritability and genetic correlations in Essential Tremor"

### Supplementary Figure 1

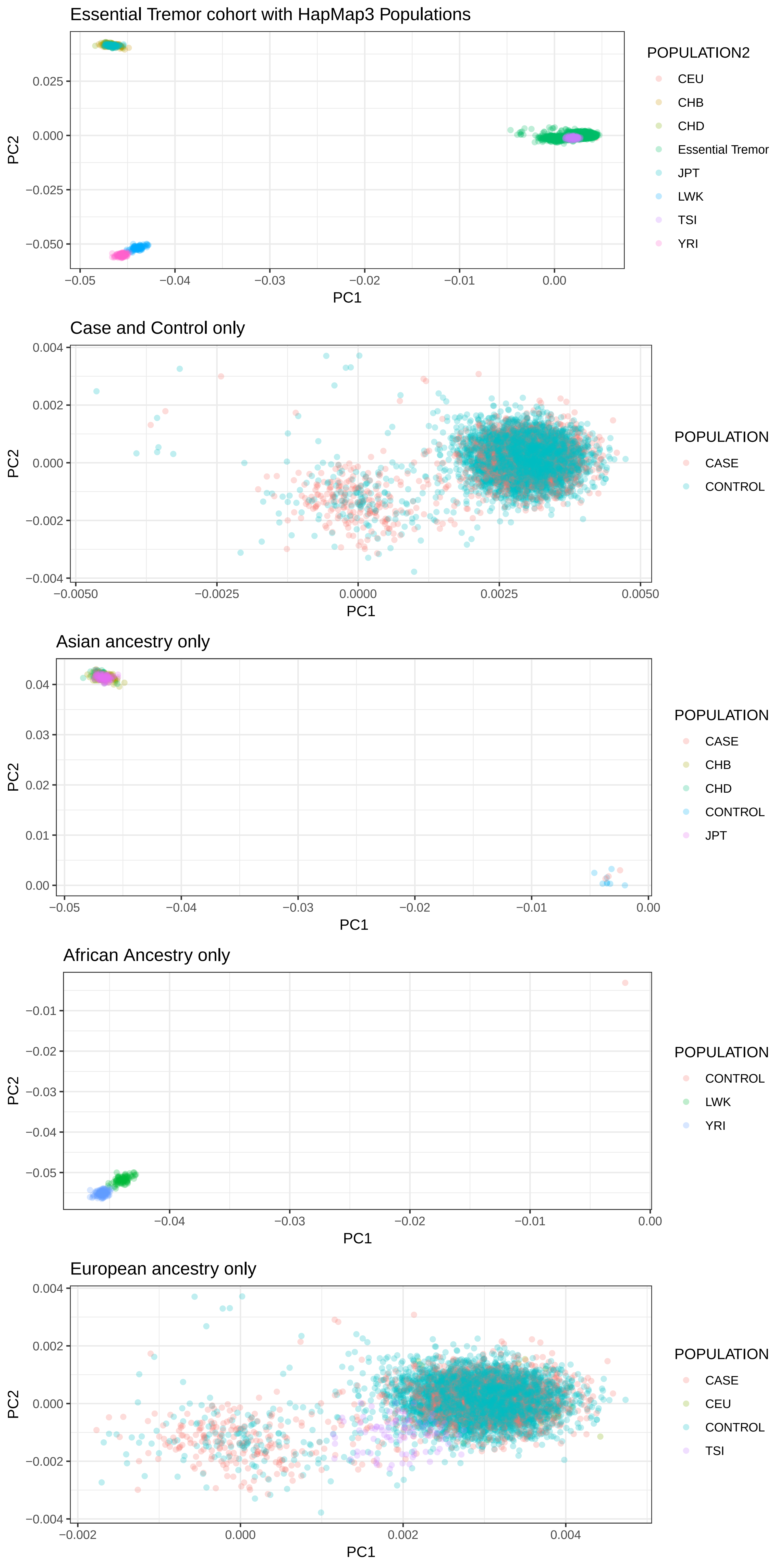
